# Spatial and single-cell expression analyses reveal complex expression domains in early wheat spike development

**DOI:** 10.1101/2025.02.15.638402

**Authors:** Xiaosa Xu, Huiqiong Lin, Junli Zhang, German Burguener, Francine Paraiso, Kun Li, Connor Tumelty, Chengxia Li, Yuchen Liu, Jorge Dubcovsky

## Abstract

Wheat is important for global food security and understanding the molecular mechanisms driving spike and spikelet development can inform the development of more productive varieties. In this study, we integrated single-molecule fluorescence *in situ* hybridization (smFISH) and single-cell RNA sequencing (scRNA-seq) to generate an atlas of cell clusters and expression domains during the early stages of wheat spike development. We characterized spatiotemporal expression of 99 genes by smFISH in 48,225 cells at the early transition, late double ridge, and floret primordia stages. These cells were grouped into 21 different expression domains, including four in the basal region of the developing spikelets and three different meristematic regions, which were consistent across spikelets and sections. Using induced mutants, we revealed functional roles associated with the specific expression patterns of *LFY* in intercalary meristems, *SPL14* in inflorescence meristems, and *FZP* in glume axillae. Complementary scRNA-seq profiling of 26,009 cells from W2.5 and W3.5 stages identified 23 distinct cell clusters. We used the scRNA-seq information to impute the expression of 74,464 genes into the spatially anchored smFISH-labelled cells and generated a public website to visualize them. We used experimental and imputed expression profiles, together with co-expression studies and correlation matrices, to annotate the scRNA-seq clusters. From co-expression analyses, we uncovered genes associated with boundary genes *TCP24* and *FZP*, as well as the meristematic genes *AGL6* and *ULT1.* The smFISH and scRNA-seq studies provided complementary tools for dissecting gene networks that regulate spike development and identifying new co-expressed genes for functional characterization.

## BACKGROUND

In 2023, the world produced approximately 799 million tons of wheat grain [1], yet further increases are necessary to feed a growing human population. Wheat grains develop within inflorescences called spikes, thus improving spike productivity is an important breeding objective. Traditional and molecular breeding methods have made small incremental advances in spike productivity, but a deeper understanding of the regulatory mechanisms underlying spike development could accelerate the engineering of higher-yielding wheat varieties.

Wheat spike development starts with the transition of a shoot apical meristem (SAM) that produces leaves to an inflorescence meristem (IM) that produces spikelets. During the vegetative phase, the SAM produces lateral meristems that develop into leaves (Fig.S1A), whereas the subtended axillary meristems generally fail to grow out immediately [2]. By contrast, the wheat IM produces lateral meristems (Fig.S1B), that later differentiate into two distinct ridges (the double ridge stage, Fig. S1C). The lower ridges (also known as leaf ridges) are suppressed, whereas the upper ridges rapidly develop into spikelet meristems (SMs) that generate sessile spikelets attached directly to the inflorescence axis (rachis. Fig. S1D). Therefore, the transition from the vegetative to the reproductive phase involves both a drastic shift in the relative developmental rates of bract and axillary meristems and a change in the identity of the axillary meristems.

Wheat MADS-box genes from the *SQUAMOSA* clade (*VRN1*, *FUL2*, and *FUL3*) play an essential role in the repression of the lower ridge. Wheat plants with simultaneous loss-of-function mutations in these three genes show a normal transition of the SAM to the double ridge stage and stem elongation, but the lower ridges are derepressed and develop into leaves or bracts [3]. Bract outgrowth has also been reported in wheat plants with loss-of-function mutations in the *SQUAMOSA PROMOTER BINDING PROTEIN LIKE17* (*SPL17*) and *SPL14*, two regulators of the *SQUAMOSA* genes [4]. The maize orthologs of these genes (*TSH4*, *UB2*, and *UB3*) play a similar role [5, 6], suggesting a conserved function across grass species.

The wheat *SQUAMOSA* genes also play a critical role in the determination of the meristem identity of the upper ridge. In the absence of these genes, the upper ridges develop into vegetative meristems rather than SMs, resulting in a “spike of tillers” [3]. Taken together, these results indicate that the *SQUAMOSA* genes are essential for conferring SM identity to the axillary meristem and suppressing bract development, but not for the initial transition of the SAM to the double ridge stage and the subsequent stem elongation [3].

The MADS-box genes from the *SVP* clade (*VRT2, SVP1*, *SVP3*) play similar roles as the *SQUAMOSA* genes during the initial stages of spike development. Individual loss-of-function mutations in *VRN1* or *FUL2* [3], or simultaneous mutations in *VRT2* and *SVP1* [7], both result in delayed heading, shorter stems and increased spikelet number per spike (SNS). However, at later stages of spike development, the SQUAMOSA proteins transcriptionally repress the *SVP-*clade genes to facilitate interactions with SEPALLATA1 proteins, which are required for normal spikelet development [7, 8].

The normal development of the wheat lateral spikelets begins with the formation of two sterile bracts called glumes, followed by slightly different bracts called lemmas that subtend the floral organs. Differences between glumes and lemmas are regulated by the *APETALA2-like* genes *AP2L5* (*Q* gene) and *AP2L2*, which are cleaved by the microRNA miR172. Mutations in the miR172 binding sites of these genes result in increased transcript levels and conversions of sterile glumes into fertile lemmas [9, 10]. Conversely, loss-of-function mutation in *AP2L5* and *AP2L2* result in conversions of fertile lemmas into sterile glumes [11]. These results indicate an important role of *AP2L5* and *AP2L2* in the regulation of glume and lemma organ identities and the development of the axillary floret meristem (FM).

In grasses, the first whorl of floral organs generated by the FM is the palea, which is regulated by the MADS-box gene *AGL6* [12]. Development of the internal floral whorls (lodicules, stamens, and pistils) is severely disrupted in wheat plants with loss-of-function mutations in *LEAFY* (*LFY*) or *WHEAT ORTHOLOGUE of APO1* (*WAPO1*), two genes that are expressed in partially overlapping domains at the FM and encode interacting proteins [13, 14]. In addition to the disruptions in floral development, loss-of-function mutations in either or both genes significantly reduce the rate at which the IM generates lateral meristems, leading to drastic reductions in SNS [13]. Natural alleles with increased SNS have been identified for both *WAPO1* [14–17] and *LFY* [18], and allele combinations that maximize SNS have been also reported [18].

Despite progress in the genetic characterization of wheat spike development, the diversity of expression domains within the wheat spike and their associated gene networks remain poorly understood. In this study, we characterized the spatial expression of 99 wheat genes during the initial SAM-IM transition, the late double ridge and floret primordia stages using single-molecule fluorescence *in situ* hybridization (smFISH). To expand the list of characterized genes, we integrated the smFISH data with single-cell RNA sequencing (scRNA-seq) from protoplasts isolated during the late double ridge and floret primordia stages. These complementary datasets provided complementary resources to study the gene networks regulating spike development and revealed opportunities to engineer more productive wheat spikes.

## RESULTS AND DISCUSSION

### 1. Spatial expression and cell segmentation

To identify the distinct cell clusters and expression domains involved in early wheat spike development, we analyzed the expression of 99 genes simultaneously (Data S1) by smFISH at three early stages of spike development (W1.5, W2.5, and W3.5 as defined by the Waddington scale [19]). W1.5 marks the initial transition of the shoot apical meristem (SAM) to the reproductive stage (Fig. 1A). W2.5 is defined as the late double ridge stage before glume primordia initiation (Fig. 1B and S1A), and W3.5 as the floret primordia stage (Fig. 1C and S1B). At W3.5, the inflorescence meristem (IM) begins its transition to a terminal spikelet (IM→TS), and the most advanced spikelet meristems (SMs) at the center of the spike initiate floret meristems (FMs, Figs. 1C and S1B).

**Fig. 1.**
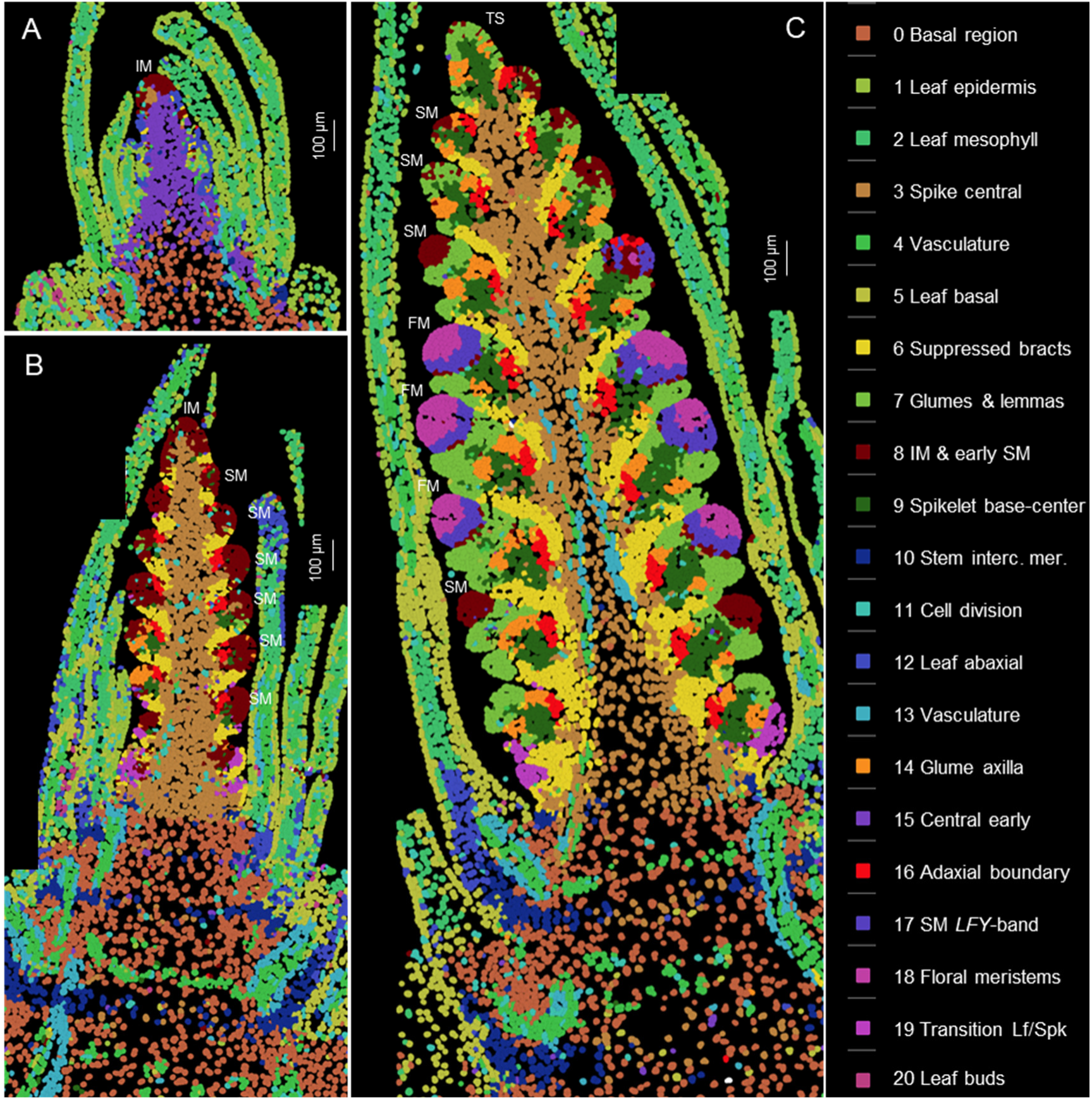
Cell clusters based on the expression of 99 genes analyzed by smFISH at three stages of wheat spike development. Cell clusters were generated from the three sections shown in this figure and replicate sections presented in Fig. S2. Identical colors represent the same clusters across the six sections. **A** W1.5: initial transition from the vegetative to the reproductive stage (section C1-1). **B** W2.5: late double ridge stage (section B1-3). **C** W3.5: floret primordia stage (section A1-2). Scale bars = 100 μm. A detailed description of the cell clusters, cell counts, and preferentially expressed genes in each cluster is available in Data S2. IM= inflorescence meristem, SM= spikelet meristem, FM= floral meristem, TS= terminal spikelet.

We analyzed two sections from each developmental stage (Figs. 1, S2) using the cell segmentation program QuPath [20] and identified a total of 48,225 cells. In the same sections, cell walls were visualized with calcofluor-white (Figs. S3, S4). Based on the expression of 99 genes (Data S1), these cells were grouped into 21 clusters, represented by consistent colors across all three developmental stages and replications (Figs. 1, S2). The individual cell clusters are presented separately in Fig. S5 and can be also visualized, together with heatmaps of the 99 genes, at https://dubcovskylab.ucdavis.edu/JD99-wheat-spike-smFISH.

We characterized 7,624 cells at W1.5, 16,400 at W2.5, and 24,210 at W3.5 (Fig. 1 and S2), showing a clear increase in cell number during spike development. These cells also exhibited increasing differentiation, with 14 clusters at W1.5, 19 at W2.5, and 21 at W3.5 (Fig. 1 and S2, Data S2). At the two later stages, cell clusters formed at the SMs and beneath them were consistent across spikelets within the same spike and between sections in Figs. 1 and S2, demonstrating good reproducibility. The gene expression profiles obtained with Molecular Cartography also showed good reproducibility with previous *in situ* hybridization results (Fig. S6) and recently published wheat spatial transcriptomics results generated using MERFISH [21] (Fig. S7).

Annotation of the cell clusters and their preferentially expressed genes are summarized in Data S2, whereas the correlations among cell clusters are presented in Data S3. The five cell clusters detected in the leaves (c1, c2, c5, c12, and c20) were highly correlated (*R* ≥ 0.78, Data S3) and are not discussed further in this study. To organize the description of these clusters and their changes during spike development, we separated the developing spike into five regions: the basal region below the spike, the spike central region, the region below the SMs, the SMs, and IM.

#### 1.1. The basal region below the spike

The basal region below the spike consists of cells that will later differentiate into nodes and internodes, forming the stem and contributing to important agronomic traits such as plant height and stem strength. This region is primarily composed of cells grouped in cluster c0, which are located below the epidermis and outside the vascular tissue and are referred hereafter as ground tissue. At W2.5 and W3.5 this basal region is interrupted by alternating bands of cells from clusters c4 (vasculature) and c10 (stem intercalary meristem, Figs. 1, S2, S5).

**Fig. 2.**
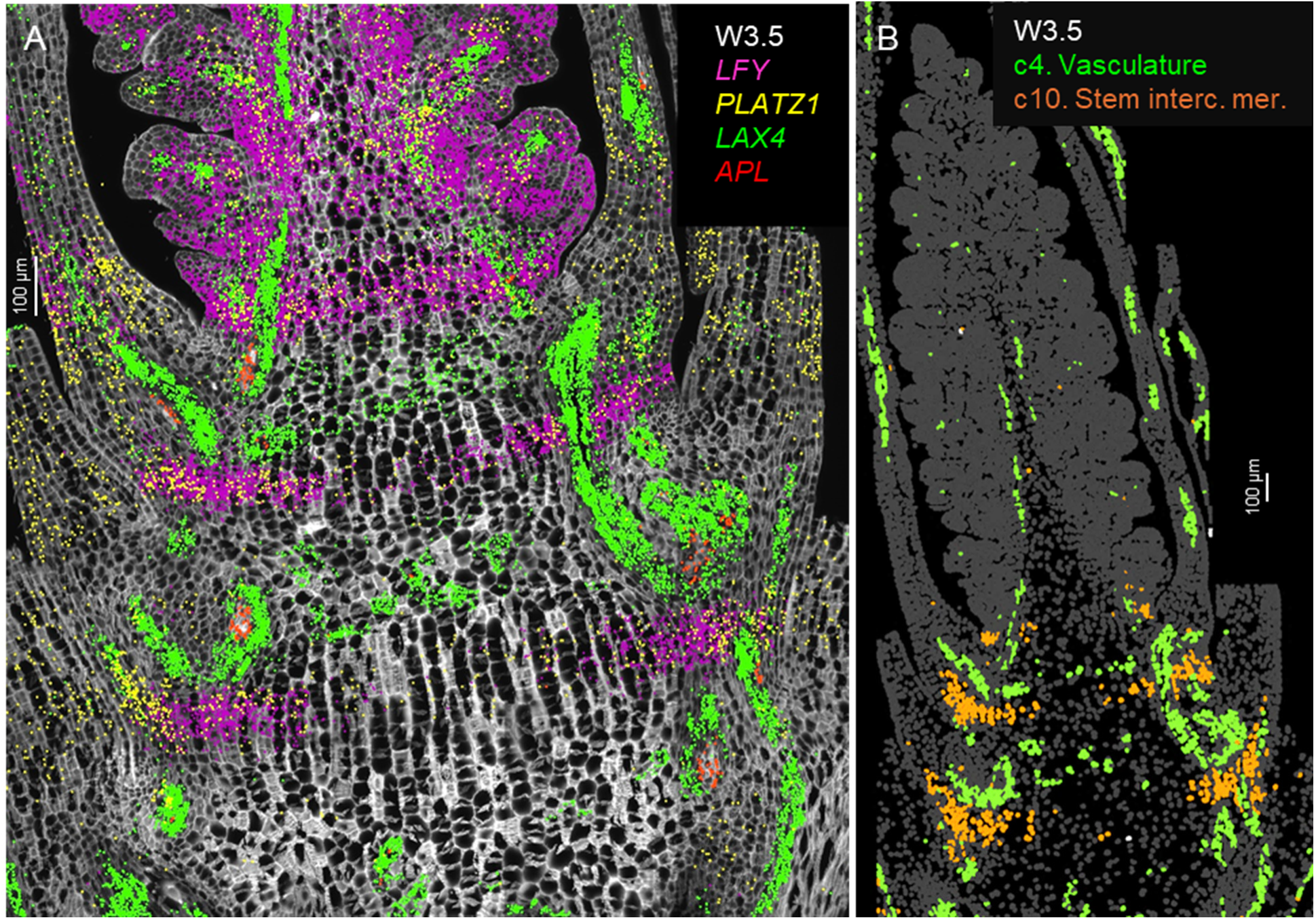
Single-molecule fluorescence *in-situ* hybridization (smFISH) characterization of the basal region below the spike at W3.5. Cell walls were stained with calcofluor-white. **A** Vasculature patterning genes *LAX4* (green) and *APL* (red) are concentrated in the region with shorter cells (likely future nodes). *LFY* (purple) and *PLATZ1* (yellow) mark a band in the region with longer cells (likely future internodes). **B** Cell segmentation with highlighted clusters c4 in green (vascular tissue) and c10 in orange (stem intercalary meristem). Scale bar= 100 μm.

In longitudinal sections stained with calcofluor-white at W2.5 and W3.5 (Figs. S3, S4), two alternating zones are visible at the basal region: one with shorter cells likely corresponding to the nodes, and the other with longer cells likely corresponding to the internodes of the future stem. Enriched expression of the procambial marker *LAX4* [22] and the phloem marker *APL* [23] was observed in the smaller cells, supporting their association with future nodes (Fig. 2A). These cells, together with those expressing the vascular genes in the leaf and spike vascular tissues were all included in cluster c4 (Fig. 2B). Based on the location of c4 cells in the nodes and the orientation of the vascular bundles we hypothesize that these cells may interconnect the parallel vascular bundles of the internodes.

Cells classified as cluster c10 (Fig. 2B) appeared as incomplete bands in the future internode regions at W2.5 and W3.5 (Figs. 1, S5). In the three-dimensional space, c10 cells may form a transversal disc across the stem located at the point where the abaxial base of the leaves joins the stem. These c10 cells were characterized by the differential expression of *LFY*, *PLATZ1* (Fig. 2A), *NAL1*, *ULT1*, and *SVP1* (Fig. S8, Data S2). Previous studies have demonstrated that loss-of-function mutations in wheat *PLATZ1* [24] or *SVP1* [7] significantly reduced plant height, and that overexpression of these genes partially restored normal plant height. The functional validation of the roles of *PLATZ1* and *SVP1* in stem development suggested that cluster c10 may correspond to the stem intercalary meristem, a zone at the base of grass internodes that contributes new cells to the elongating internodes [25]. This hypothesis was further supported by our finding that *LFY* also plays an important role in wheat stem development.

Using the *lfy*-null mutant developed in a previous study [13], we showed here that the mutant has no significant effect on stem length, but resulted in significantly wider nodes (22.3% ± 1.8%) and internodes (11.5% ± 2.0%). Moreover, the mutants produced more solid stems, showing a 62% ± 17% reduction of the diameter of the hollow central region relative to the WT (Fig. S9, Data S4). The *Short and Solid Culm* (*ssc*) rice mutant is a *LFY* allele with a D16N amino acid polymorphism and a 3-fold higher transcript levels than the wildtype, which has been associated with reduced stem length and diameter [26]. These results revealed a shared role of *LFY* on stem development between wheat and rice and are a good example of the value of the smFISH data to generate functional hypothesis.

Since the gibberellic acid (GA) hormone plays a critical role in stem elongation, we examined the spatial distribution of two genes involved in GA biosynthesis (*GA20ox1* and *GA20ox2*) and one involved in GA degradation (*GA2ox1*). At W2.5 and W3.5, the two GA biosynthetic genes were enriched in c0 at the basal region below the spike (Fig. S10B-C, Data S2). Loss-of-function mutations in either *GA20ox1* or *GA20ox2* reduced the levels of bioactive GA_1_ and GA_4_, resulting in shorter plants than the wildtype [27]. These findings support a positive role of these genes in wheat stem elongation, consistent with their spatial expression adjacent to the stem intercalary meristem region (c10).

The catabolic gene *GA2ox1* showed higher levels of expression in the central region of the developing spikes at W1.5 (cluster c15) than at later stages (cluster c3, Fig. S10A, Data S2). We speculate that the elevated *GA2ox1* expression at early spike development prevents premature cell elongation and induction of *SOC1* and *LFY*, which are associated with increased GA levels [28]. At W2.5 and W3.5, *GA2ox1* expression was restricted to the periphery of the region enriched in GA biosynthetic genes, possibly forming a ring of reduced GA abundance around the elongating cells. The negative role of *GA2ox1* in stem elongation has been functionally validated in rice plants by both over-expression and ectopic expression of *GA2ox1* under the *OsGA3ox2* promoter [29]. We hypothesize that the spatial distribution of GA metabolic and catabolic genes described in this study may play an important role in normal wheat stem elongation.

#### 1.2. The transition zone between leaf and spike

At W2.5 and W3.5, a group of cells within cluster c19 was detected in the transition region between leaves and spike (Fig. S11A-B). The genes preferentially expressed in cluster c19 include the *SPL* transcription factors *SPL2*, *SPL14*, and *SPL17* (Fig. S11C-D), the *TCP* transcription factor (*TEOSINTE BRANCHED1/CYCLOIDEA/PROLIFERATING CELL FACTOR*) *TB1* and *TB2* (Fig. S11E-F), and the MADS-box genes *FUL2* and *SOC1-1* (Data S2). Several of these genes have previously been associated with the regulation of spike architecture.

Mutations in the *SPL14* and *SPL17* genes result in the formation of ectopic bracts at the base of the inflorescence in rice [30], maize [5] and hexaploid wheat [4]. Based on the basal location of these bracts, we hypothesize that the preferential expression of *SPL14* and *SPL17* in cluster c19 may contribute to bract repression in the transition zone. *TSH4,* the maize ortholog of *SPL17* and its close paralogues *UB2* and UB3 (*SPL14*) work together to target genes responsible for axillary meristem suppression such as *TB1* and auxin response regulators [31].

In rice, *SPL14* is a direct regulator of *TB1* [32], which in our study was also preferentially expressed in cluster c19. *TB1* reduces lateral branching, and mutants of this gene in maize [33] and rice [34] show increased tillering. *TB1* mutants in barley and wheat also show increased tillering and produce additional lateral spikelets in barley [35] and paired spikelets in wheat [36, 37]. These findings indicate a conserved role of *TB1* in the regulation of branching.

*SQUAMOSA* genes are direct targets of the SPL proteins [38, 39] and combined loss-of-function mutations in the wheat *SQUAMOSA* genes *VRN1* and *FUL2* result in de-repression of the lower ridge, leading to bract outgrowth below the spikelets at the base of the spike [3]. Individual *vrn1* or *ful2* mutants show no bract outgrowth, demonstrating that *FUL2* and *VRN1* play redundant roles in bract repression.

Based on the physical location of the c19 cells and the function of the genes preferentially expressed in this region (Data S2), we hypothesize that the C19 cells contribute to the transition between the proximal region (where leaf meristems exhibit rapid development and axillary meristems undergo a temporary arrest) and the distal region (where the subtending bracts are suppressed and upper ridges rapidly develop into spikelet meristems).

#### 1.3. The central region of the spike

The ground tissue cells in the central region of developing spikes were primarily assigned to cluster c15 at W1.5 and cluster c3 at both W2.5 and W3.5 (Figs. 1, S5), which were highly correlated with each other (*R*= 0.70). These two clusters also showed strong correlations with the ground tissue at the adjacent basal region beneath the spike (cluster c0, Data S3). Cells from cluster c15 showed stronger correlations with the leaf mesophyll (*R=* 0.74) than cluster c3 (*R=* 0.46, Data S3), suggesting progressive differentiation of the ground tissue at the center of the spike. Several genes were preferentially expressed at the spike central region across all three developmental stages, including *AP3*, *bZIPC4* (Fig. S12A-C), *IDD4* and *OSH1* (synonymous with *KNOX3*, Fig. S12D-F). The homeobox gene *OSH1* was expressed at the base and center of the spikelets, but not in developing glumes and lemmas. This expression pattern resembles that of its orthologs in maize and rice [40].

In Fig. 3, we present the expression profiles of four vascular markers: the auxin influx carrier *LAX4* [41], the phloem marker *APL* [23], and the duplicated auxin efflux carriers *PIN1a* and *PIN1b* [22, 42] (Fig. 3). In *Brachypodium*, *PIN1b* plays a role in sink-finding and connecting new primordia to existing vasculature, whereas *PIN1a* forms narrower canals than *PIN1b* and is likely involved in vein patterning [42]. A similar pattern was observed in wheat, where *PIN1b* was expressed in a broader area than *PIN1a* (Fig. 3).

**Fig. 3.**
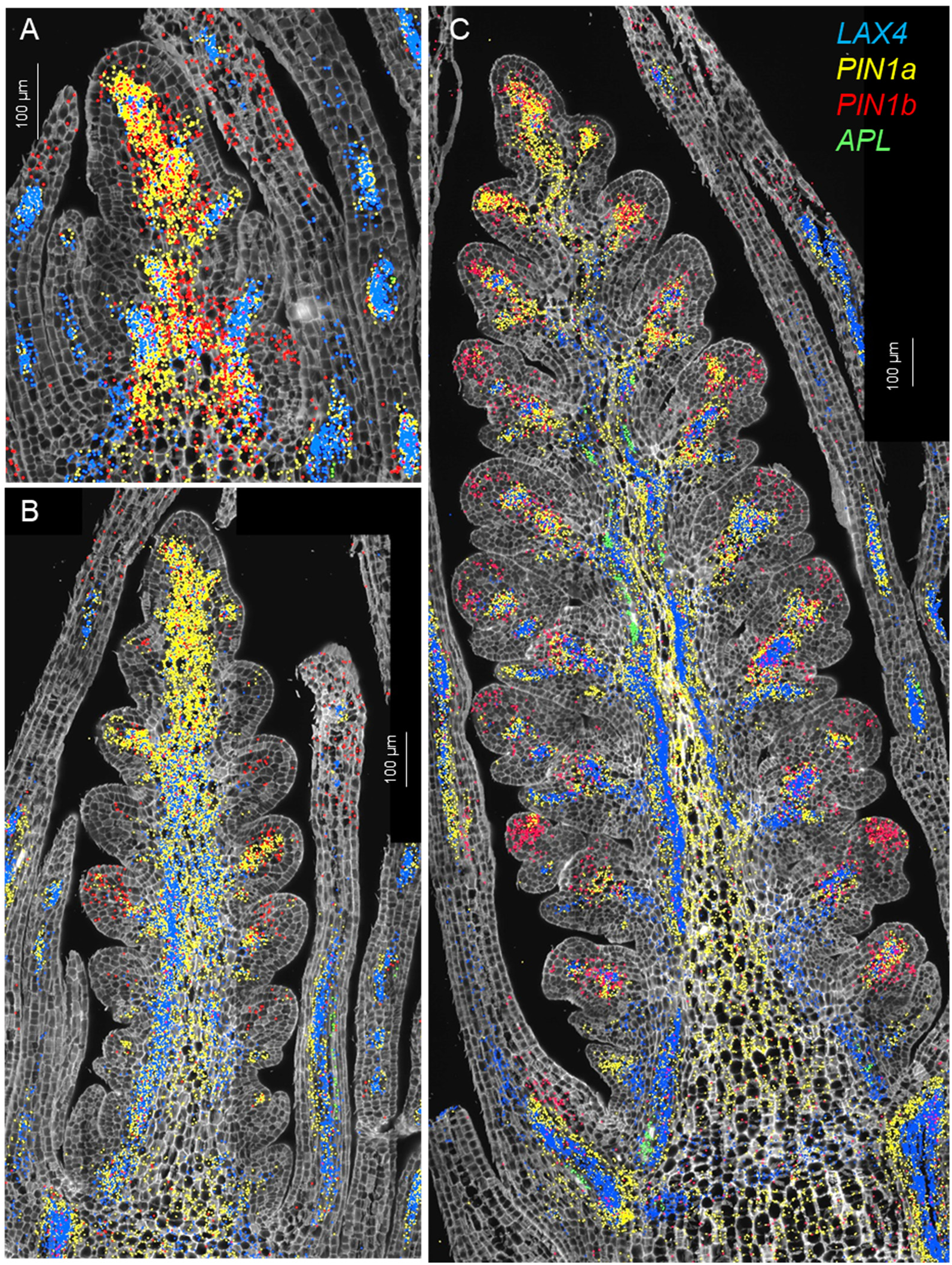
Single-molecule fluorescence *in-situ* hybridization characterization of the spike central region (cluster c3). Spike central region (cluster c3). **A** W1.5, **B** W2.5, **C** W3.5. In all three panels, *LAX4* hybridization signal is shown in blue, *APL* in green, *PIN1a* in yellow, and *PIN1b* in red. Cell walls are stained with calcofluor-white.

*PIN1a* and *PIN1b* were preferentially expressed in clusters c4 and c13 (Data S2, Fig. S13A), which were annotated as vasculature clusters. *APL* was among the preferentially expressed genes in both c4 and c13 (Data S2) but most of its expression was outside these clusters (Fig. S13B).

The expression of *CEN5* overlapped well with c13 (Fig. S13B) whereas the expression of *LAX4* overlapped better with c4 (Fig. S13C). *PIN1a* showed extensive overlap with both clusters (Fig. S13D). The early accumulation of *PIN1a* and *LAX4* (=*ZmLAX2*) in maize leaves marked the initial procambium specification [22], suggesting that c13 and c4 represent early stages of vascular development.

In maize, *LAX4* and *PIN1a* have antagonistic functions and are frequently co-expressed in pro-vascular cells [22, 41], an overlap that was also apparent in wheat spikes at all three developmental stages (Fig. 3). The phloem marker *APL* was detected at developing vascular tissue of leaves at W1.5 and W2.5 (Fig. 3A-B), and in the central region of the spike at W3.5 (Fig. 3A).

In summary, an important function of the c4 and c13 vascular cells in the central region of the spike is to provide an efficient vascular system that transports water and nutrients to the developing spikelets. The central c3 cells are green from early developmental stages, suggesting that they contribute photosynthetic products to the developing spikes (see section 2.5) and also provide structural support to flowers and grains.

#### 1.4. Region proximal to the spikelet meristem (SM)

At W2.5 (Fig. 4A), cortex cells proximal to the spikelet meristem were primarily classified into clusters c6 (suppressed bract), c9 (central SM basal region), and c16 (adaxial boundary). These clusters remained evident at W3.5, with the addition of clusters c7 (glume and lemma) and c14 (glume axilla). A few cells from clusters c7 and c14 were also detected at W2.5 (Fig. 4A).

**Fig. 4.**
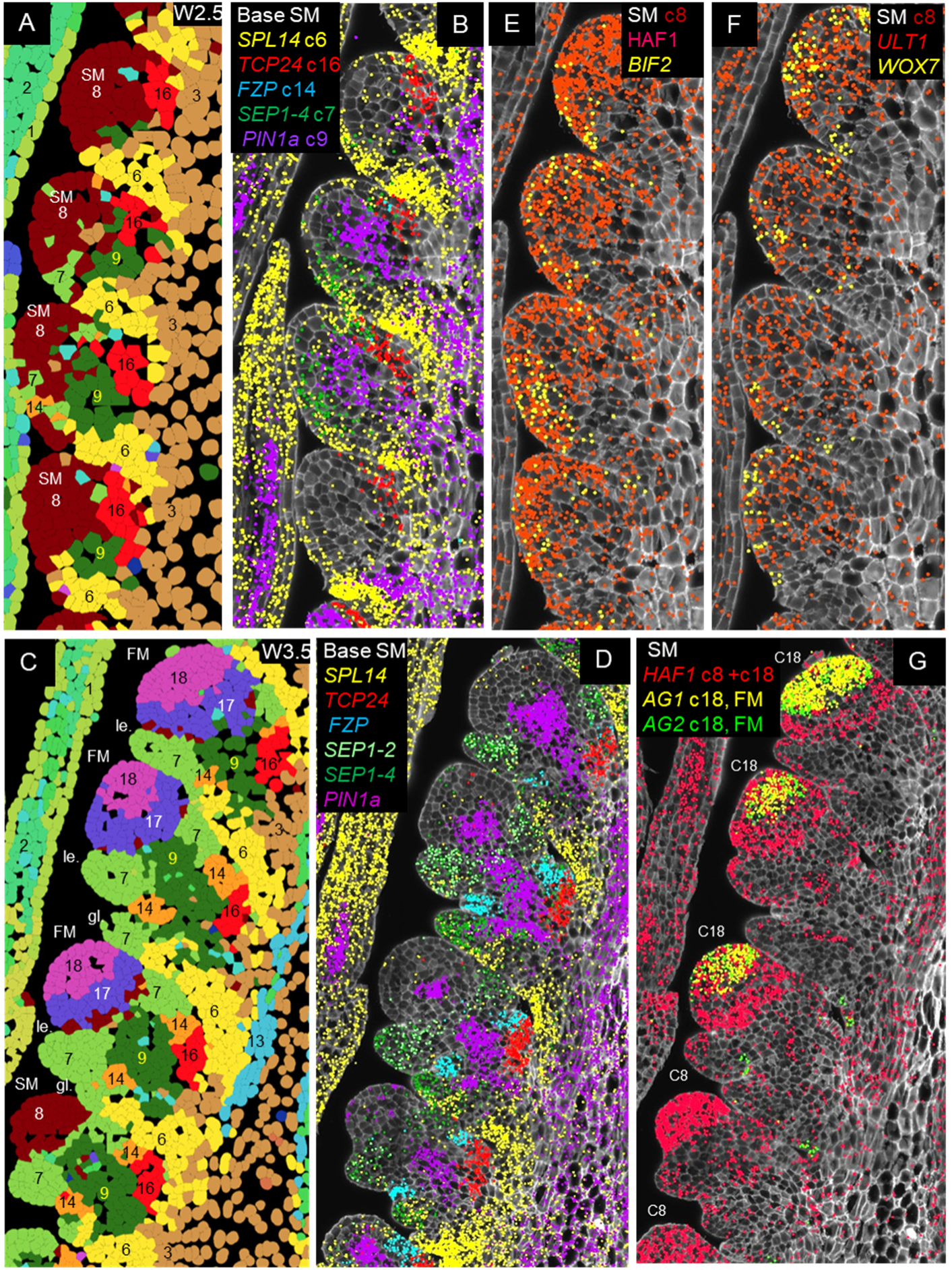
Expression profiles of genes differentially expressed during spikelet development. **A-B, E-F** W2.5 (late double ridge stage). (**C-D, G**) W3.5 (floret primordia stage). **A, C** Cell segmentation and clustering. **B, D** Genes differentially expressed at the base of the SM. **E-G** Genes differentially expressed in the SMs. In **G**, the more basal SMs are at an earlier stage (cluster c8) than the central spikelets, which have already transitioned to the floret development stage (FM, cluster c18), and are already expressing floral homeotic genes *AG1* and *AG2*. *SEP1-4* marks the developing glume (stronger) and lemma, *SEP1-2* the developing lemma, *PIN1a* the developing vasculature, *SPL14* the suppressed bract, *FZP* the glume axilla and *TCP24* the adaxial boundary. Complete spike images showing expression of these genes are presented in Figs. S15-S18. SM= spikelet meristem, FM= floret meristem, le.= lemma, gl.= glume.

##### 1.4.1. Cluster c6 “suppressed bract”

Cluster c6 includes the suppressed bract region (Fig. S14). This cluster showed a relatively high correlation with cluster c19 in the transition zone (*R*= 0.71, Fig. S14 and Data S3), which was also associated with leaf ridge repression. Both clusters showed preferential expression of *SPL14* (Fig. S14A-B)*, FUL2*, and *LFY* (Data S2), but cluster c6 differed from cluster c19 by its preferential expression of *SPL13* (Fig. S14C-D) and lower expression of *TB1* and *TB2*. In both c6 and c19 clusters, preferential expression of *SPL* genes was associated with higher *VRN1* and *FUL2* expression (Fig. S14E-F). Combined *vrn1 ful2* mutations resulted in bract outgrowth at the basal spikelets (Fig. S14G), suggesting a role of c6 and c19 cells in bract repression [3].

##### 1.4.2. Cluster c9 “Spikelet base-center”

Cluster c9 is located at the center of the region proximal to the SM (Fig. 4) and showed high correlations (*R=* 0.75-0.80) with the surrounding clusters c3, c7, c8, c14, and c16 (Data S3). This central cluster was enriched in *PIN1a* (Fig. 4A-D), *PIN1b,* and *LAX4* (Data S2), suggesting a role in vascular development and the transport of water and nutrients to developing spikelets. Additional genes preferentially expressed in c9 included *SOC1-1*, *SEP1-4, OSH1* and *TB1*.

##### 1.4.3. Cluster c7 “Glume and lemma”

The glume and lemma primordia, which became visible at W3.5, were both included in cluster c7. Both organs showed preferential expression of *AP2L5* (Fig. S6B), *FUL2* and *VRN1* (Fig. S14E-F) and reduced expression of *OSH1* (Fig. S12E) and could be differentiated by the preferential expression of *SEP1-2* in lemmas and *SEP1-4* stronger expression in glumes than lemmas (Figs. 4C-D, S15B). The latter result is consistent with previous reports in rice [43]. Cluster c7 showed the highest correlation with the early meristem cluster c8 (*R=* 0.81) from which glumes and lemmas originate, and also relatively high correlations with the leaf cell clusters (*R=* 0.63-0.75, Data S3). This latter result supports the hypothesis that glumes and lemmas are modified leaves [44]. Similar to leaves, glumes and lemmas later turn green, contributing photosynthetic products to flowers and developing grains, in addition to additional protection to the reproductive organs.

##### 1.4.4. Cluster c16 “Adaxial boundary”

This cluster was located adaxial to c9 (Figs. 4A-D, S15) and was defined by the differential expression of the transcription factor *TCP24* (Fig. S15, Data S2). *TCP24* and its closest paralogs *TCP22*, *TB1*, and *TB2* belong to the CYC/TB1 subclass of TCP transcription factors (Fig. S11E-F) and can form homo and heterodimers [45]. In the maize male inflorescence [46] and the *Brachypodium distachyon* spike [47], the orthologue of *TCP24, BRANCH ANGLE DEFECTIVE 1* (*BAD1*), is preferentially expressed in the pulvinus and affects branch or spikelet angle, respectively. The pulvinus is located at the axilla of the tassel branches in maize and at the spikelet axilla in *Brachypodium* but is absent in the *Triticeae species*. Loss-of-function mutations of *TCP24* in rice (=*OsTB2* = *RETARTDED PALEA 1*) are associated with mild reductions in the size of the palea with no other obvious effects on inflorescence or flower development [48]. By contrast, loss-of-function mutations in the barley ortholog *HvTB2* (also known as *COMPOSITUM1*) result in spikes with branches in the basal region [45], suggesting functional differentiation among grass species.

##### 1.4.5. Cluster c14 “Glume axilla”

Cluster c14 is defined by the expression of *FRIZZY PANICLE* (*FZP,* Fig. S15B, Data S2), which is expressed at the axillae of the developing glumes (Fig. 4C-D). A similar localization has been observed in previous *in situ* hybridization studies in rice and maize [49, 50]. In spike longitudinal sections, c14 cells frequently appear as two small cell clusters flanking c9, with the adaxial one adjacent to c16 (Figs. 4C-D, S15, S16). Cluster c14 also showed preferential expression of the homeobox domain gene *WOX7* (Fig. S16A) and the PINOID-like serine/threonine kinase *BIF2* (Fig. S16B), although these genes are also expressed outside of c14. Previous studies have shown that *FZP* plays an important role in the repression of the glumes’ axillary meristem. Natural or induced amino acid changes in FZP result in the formation of branches at the base of the spikes in “miracle wheat” (*bh1* mutant) and barley (*com2* mutant) [51, 52].

To characterize the function of *FZP* in tetraploid wheat, we generated loss-of-function mutations in both *FZP* homeologs by CRISPR (Fig. S17). Eight independent T_0_ lines with mutations in both *FZP* homeologs showed spikelets replaced by dense arrays of nested glumes (Fig. S17B and E). Dissection of these structures revealed new axes emerging from the axillae of the initial glumes, each producing multiple secondary glumes. The axillary meristems of these secondary glumes repeat the cycle but produce fewer and smaller tertiary glumes. The patterns repeat until the meristems were exhausted (Fig. S17F). Poorly developed floral organs were occasionally observed among these recurrent glumes (Fig. S17G). The rachilla continued its development but only vestigial florets were observed in its nodes (Fig. S17H). A diagram describing the development of this structure is presented in Fig S17I, and the dissection of a wildtype spikelet and the corresponding diagram in Fig. S17J-K. In some distal nodes, we observed lemmas with longer awns subtending more normal floral organs. These results indicate that *FZP* is required to suppress the axillary meristems in the glumes to ensure normal floret development. Similar phenotypes have been described for *fzp* loss-of-function mutants in rice and maize [49, 50], suggesting a conserved function in grasses.

In summary, the different expression domains detected in the spikelet region proximal to the SM is consistent with recent grass inflorescence models proposing that the basal region of the grass spikelets as an important signaling center for spikelet development [53].

#### 1.5. Spikelet meristem (SM) and floret meristem (FM) regions

At all three developmental stages, cells in the IM and early SMs were assigned to cluster c8. At W2.5, this cluster exhibited preferential expression of *HAF1*, *BIF2*, (Figs. 4E, S18A), *ULT1* and *WOX7* (Figs. 4F, S18C-D). *HAF1*, a C3HC4 RING domain-containing E3 ubiquitin ligase, has no reported role in inflorescence development [54, 55], but the other three genes are well-established regulators of meristem function. *BIF2* regulates auxin transport and is required for axillary meristem initiation and lateral primordia development in maize and rice [56]. *ULT1* encodes a SAND domain transcription factor that controls shoot and FM activity in Arabidopsis [57, 58], whereas *WOX7* is a *WUSCHEL*-related homeobox gene that contributes to inflorescence architecture in rice [59]. Based on their physical location and preferential gene expression, we propose that c8 cells contribute to meristem function.

At W3.5, the more developed central spikelets showed the formation of floral meristems (Fig.S1B) that were assigned to cluster c18 (FM, Figs. 1, S2). This cluster was defined by the expression of floral homeotic genes *SEP3-1*, *SEP3-2* (Fig. S18E), *PI1*, *AP3* (Fig. S18F), *AG1* and *AG2* (Figs. 4G, S18G, Data S2), strongly supporting a role in floral development. The SMs are not visible in the sections of the central spikelets (Fig S1), either because the larger FMs displaced the adjacent SMs from the section plane or because the section was not perfectly centered (Fig. S1E).

Cluster c18 also showed preferential expression of *WAPO1*, a gene encoding a protein that interacts with LFY (Fig. S18H). Loss-of-function mutations in wheat *WAPO1* or *LFY* result in downregulation of floral homeotic genes and floral defects in wheat [13, 14] and other grasses [60–62].

The floral meristems in c18 are separated from the lemma primordia (c7) by a group of cells forming a distinct band assigned to cluster c17 (Figs. 1, 4C). This cluster was defined by the preferential and high expression of *LFY* (Fig. S18H, Data S2). A previous study suggested that the overlap between the *WAPO1* and *LFY* expression domains (Fig. S18H) provides critical spatial information for correct flower development in wheat [13]. The dynamic changes in gene expression during spike development are described later in section 1.7.

#### 1.6. Changes within the inflorescence meristem (IM) during the transition to a terminal spikelet (IM→TS) and *SPL14* functional validation

In wheat, the rate of lateral meristem formation and/or the timing of IM transition to a terminal spikelet (IM→TS) determine SNS. Our previous studies showed that loss-of-function mutations in *LFY* or *WAPO1* reduce the rate of SM formation and SNS [13]. The reductions in SNS were comparable in the single and combined *lfy wapo1* mutants, suggesting that the LFY-WAPO1 protein complex is essential for establishing and/or maintaining a normal rate of SM generation. However, *LFY* and *WAPO1* are co-expressed in the IM only during the earliest stages of spike development (W1.5), suggesting that the early presence of the LFY-WAPO1 protein complex may induce a persistent change in IM function [13].

Since changes in the timing of the IM→TS transition can also influence SNS, we focused on genes involved in the early stages of this transition. We were unable to identify marker genes that clearly distinguished IMs from early SMs (both grouped in cluster c8). Because spikelets are “small spikes”, SMs are also inflorescence meristem, making the identification of differential markers between IMs and early SMs more challenging. However, as SMs begin to differentiate into spikelets, an increasing number of genes become differentially expressed relative to the IM.

We manually delineated the apical meristem region and compared the hybridization density of the 99 genes in this region before the IM→TS (W2.5) and at the early stages of the transition (W3.5). These are within-gene comparisons and, therefore, were not influenced by probe hybridization efficiency. For each stage, we analyzed four sections (four biological replications for W2.5 and two for W3.5, each with two subsamples, see M&M). To verify that the selected W3.5 sections had already initiated the IM→TS transition, we confirmed the expression of *FZP* in the youngest lateral meristems (LM, Fig. S19). The selected region encompassed both the terminal spike meristem (TM) and the two youngest LMs (henceforth, TM+2LM). At W2.5 the TM corresponds to the IM and LMs to the SMs, whereas at W3.5 the TM corresponds to the SM of the terminal spikelet and the LMs the future glumes. Using this approach, we previously observed a significant decrease in *LFY* expression and a significant increase in *FUL2* expression between W2.5 and W3.5 in the TM+2LM region [13].

In this study, we identified 11 additional genes with significantly altered hybridization densities (spots/100 μm^2^) in the selected region between W2.5 and W3.5 (*P*< 0.05, Fig. S20, Data S5). Among them, *SPL14* exhibited a highly significant decrease in hybridization density during the transition, (6.7-fold reduction, *P=* 0.0031, Figs. S19, S20, Data S5). We also identified two additional MADS-box genes (*SOC1-2* and *SEP1-4*), four bZIP transcription factors (*bZIPC1*, *bZIPC3*, *FDL2*, and *FDL6*), one bHLH transcription factor (*BC1*), one auxin efflux carrier (*PIN1b*), and one receptor-like kinase (*CRN*) involved in the CLV-WUS negative feedback loop (Fig. S20) that were significantly upregulated during the transition. Several of these genes have been associated with inflorescence phenotypes. For example, mutations in *bZIPC1* in wheat drastically reduce SNS [63], whereas mutations in the rice ortholog of *SEP1-4* (*OsMADS5*) impact inflorescence branching [64].

To explore the phenotypic effect of reduced *SPL14* expression during the IM→TS transition, we generated loss-of-function mutations in both *SPL-A14* and *SPL-B14* using CRISPR (Fig. S21A-B). Plants with homozygous mutations for both homeologs showed a premature IM→TS transition and a 42% reduction in SNS (Fig. S21C and E, Data S6). These results suggest that *SPL14* plays an important role in delaying the timing of the IM→TS transition in durum wheat. In addition, the *spl14* mutant plants were 12 cm taller (Fig. S21D and F), headed two days later (Fig. S21G), and produced 2.5 more leaves than the wildtype (Fig. S21H, Data S6), indicating that *SPL14* plays additional roles. Similar effects on SNS and heading time have been previously reported in *spl14* CRISPR mutants in hexaploid wheat, and natural alleles have been identified with favorable effects on wheat grain yield [4]

Since SPL proteins directly regulate the expression of *SQUAMOSA* MADS-box genes, we hypothesize that the reduced *SPL14* expression may be linked to the correlated increase in *FUL2* and *VRN1* expression in the IM observed during the IM→TS transition. Supporting this hypothesis, we previously showed that a mutation in a predicted *SPL*-binding site in the *VRN1* promoter was associated with a significant increase in *VRN1* expression during early spike development [65]. In summary, we identified multiple wheat genes involved in the early IM→TS transition and validated the association between *SPL14* and the final number of spikelets per spike.

#### 1.7. Time and pseudotime changes in spike development

Spike and spikelet developmental changes over time were explored across three developmental stages (W1.5, W2.5 and W3.5), along the longitudinal axis of the spike, and through trajectory analysis (Fig. S22). At the early double ridge stage (W1.5), the IM and the upper ridge cells were included in cluster c8 (early meristem) and a few cells in cluster c6 (bract repression, Fig. 1A and S5). These two cell clusters expanded at the late double ridge stage (W2.5, Fig. 1B), and a gradient of spikelet differentiation started to form along the longitudinal axis, with the most advanced spikelets located at the center of the spike. This developmental gradient became more pronounced at W3.5, when well-developed floral meristems were observed at the central spikelets (Fig. 1C, clusters c17 and c18).

These dynamic changes were captured in a trajectory analysis (Fig. S22), which used cluster c8 (IM and early SM meristems) as the root. From c8, two trajectories diverged in opposite directions: one leading to the formation of floral meristems (c17 and c18) and the other branching towards the formation of the transition zone (c19), the repressed bract (c6) and the central zone (c3). A third trajectory diverged into three branches: one leading to the central spikelet (c9), another to the glume and lemma primordia (c7), and a third one to the boundary clusters c14 and then c16 (Fig. S22). Cluster c9 was also connected to the suppressed bract trajectory (Fig. S22).

These predicted trajectories reflected many of the developmental changes observed during spike development but also showed a few discrepancies. Although the c14 cluster appeared earlier than c16 in the predicted trajectory, c16 cells were observed at the base of the developing spikelets earlier than those assigned to c14, both across developmental stages and along the longitudinal gradient of the spike (Fig. 1 and S2). A closer examination of Fig. S22 showed that multiple c16 cells connect directly to the meristematic region (c8), suggesting an alternative trajectory from c8 to c16 independent of c14.

The comparison of the different spikelets within a spike provided a precise view of the dynamic gene changes leading to the formation of the FM. The meristematic markers *HAF1* and *ULT1* were expressed in both early and advanced meristems, whereas *BIF2* and *WOX7* expression in the meristems was restricted to the youngest SMs (Fig. S18B and D). In more advanced SMs or FMs, *BIF2* and *WOX7* were detected in c14 and other clusters (Fig. S16). In SMs slightly more developed than those expressing *BIF2* and *WOX7*, we detected the MADS-box gene *AGL6*, which is known to play a role in paleae and lodicule development [12]. Finally, in the more developed central spikelets, *AGL6* expression was restricted to a region between the lemma primordia and the FM that overlapped with c17, and the FM showed expression of floral homeotic genes and *WAPO1* (Fig. S18E-H).

### 2. Single-cell RNA sequencing (scRNA-seq) of the developing wheat spike

To identify additional genes regulating wheat spike development, we conducted single-cell RNA-seq (scRNA-seq) using protoplasts isolated from *Kronos* spikes at the W2.5 and W3.5 stages (Fig. 5A). We optimized the cell wall digestion method to enhance meristem cell recovery, following a previous single-cell study on maize inflorescence stem cell [66] (see Methods). Raw scRNA-seq data are available in the NCBI Sequence Read Archive (SRA) under BioProject accession number PRJNA1199502. Reads were mapped to the Kronos reference genome v1.1 (annotation v1.0) [67].

**Fig. 5.**
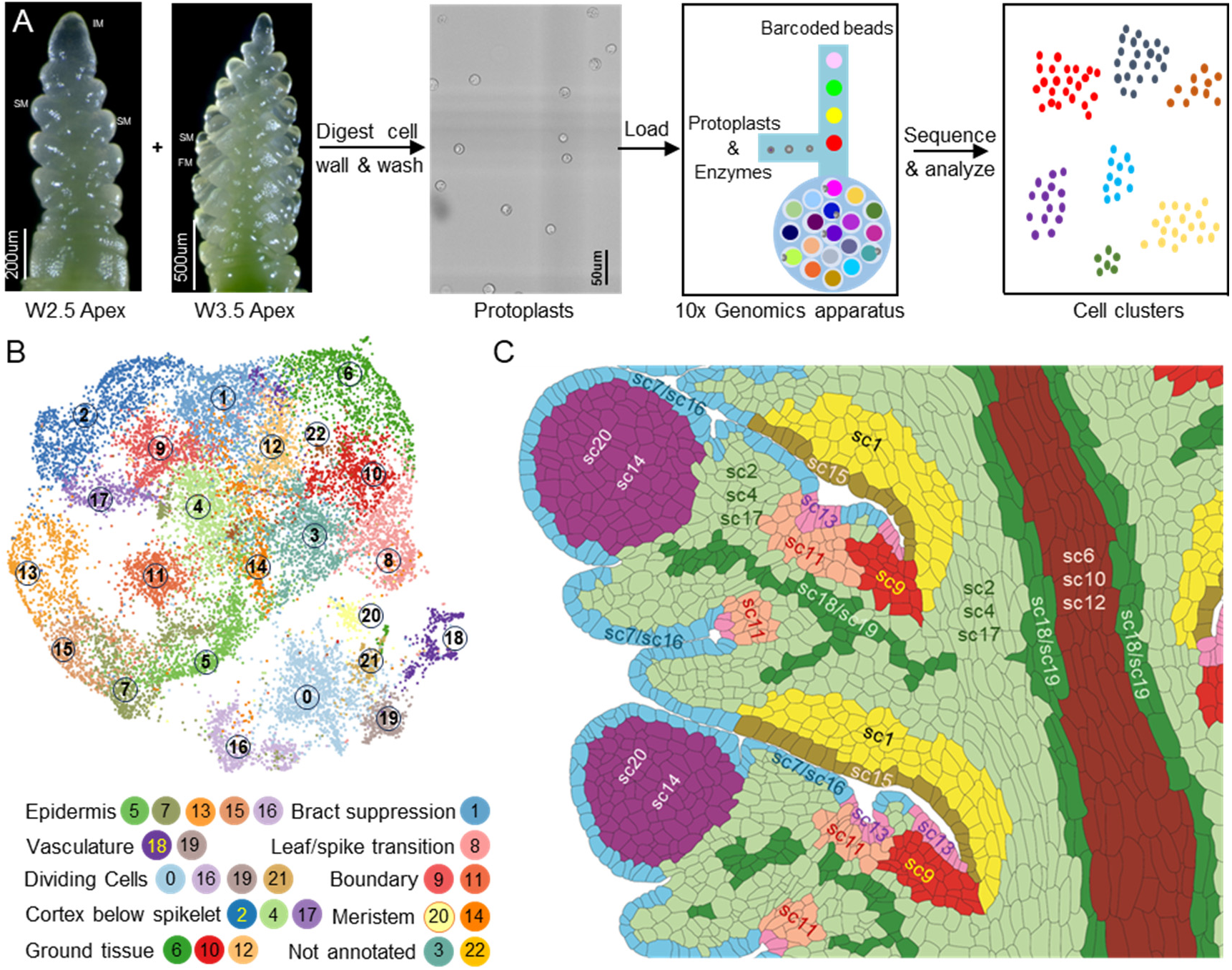
Single-cell RNA-seq (scRNA-seq) analysis of wheat inflorescence spikes at the W2.5 and W3.5 stages. **A** Experimental workflow, including wheat protoplast isolation and subsequent scRNA-seq. **B** UMAP plots showing cell clustering and their annotations, **C** Cartoon with inferred positions of the different scRNA-seq clusters based on information discussed in the next sections. The cartoon is based on the same two central spikelets shown in Fig. 4C, allowing for direct comparisons with the smFISH clusters.

In the initial analysis, we profiled 12,304 and 13,705 individual cells from three independent replicates at W2.5 and W3.5 stages, respectively (Data S7). Good integration was observed among both replicates and stages (Fig. S23A). We detected expression of 67,629 genes at W2.5 and 70,047 genes at W3.5 (Data S7), comparable to the number of genes detected in bulk RNA-seq of Kronos spikes at the same developmental stages [68]. The mean and median unique molecular indices (UMIs), the number of detected genes per cell, and other statistics are reported in Fig. S23A and Data S7. Cell clustering of 26,009 cells from both stagers using Seurat v5 [69] identified 25 distinct clusters in the initial analysis (Fig. S23A). Nine clusters were identified with cells at either the G2/M or S phase, marked by *CYCB2* and *HIS2A1* expression, respectively (Fig. S23B-C).

To improve the resolution of biologically relevant clusters, cells from the nine cell-cycle clusters in the initial analyses were removed, and the remaining 17,283 cells were re-clustered into the final 23 clusters described in this study (sc0 to sc22, Fig. 5B). The final single-cell clusters were visualized using two-dimensional (2D) uniform manifold approximation and projection (UMAP) plots (Fig. 5B) and a 3D-UMAP (https://dubcovskylab.ucdavis.edu/content/kronos-scrna-3d-clusters). We also developed a Shiny App (https://dubcovskylab.ucdavis.edu/content/kronos-integrated-clusters) to visualize gene expression across clusters. To quantify the relationships among clusters, we calculated the average normalized gene expression across all cells for each cluster and used these expression profiles to calculate Pearson correlations for all cluster pairs (Data S8).

#### 2.1. Characterization of scRNA-seq cell clusters and integration with spatial clusters

To annotate the scRNA-seq clusters, we identified and annotated genes preferentially expressed in each cluster (Data S9). We used this annotation to identify known marker genes (Data S1) and to perform KEGG pathway analyses (Data S10). We also incorporated spatial information into the scRNA-seq clusters using RNA-seq data from the basal, middle, and top sections of Kronos developing spikes at W3.0 (Fig. S24 and Data S11), as well as from correlations with published spatial transcriptomics clusters from hexaploid wheat (Data S13-15) [21].

To integrate the scRNA-seq and smFISH data, we used an imputation approach recently published in barley that incorporates scRNA-seq information into smFISH-labelled cells while retaining their spatial information [70]. Briefly, we analyzed the cells from both studies together using the 91 common genes and for each smFISH-labelled cell we identified the five closest scRNA-seq cells. We then used the average expression of these five cells (weighted by their distances) to impute the scRNA-seq expression into the smFISH-labelled cell (see Materials and Methods). We developed a web tool to visualize the spatial expression profiles of the 74,464 imputed genes in two spike sections at W2.5 and two at W3.5 (https://dubcovskylab.ucdavis.edu/imputed-genes). This tool greatly expanded the spatial information available to annotate the scRNA-seq clusters.

We used a similar approach to project the smFISH cluster ID into the scRNA-seq cells (see Materials and Methods). Since the number of genes in the smFISH clusters affected the number of identified cells, we divided the number of projected cells by the total number of cells in the smFISH clusters and generated a table including the “%-contribution” of each smFISH cluster to each scRNA-seq cluster (Data S12).

The different sources of information used to annotate the scRNA-seq clusters are summarized in Data S16 along with the final cluster annotations. The use in these annotations of similar names as the smFISH clusters indicates a better overlap than with other clusters and not a perfect one-to-one correspondence. The tentative physical locations of the scRNA-seq clusters are presented schematically in Fig. 5C as a reference for the evidence discussed in the sections below, which are grouped by similar functions.

#### 2.2. Cell cycle enriched clusters

Although cells from the initial cell-cycle clusters were removed, four of the 23 final clusters exhibited enrichment in cell-division marker genes combined with markers for other tissues. The S-phase markers *HIS2A1-1* and *HIS2A1-2* were enriched in clusters sc0, sc16, sc19, and sc21 (Fig. S25). Cluster sc16 was also enriched in G2/M phase cyclin markers (*CYC7*), cyclin-dependent kinases (*CDKB2*), and the *GROWTH REGULATING FACTORs GRF2*, *GRF4*, and their cofactor *GIF1* (Fig. S25). Gene IDs and references for all marker genes are provided in Data S1.

Among the genes preferentially expressed in cluster sc21 (Data S9) that exhibited significant differences in the RNA-seq study, 65% were expressed at higher levels in the base of the spike (Data S11-S16) suggesting a predominantly basal location for sc21. The smFISH cluster with the highest contribution to sc21 was c3 (central spike) supporting the previous result (Data S12). The basal and central region of the Kronos spike was characterized by a green coloration (Fig. 5A), indicative of the presence of chlorophyll. This observation suggested that sc21 was enriched in genes within the photosynthetic pathway, a hypothesis supported by the KEGG analysis (Data S10). Based on these findings, cluster sc21 was annotated as “cell cycle / photosynthetic ground tissue / basal region” (see section 2.5 for additional support).

Most of the genes preferentially expressed in sc0, sc16, and sc19 (Data S9) were expressed at higher levels in the distal third of the developing spikes (Data S16). This enrichment was strongest in sc0 (100% of the genes), intermediate in sc16 (78%), and weak but significant in sc19 (47%). Based on its distal position and enrichment in meristematic markers (section 2.10), sc0 was annotated as “cell cycle / meristematic / distal region”. Among the smFISH clusters, the one with the highest %-contribution to sc0 was c8 (early meristem) supporting the previous result (Data S12). Cluster sc19 was also enriched in vascular markers (see section 2.4), and among the smFISH clusters projected into sc19 the highest value was for c13 (vasculature, Data S12). Based on these results, sc19 was annotated as “cell cycle / vasculature / enriched in spike tip”. Finally, sc16 showed multiple epidermal markers (see section 2.3) and KEGG enrichment in cutin, suberin, and wax biosynthesis (Data S10),which supported its annotation as “cell-cycle / epidermis / enriched in spike tip”. Additional information for sc16 is provided in the next section on epidermal markers.

#### 2.3. Epidermal clusters

A similar enrichment in cutin, suberin, and wax biosynthesis was detected in the KEGG analysis for cluster sc15, supporting its annotation as an epidermal cluster. Clusters sc5, sc7, and sc13 were also annotated as epidermal clusters based on the preferential expression of known epidermal markers, including *ONI* (synonymous with *KCS10*), *PEL3*, *SCR*, *ROC2*, *ROC3*, *ROC7* (Data S1, Fig. S26). The Kronos imputed expression data confirmed the epidermal expression of these six genes (Fig. S26H). Since no epidermal markers were included in our smFISH experiment, these results exemplify the power of the imputation approach.

We compared the single-cell epidermal clusters to a recent hexaploid spatial transcriptomics study that identified two expression domains, ED1 and ED4, associated with epidermal cells [21]. These two epidermal expression domains had the highest correlations with sc7, sc13, sc15, and sc16 (Data S13-15), further supporting the epidermal annotation of these clusters. However, their correlations with sc5 were intermediate (Data S13-15). To identify markers that differentiate the five epidermal clusters, we performed a separate comparison including only these clusters and summarized the results in Data S17.

Epidermal clusters sc16 and sc7 were highly correlated (*R=* 0.79) and were both enriched in genes with higher expression levels at the spike tip (sc7= 50% and sc16= 69%). These clusters also showed high expression levels of ribosomal genes (Data S16), suggesting rapid cell growth and division. Cluster sc16 showed preferential expression of multiple cell-cycle genes (Fig. S25A-G, Data S17) and higher correlations with cell-cycle clusters sc0, sc19 and sc21 than all other epidermal clusters (Data S8). Cluster sc16 was also enriched in meristematic markers *WOX3a* and *WOX3b* (Data S17). The smFISH cluster c8 (early meristem) showed the highest %-contribution to sc16 relative to other smFISH clusters, supporting the previous result (Data S12). Taken together, these results justify the annotation of sc16 as “cell cycle / epidermis / enriched in spike tip”.

Epidermal cluster sc7 was enriched in both meristematic and lateral organ markers. The enriched meristematic markers included *AGL6* (Fig. S18B and S27A), and the imputed *WOX3b* (Fig. S27J) and *CLE33* (Fig. S27K). Cluster sc7 was also enriched in *YAB3* and *YAB4* genes (Fig. S27C, Data S17), which were detected at the lower ridge and lateral organs in hexaploid wheat [21]. This expression pattern was validated in our imputed expression data (Fig. 27L). Correlations between epidermal clusters sc7 and sc16 and the hexaploid wheat expression domains ED4 (meristem epidermis) and ED1 (glume and lemma epidermis) [21] were the highest in both directions and in all three developmental stages (Data S13-S15). In addition, the smFISH clusters with the highest %-contribution to sc7 were c7 (glume and lemma) and c8 (early meristem) relative to other smFISH clusters (Data S12). Taken together, these results support the annotation of epidermal cluster sc7 as “epidermis / meristem and lateral organs” (Figs. 5C).

Epidermal cluster sc15 showed the second highest correlations with ED1/ED4 after sc7/sc16 in hexaploid wheat (Data S13-15). However, sc15 differed from sc7 by its lower expression of genes encoding ribosomal proteins and in its more uniform distribution along the spike (Data S16). Among the markers genes identified for sc15, *SPL13* and *SPL14* (Fig. S26G and S27D-E) suggest that sc15 may correspond to the abaxial base of the spikelets including the suppressed bract region (Fig. S14A-D). This hypothesis is also supported by the higher correlation observed between sc15 and the suppressed bract cluster sc1 (*R=* 0.63) than with meristematic clusters sc14 and sc20 (Data S8). Consistent with the previous result, the smFISH cluster with the highest relative %-contribution to sc15 was c6 (suppressed bract, Data S12). Based on these results, sc15 was annotated as “epidermis / suppressed bract” (Fig. 5C).

Epidermal cluster sc13 showed a relatively high correlation with sc15 (*R=* 0.77) but differed by its preferential expression of *TCP24*, a marker gene for the adaxial boundary domain (Fig. 4A and C). Cluster sc13 was enriched in additional boundary markers such as *TCP22, CUC3, LAX1*, and *DP1*, which have been found to be preferentially expressed in the boundary domain in hexaploid wheat [21] and in our imputed expression profiles (Figs. S27M-P, see also later section 2.12). The sc13 cluster sowed higher correlations with the hexaploid wheat expression domain ED12 (boundary domain) [21] than with epidermal clusters ED1/ED4 and all other clusters (Data S13-15). Relative to other smFISH clusters, c16 (adaxial boundary) showed the highest %-contribution to sc13 (Data S12). Based on these results, sc13 was annotated as “Epidermis / adaxial boundary” Fig. 5C.

Finally, sc5 was the most divergent of the epidermal cluster, showing the lowest correlation with other epidermal clusters, the lowest expression levels of epidermal markers *SCR*, *PEL* and *ONI* (Fig. 26G), and the largest number of differentially expressed genes (Data S17) with two examples in (Fig. S27H-I). The smFISH clusters with the highest contribution to sc5 was c19 (transition zone), suggesting an enrichment in epidermal cells from the transition zone. However, based on sc5 intermediate/high levels of protoplasting induced genes (Data S16), high correlations with unknown clusters sc3 and sc22 (Data S8), and uniformly low correlations with spatial clusters in hexaploid wheat (Data S13-15), we designated sc5 as “Epidermis / unknown”.

#### 2.4. Vascular clusters

The vascular genes *APL*, *LAX4*, *CEN5*, *PIN1a*, and *PIN1b* characterized by smFISH (Fig. 3) were all preferentially expressed in sc18 and sc19, along with additional known vascular marker *SHR1*, phloem markers *JUL1* and *DOF19*, and xylem markers *TMAAT, TMO5L1, TMO5L3*, *WAT1*, and *XCP1*, *LOG10b*, and *NAC52* (Fig. S28, Data S1). The vascular location of the last ten markers was validated in the Kronos imputed expression profiles (Fig. S28H), together with *APL* and *LAX4* controls characterized also by smFISH (Fig. 3). The smFISH cluster with the highest contribution to sc18 and sc19 was c13 (vasculature, Data S12). These results support the annotation of sc18 and sc19 as vascular clusters (Fig. 5C).

Both vascular clusters shared relatively high expression levels of *OSH1* and genes encoding ribosomal proteins (Data S16). However, the genes preferentially expressed in sc19 (Data S9) were expressed at higher levels at the tip of the spike (Data S16), whereas those preferentially expressed in sc18 (Data S9) were expressed at higher levels at the base and middle of the developing spikes (Data S16). Since sc19 was enriched in *HIS2A* expression and sc18 was not (Fig. S25), sc19 was annotated as “cell cycle / vasculature / enriched in spike tip” and sc18 as “vasculature / enriched in spike base and middle”.

Cluster sc18 showed two distinct regions: one preferentially expressing phloem markers (Figs. S28B and D) and the other expressing xylem markers (Figs. S28C, E-F). To compare these regions, we further divided sc18 into three subclusters: sc18_0 with the highest correlation to the ground tissue cluster c3 (*R*= 0.625), sc18_1 including phloem markers *JUL1* and *DOF19*, and sc18_2 including xylem markers *WAT1*, *XCP1*, *TMO5L1*, *LOG10b* and *NAC52* (Fig. S28H). Genes differentially expressed in sc18_1 and sc18_2 are available in Data S18.

Consistent with the key role of hormones in vascular development [71, 72], sc18 showed preferential expression of both homeologs of the auxin signaling genes *PIN1*, and *ARF11*, and the JA biosynthetic gene *OPR3*. Cluster sc18 also showed preferential expression of the *CPS1* gene encoding a GA biosynthesis enzyme, and genes *CKX-B3*, *CKX-B5*, and *CKX-B11*, all involved in cytokinin degradation (Fig. S28I).

Interestingly, the floral homeotic genes *AP3* and *PI1* (Fig. S18F) were also detected in cluster sc18 (Figs. S29A-B). A closer examination of the smFISH profiles revealed that these two MADS-box genes were co-expressed with the vascular marker *APL* at all three developmental stages (Figs. S29C-E). Expression of class-B MADS-box genes in the developing vasculature has been reported in a few eudicot plant species [73]. Our findings extend this conserved function to monocots.

#### 2.5. Central spike clusters

In addition to vascular and pro-vascular cells, the central region of the developing spike contains ground tissue cells, which correspond to cluster c3 (Fig. 1 central spike) in our smFISH study, and expression domains ED3 (developing rachis) and ED5 (rachilla) in the hexaploid wheat study [21]. These ground tissue cells showed a green coloration indicative of chlorophyll presence, which was stronger at the base of the spike (Fig. 5A). The KEGG analysis indicated that genes preferentially expressed in single-cell clusters sc6 and sc10 were enriched in photosynthesis pathways (Data S10). Eighty two percent of the top 50 genes preferentially expressed in sc6 were annotated with photosynthesis-related functions (Data S9), including *CAB2R* and *LHCB1.1*, which were detected in the central spike region in hexaploid wheat [21] and were validated in the Kronos imputed expression data (Fig. S30A-H). The expression levels of these photosynthetic genes across all clusters are presented in a heat-map in Fig. S31.

The highest average enrichment in photosynthetic genes was detected for sc6 followed by cell-cycle cluster sc21 (Fig. S31, Data S16). This finding supported the previous annotation of sc21 in section 2.2 as “cell cycle / photosynthetic ground tissue / basal region”. Enrichment in photosynthetic genes was also observed in sc10 and sc12, but at decreasing levels relative to sc6 (Data S16 and Fig. S31). A large proportion of the genes preferentially expressed in sc10 (44%) and sc12 (36%) were also preferentially expressed in sc6 (Fig. S32), supporting the connections among these clusters.

The genes preferentially expressed in sc6, sc10, and sc12 showed higher levels of expression at the basal region of the developing spikes based on the RNA-seq data (Data S11-S16). This location was further supported by the enrichment of *FWL3* and *SVP1* in both sc6 and sc10, which are expressed in the central and basal parts of the spike (Figs. S32A-C and H). In addition, both sc10 and sc12 were enriched in *OSH1*, a gene highly expressed in the central spike region (Fig. S12). Cluster sc6 was enriched in additional genes expressed in the central spike region, including *AP3*, *bZIPC3*, *FDL15*, *FT2*, *OSH1*, *SPL3*, and *VRT2* (Figs. S12 and S30), as well as *CAB2R* and *LHCB1.1* both in hexaploid wheat and in our imputed expression data (Fig. S30H). The smFISH cluster with the highest relative %-contribution to sc6, sc10 and sc12 was c3 (spike central, Data S12), supporting a central spike location for the cells of these three clusters. Based on the above information, clusters sc6 and sc10 were annotated as “central spike / photosynthetic ground tissue / spike base”, and sc12 as “tentative ground tissue / enriched spike base” (Fig. 5C).

#### 2.6. Transition zone between leaves and spike

The transition zone between the leaves and the spike is represented by the smFISH cluster c19 (Figs. 1, S5), which shared several preferentially expressed genes with scRNA-seq cluster sc8. These genes included *TB1*, *TB2*, *SPL17*, *GNI2*, *IDD10*, and *GA20ox1* (Fig. S33A) from our smFISH study. This region was also enriched in *LEC1* and *TRD1*, two genes identified in the hexaploid wheat data [21] and validated in our imputed expression data (Fig. S33B-F). Cluster sc8 also showed high enrichment in jasmonic acid signaling genes *MYC2* and *JAZ8*, although both genes were also expressed in other clusters (Fig. S33E-F).

Among the genes preferentially expressed in sc8, *SPL17* and its downstream target *TRD1* [30] (ortholog of *OsNL1* and *ZmTSH1*) were previously shown to play important roles in bract suppression in the transition zone. Loss-of function mutations in *SPL17/SPL14* in hexaploid wheat [4], and its orthologs in rice [30] and or maize [5] were associated with bract outgrowth at the base of the inflorescences. A similar phenotype was observed in *TRD1* mutants in barley [74], rice *OsNL1* [75], or maize *ZmTSH1* [76]. This phenotype was consistent with the significantly higher expression at the base of the spike of *SPL17*, *TRD1* (Data S11). Among the genes in the RNA-seq experiment with significant differences among the three sections of the spike, 66% of the ones expressed in sc8 showed higher levels at the base of the spike (Data S16). Among the smFISH clusters, the one with the highest contribution to sc8 was c19 (transition leaf/spike) followed by c3 (central spike, Data S12). Based on these results, sc8 was annotated as “transition leaf-spike / spike base”.

#### 2.7. Suppressed bract cluster

Cluster c6, which includes the suppressed bract cluster, was consistently detected in the smFISH study at the base of each developing spikelet (Fig. 1). This cluster showed strong differential expression of *SPL13* and *SPL14* (Data S2), two markers that were also enriched in the scRNA-seq cluster sc1 (Fig. S34) and that supported its annotation as “suppressed bract region” (Fig. 5C). This annotation was also supported by smFISH cluster c6 (suppressed bract) showing the highest contribution to sc1 among the different smFISH clusters (Data S12). The suppressed bract clusters c6 and sc1 also showed high correlations with the transition zone clusters c19 and sc8, respectively (Data S3 and S8), probably reflecting their shared function in repressing the leaf ridge.

The overlap between c6 and sc1 is likely incomplete. Cluster sc1 also showed a relatively high contribution of smFISH cluster c3 (Fig Data S12), suggesting that it may include cells from both the suppressed bract and ground tissue. Conversely, the c6 cluster includes epidermal cells (Fig. 4C-D), which were included in the “epidermal / suppressed bract” cluster sc15 in the scRNA-seq study (section 2.3).

#### 2.8. Boundary clusters

Among the genes analyzed in the smFISH experiment, *FZP* and *TCP24* showed the most specific expression domains (Figs. 4, S15, Data S2). *FZP* was expressed in the glume axilla (c14), while *TCP24* was expressed adjacent to *FZP* at the adaxial boundary region below each spikelet (c16, Fig. 4C-D and S15-16). In addition to *FZP,* the hexaploid wheat spatial expression data showed enriched expression in the adaxial boundary for the *WUSCHEL-RELATED HOMEOBOX9c* (*WOX9c*, Fig. S35A), *TCP22*, *LAX1*, and *DP1* (synonymous with *BAF1*) [21]. We validated this expression profiles in the Kronos imputed expression data (Fig. S27M, O-P).

Mutations in wheat *MULTI-FLORET SPIKELET 1* (*MFS1*, synonymous with *DUO* and *ERF130*) wheat have been previously associated with mild supernumerary spikelets and increased grain number per spike, highlighting its role in wheat spike development [77]. This motivated us to include *MFS1* in the smFISH experiment, which showed high levels of expression in clusters c14 and c16, and lower expression in other clusters (Fig. S35B-C). The high expression in c14 and c16 correlated with high *MFS1* expression in the scRNA-seq clusters sc9 and sc11 (Fig. S35D), prompting us to investigate a potential overlap among these clusters.

Three lines of evidence supported the annotation of single-cell cluster sc11 as “glume axilla” (Fig. 5C). First, glume axillary markers *FZP* and *WOX9c* were both preferentially expressed in sc11 (Fig. S36). Second, c14 (glume axilla) showed the highest %-contribution to sc11 among the smFISH clusters, (Data S12). Finally, RNA-seq results showed that sc11 was significantly enriched in genes with higher expression in the center of the spike (Data S16), mirroring the expression patterns of *FZP* and *WOX9c*.

We annotated the scRNA-seq cluster sc9 as “adaxial boundary” based on multiple shared markers with smFISH cluster c16. *TCP24* was preferentially expressed in sc9 and was also abundant in sc13 (Fig. S37), which includes the epidermal cell layer of the adaxial boundary domain (section 2.3). Other markers preferentially expressed in the adaxial domain, such as *TCP22, LAX1,* and *DP1* were also preferentially expressed in sc9 and/or sc13 (Fig. S37). The smFISH cluster c16 (adaxial boundary domain) %-contribution to sc9 was more than two-fold higher than the second highest (c9 central-base spikelet), which was also more than two-fold higher than the other smFISH cluster (Data S12).Taken together, these results support the annotation of sc9 as adaxial boundary domain.

#### 2.9. Cortex cells below developing spikelets

In the wheat spike, the cortex consists of a layer of cells situated between the vascular tissues and developing spikelets (Fig. 1). The parenchymatic cells in this region are expected to resemble those in the central ground tissue. As shown in Fig. 4C, the cell layer between the vasculature and the SM base includes ground tissue cells from the central cluster c3 as well as the innermost cells from clusters c6 (suppressed bract), c7 (glume – lemma), c9 (SM base-center), c16 (adaxial boundary) and c14 (glume axilla). Since cortical cells show no greening and are not meristematic, the scRNA-seq clusters including these cells are expected to show low enrichment in photosynthetic genes and reduced expression of *OSH1* or ribosomal protein-encoding genes.

Clusters sc2, sc4, and sc17 exhibited high correlations with the spatial clusters located proximal to the developing spikelets in hexaploid wheat (ED0, ED3, ED5, and ED11, Data S13-S15) [21]. In addition, we found relatively high %-contributions of the smFISH clusters c16 and c6 to sc2, from clusters c9 and c7 to sc4 and from cluster c16 to sc17 (Data S12). They also exhibited low expression levels of photosynthetic and meristematic genes (Data S16) and uniform expression along the spike based on the RNA-seq data. Taken together, these results suggest that sc2, sc4, and sc17 may correspond to cortical cells below the spikelets (Fig. 5C).

The annotation of sc2 as a cortex cluster was further supported by the preferential expression of *LAX1* and *DEPRESSED PALEA1* (*DP1*) (Fig. S38), two genes previously shown to be preferentially expressed in the adaxial boundary below the spikelets [78–81] and validated in the Kronos imputed data (Fig. S27O-P). Two additional genes, *KAN1* and *RL9*, expressed in the adaxial boundary region below the spikelet in hexaploid wheat [21], were also preferentially expressed in sc2 (Fig. S38), and their imputed expression is presented in Fig. S38H. Additional markers enriched in the correlated sc2 and sc13 clusters (Data S8) included *ABCG11* [82] and *SOT17* [83] (Data S9) and were validated in Fig. S38H.

A more central location of sc4, relative to sc2 and sc17, was supported by the higher %-contributions of the smFISH cluster c9 (central region below spikelet) to sc4 than to the other two spatial clusters (Data S12). This hypothesis was also supported by the 2.4 to 3.4-fold higher expression of *OSH1* in sc4 than in sc17 and sc2, respectively (Figs. S12, S39, Data S16). In addition, the cortex marker gene *NPF3.1* (also known as *NPF6.4-LIKE2* in maize [84]) was enriched in sc4 (Fig. S39F-G), suggesting that this cluster corresponds to cortical cells.

Overall, the scRNA-seq cluster sc17 showed lower correlations with the spatial clusters in hexaploid wheat than sc2, but they were still higher for the contiguous cortex clusters than for the rest of the spatial clusters (Data S13-15). The highest %-contribution from the smFISH clusters to sc17 was from c16 (adaxial boundary, adjacent to cluster, Data S12). We annotated sc17 as “tentative cortex region below spikelet lateral regions” due to the lack of validation with marker genes of known spatial distribution. Additional genes preferentially expressed in sc4 and sc17 are listed in Fig. S39 and Data S9.

#### 2.10. Meristem clusters

The RNA-seq clusters sc14 and sc20 showed preferential expression of meristematic genes *ULT1*, *AGL6* and *SEP3-1* (Fig. S40A-D) suggesting that they included meristematic cells. This hypothesis received additional support from the differential expression of the two homeologs of *ULT1.* The differences between *ULT-A1* and *ULT-B1* were evident in the bubble plot in Fig. S40G, in the UMAPS for the separate homeologs (Fig. S40H-I), and in the imputed expression profiles (Fig. S40J-K). In the bubble plots and UMAPS, *ULT-A1* showed high expression in meristematic clusters sc14 and sc20 (Fig. S40H) whereas *ULT-B1* did not (*ULT-B1* showed differential expression in the central clusters sc6 and sc10, Data S9). These differences were reflected in the imputed expression profiles, with *ULT-A1* expression concentrated in the early (c8) and floral meristems (c17 and c18) and *ULT-B1* expression concentrated in the central part of the spike (Fig. S40J-K). The correspondence between the spatial location and *ULT1* homeologs and their scRNA-seq cluster assignment in the UMAPS and bubble plots supported the conclusion that sc20 and sc14 include meristematic cells. More generally, the *ULT1* results suggest that the expression imputation method developed by Demesa-Arevalo et al. [70] in diploid barley, can be also used differentiate expression profiles between wheat close homeologs in polyploid species.

The presence of meristematic cells in sc20 and sc14 was also supported by high levels of expression of ribosomal genes (Data S16), intermediate expression levels of *OSH1* (Fig. S40G) and by the higher %-contributions of the smFISH cluster c8 to both sc20 (1.5% = 33 cells) and sc14 (5.3% = 114 cells) relative to other smFISH clusters (Data S12). Taken together these results support the annotation of sc14 and sc20 as meristematic clusters.

*AGL6* was detected in early meristems (c8) immediately before their transition to floral meristems (Fig. S40L), after which it became more abundant at the base of the FM (c17, Fig. S40L). This result is consistent with the role of *AGL6* role in paleae and lodicule development [12]. *SEP3-1* showed a similar profile but its expression started slightly later than *AGL6* and was detected in both the FM (c18) and its base (c17, Fig. S40M). Fig. S40N-P shows three additional genes differentially expressed in sc14 (Data S9). *SEP1-2* is preferentially expressed in lemmas and a few meristematic cells (*SEP1-2*, Fig. S40N and Fig. 4). The rice homolog of *SEP1-2*, *LHS1/OsMADS1*, controls floret meristem specification [85]. The second gene, *JLO*, encodes a LOB domain protein gene expressed in Arabidopsis at the base of the floral meristem (c17, Fig. S40O) [86]. Finally, *FCP1* was expressed in the floral meristem (c17 and c18) and c14 at the glume axilla (Fig. S40P). In rice, *FCP1* is a negative regulator of *WOX4* and of meristem maintenance [87]. Based on these results sc14 was annotated as “meristem proximal region”. Cluster sc20 was enriched in genes differentially expressed in the distal one-third of the spike (Data S16), so it was annotated as “meristem / enriched in spike tip”.

Many of the genes enriched in sc14 and sc20, also show some enrichment in epidermal clusters sc7 and sc16 (Fig. S40G, small arrows), supporting their previous annotation as “epidermis / meristem and lateral organs”. The imputed expression data showed enriched expression of *WOX3b* and *CLE33* in c8 including the epidermal cells (Fig. S27J-K). Additional genes preferentially expressed in sc14 and sc20 are listed in Data S9.

#### 2.11. Protoplasting affected clusters

The last two scRNA-seq clusters, sc3 and sc22 (smallest cluster with only 163 cells), showed evidence of being affected by the protoplasting. First, these clusters showed the highest proportion of preferentially expressed genes that were not detected in RNA-seq studies of Kronos developing spikes (this study and [68]). Approximately 19% of the preferentially expressed genes in sc3 and 28% in sc22, were not detected at the same stages in Kronos spikes under normal conditions (Data S16). The 235 genes detected only in the scRNA-seq were enriched in stress-related GO terms (Data S19), further supporting their induction by protoplasting. The preferentially expressed genes in sc3 and sc22 also showed a relatively high overlap with a list of genes previously annotated as potentially affected by protoplasting [84] (sc3= 11% and sc22= 13%, Data S16). The percentage of genes identified by these two criteria across the 23 single-cell cluster showed a high correlation (*R=* 0.81), indicating a similar underlying process. Cluster sc3 showed a high correlation with sc10 (*R=* 0.85, Data S8) and sc22 with cluster sc14 (*R=* 0.88, Data S8), suggesting that these clusters may contain cells related to those clusters but affected by protoplasting. Based on these results, we annotated sc3 and sc22 as “unknown”.

#### 2.12. Co-expression analyses

The generation of smFISH and scRNA-seq datasets from the same genotype, tissues, and developmental stages provided the opportunity to identify additional genes co-expressed in specific expression domains. We focused on four genes with specific expression domains ―*FZP* and *TCP24* in the boundary domains and *ULT1* and *AGL6* in the meristem― and calculated Pearson correlations between each of them and all other genes across all cells in the scRNA-seq dataset. The genes with *R* values exceeding four standard deviations above the mean (close to zero as expected) were selected as co-expressed genes for further analyses (Data S20-S24).

##### 2.12.1. Genes co-expressed with boundary genes FZP (c14) *and* TCP24 (c16)

The genes with the highest co-expression values for *FZP* and *TCP24* were their respective homeologs, serving as positive controls (Data S20-S21). We found 608 genes significantly co-expressed with *FZP* and 606 with *TCP24*, with 226 shared (988 total).

Among the shared co-expressed genes, we identified the well-studied gene *CUP-SHAPED COTYLEDON 3* (*CUC3*, Data S20-S21), a NAC-domain transcription factor involved in meristem / organ boundary specification in rice [88] and Arabidopsis [89]. *In situ* and scRNA-seq experiments in maize demonstrated that *CUC3* is expressed in both the *FZP* and *TCP24* domains [81, 84, 90], a result validated by our imputed expression data (Fig. S27N). The *MFS1* gene also showed a significant correlation with both *FZP* and *TCP24* (Data S20-S21) and was co-expressed with both genes in the smFISH and imputed expression data (Fig. S35B-C).

Among the genes showing significant co-expression with *FZP* and not with *TCP24*, we identified *WOX-9c* and *CLE33* (Data S20. Fig. S35). *WOX-9c* is a close paralog of *WOX7* [91, 92] with a similar expression profile as *FZP* (Fig. S35A). *CLE33* showed a similar expression profile as *WOX7* (Figs S16 and S18C), changing from expression in the early meristems (Fig. S27K) to an overlap with *FZP* in the more developed spikelets with FMs (Fig. S35E). In Arabidopsis, RNA interference of *WOX9* in reproductive meristems resulted in the conversion of flowers into inflorescence-like structures [93], resembling the conversion of spikelets into branches observed in *fzp* natural mutants in wheat and other grass species [49–52]. The rice loss-of function mutations in *WOX7* (*dwt1*) resulted in larger panicles, suggesting that these two related *WOX* genes have functional roles in inflorescence development [59].

*TCP24* showed high co-expression with *TCP22* (the close paralog of *TCP24*), *DP1*, and *LAX1* (Data S21), three genes previously confirmed to be expressed in the adaxial boundary domain in hexaploid wheat [21] and validated in our imputed expression data (Fig. S27M-P). *DP1* is an ortholog of the maize gene *BARREN STALK FASTIGIATE1* (*BAF1*, Data S21), which is expressed in a narrow domain adaxial to developing axillary meristems in maize [90]. Mutations in *BAF1* resulted in the absence of ears or their fusion to the main stalk, supporting its role in demarcating the boundary of the developing axillary meristem. *LAX1* is required for the initiation of aerial lateral meristems and is expressed in the adaxial boundary region of the spikelet meristems in both rice and maize (*BARREN STALK1*, *ZmBA1*) [78, 79]. A strong correlation was also observed between *TCP24* and *RAMOSA2* (*RA2*, Data S21), which is preferentially expressed in cluster sc9 in the imputed expression data (Fig. S35F). In maize, *RA2* is expressed in a group of cells that predict the position of the axillary meristem formation [94]. Additional genes co-expressed with *TCP24* are listed in Data S21. Based on the physical location of *TCP24* and its co-expressed wheat genes, together with the known functions of the maize homologs, we hypothesize that c16 cells contribute to delimiting the wheat SM and act as an important signaling center for spikelet development [95].

To investigate the interactions centered on *TCP24* and *FZP*, we generated a gene regulatory network using GENIE3 [96], focusing on the 988 genes co-expressed with *FZP* (Data S20) and/or *TCP24* (Data S21) in the scRNA-seq dataset. From the top-500 significant interactions, we constructed a network visualized in Cytoscape 3.10.3 (Fig. S42). The 95 genes included in this network were all transcription factors (Data S22) suggesting that *FZP* and *TCP24* are part of a broad regulatory network including many other transcription factors. The analysis revealed connections between the *TCP24* homeologs, which share multiple targets, including their closest paralog *TCP22*. *TCP24* also showed regulatory links with *LAX1* homologs and *RA2*, supporting their coordinated expression in the adaxial boundary (Fig. S42, Data S22). Similarly, the two *FZP* homeologs were connected and shared common links including *WOX9c*. Additionally, *TCP24* and *FZP* shared several downstream targets, such as the boundary regulator *CUC3*, suggesting a potential functional convergence (Fig. S42, Data S22). This regulatory network analysis provides a framework to investigate the genetic interactions that define the functions of these boundary domains.

##### 2.12.2. Genes co-expressed with meristem genes ULT1 *and* AGL6

The smFISH results for *ULT1* showed expression in the IM, SM and FM meristems as well as in the central spike region (Fig. 4F and S18C-D). However, the co-expression analysis revealed limited correlation between *ULT-A1* and *ULT-B1* homeologs (Data S23) supporting their functional differentiation described in section 2.10 and Fig. S40H-K. Interestingly, among the 100 genes showing the highest correlations with *ULT-A1*, 82 encoded ribosomal proteins (Data S23), suggesting a higher rate of protein production and rapid cell growth and division. This enrichment was not observed for *ULT-B1*, supporting the conclusion that *ULT-A1* is the homeolog predominantly expressed in meristematic cells. A heatmap based on the 45 ribosomal genes with the highest correlation values with *ULT-A1* revealed that cluster sc20 exhibited the highest expression level among the 23 single-cell clusters (Fig. S43). Cluster sc14 also showed relatively high levels of ribosomal genes, but lower than those in sc20.

Co-expression analysis of *AGL6* revealed significant correlations with floral homeotic genes *SEP3-1, SEP3-2*, *WAPO1*, *AG1*, and *PI1* (Data S24). Among the 100 genes showing the highest co-expression values with *AGL6*, 43 encode ribosomal proteins (Data S24). A similar enrichment of ribosomal protein encoding genes has been reported in a single-cell study of maize stem cells [66]. This co-expression studies also revealed genes with previously uncharacterized roles in inflorescence development (Data S23-S24). We explored the imputed expression of these genes and prioritized three that were expressed in the spike meristematic regions for further functional characterization: the remodeling genes *DDM1a* [97] (Fig. S41B) and *BTBN16* (Fig. S41C), and the auxin response gene *SHI1* [98] (Fig. S41D).

#### 2.13. scRNA-seq trajectory analysis

To summarize the relationships among the scRNA-seq clusters, we performed a trajectory analysis (Fig. S44) starting from the early meristematic cluster sc14. In the lower region of the graph, one branch of the trajectory connected sc14 with the vascular clusters sc18 and sc19 (and the cell division clusters), and the other branch with the epidermal clusters (Fig. S44). The early connection of sc14 with sc5 (epidermis unknown) and sc7 (epidermis meristem and lateral organ) suggested that these clusters may correspond to early stages of differentiating epidermis cells. The same trajectory later connected with sc15 (epidermis repressed bract) and sc13 (epidermis adaxial boundary).

In the opposite direction, sc14 was linked to two separate loops, each with multiple internal connecting trajectories. The first loop included the ground tissue clusters sc6, sc10 and sc12 and the transition zone sc8. The second interconnected loop included the cortex clusters sc2, sc4 and sc17, and the related repressed bract cluster sc1 and adaxial boundary sc9. The glume axilla cluster sc11 remained disconnected from the rest despite its proximity to cortex cluster sc4 and epidermal cluster sc13. Overall, the predicted trajectories aligned relatively well with the expected relationships among the different cell types derived from the meristematic cells in the wheat spike.

## 3. CONCLUSIONS

The use of the same germplasm, tissues, and developmental stages in our smFISH and scRNA-seq experiments allowed us to integrate the two datasets, which had positive synergistic effects. The precise spatial information provided by the smFISH experiment was useful for improving the annotation of the scRNA-seq clusters. However, the number of genes that can be included in smFISH experiments is limited. We show here that this problem can be mitigated by using the scRNA-seq data to impute the expression of a large number of genes into spatially anchored cells from a spatial transcriptomics experiment. These imputed expression profiles can be used to characterize the genes differentially expressed in each scRNA-seq cluster, thereby improving their annotation. This imputation method can be also used to prioritize genes for functional validation.

The method performed well both in diploid barley [70] and in tetraploid wheat, for which it correctly predicted the locations of epidermal genes even when no epidermal markers were included in the selected smFISH genes. The method also enabled us to distinguish expression profiles between closely related wheat homeologs, confirming that this method can be useful for studying expression in polyploid species with similar genomes.

The co-expression analyses, combined with smFISH and imputed expression profiles helped us identify wheat genes with expression domains similar to those of their orthologs from rice and maize, two crops with more extensive functional information than wheat. Comparative smFISH studies of inflorescence development across different grass species using common sets of genes can strengthen information bridges among crops. The analysis of co-expressed genes using gene networks provided another layer of information on gene relationships in a cell-type dependent manner. These localized networks can be useful for understanding the function of specific expression domains such as the glume axillae and the adaxial boundary domain. Finally, these tools can be used to identify genes with interesting expression profiles or interactions and prioritize them for functional characterization.

In addition to providing useful datasets and tools, this study offered interesting biological insights into the function of *LFY*. For example, the discovery of *LFY* expression in the stem intercalary meristem motivated us to study its role in stem development and lead to the discovery of a previously unknown role of *LFY* in stem development. The smFISH experiment also revealed consistent and previously unknown expression domains in the region proximal to the developing wheat spikelets, which likely correspond to the “signaling centers” proposed in maize to regulate SM development and determinacy [95].

In summary, the integrated smFISH and scRNA-seq dataset provides a dynamic view of gene expression during the early stages of wheat spike development with improved spatiotemporal resolution. All data and tools generated in this study are publicly available to empower future studies of wheat spike development and support comparative genomic studies across grass species.

## 4. MATERIALS AND METHODS

### 4.1. Spatial Expression

#### 4.1.1. Sample preparation

Developing wheat spikes from the tetraploid wheat Kronos were collected at three developmental stages: early transition from the vegetative to the reproductive stage (W1.5), late double ridge stage before the development of glume primordia (W2.5), and floret primordia stage, when the central spikelets initiated the transition to the floral meristem (W3.5). The different tissue sections used for W1.5 and W2.5 correspond to different plants, but the W3.5 sections A1-1 and A2-1 correspond to the same spike and the sections A1-2 and A2-2 to a single spike from a different plant. Although the two sections from the same spike contain different cells, they were averaged and treated as technical replicates in the statistical analyses. After tissue collection, samples were fixed in 4% paraformaldehyde (PFA), dehydrated, and embedded in paraffin. Longitudinal sections (10 µm-thick) from the central plane of the developing spikes were positioned within the capture areas of the slides. The slides were dried overnight at 37°C, followed by a 10-minute bake at 50°C to enhance section adhesion. The sections were then deparaffinized, permeabilized, and refixed. After complete dehydration, they were mounted using SlowFade-Gold Antifade reagent, covered with a thin glass coverslip, and shipped on dry ice to Resolve BioSciences (Monheim am Rhein, Germany) for analysis.

Spatial transcriptomics analyses were performed using Molecular Cartography^TM^ and a 100-plex combinatorial single-molecule fluorescence in-situ hybridization. Upon arrival, tissue sections were washed twice in 1x PBS for two minutes, followed by one-minute washing in 50% and 70% ethanol at room temperature. Ethanol was removed by aspiration, and DST1 buffer was added followed by tissue priming and hybridization. Tissues were primed for 30 minutes at 37°C, followed by hybridization for 48 h using probes specific to the target genes. The following day, samples were washed to remove excess probes and fluorescently tagged in a two-step color development process. Regions of interest were imaged as described below, and fluorescent signals were removed during decolorization. Color development, imaging, and decolorization were repeated across multiple cycles to generate a unique combinatorial code for each target gene, derived from raw images as described below.

#### 4.1.2. Probe design

We used our published RNA-seq data [68] to select genes expressed at appropriate levels at the three selected spike developmental stages and prioritized those that showed dynamic changes across these stages. We also prioritized MADS-box genes and other transcription factors previously shown to be involved in inflorescence or floral development in other plant species. Finally, we also included marker genes to facilitate cluster annotations.

For each of the selected tetraploid wheat genes, we selected the homeolog with higher expression based on published RNA-seq data for Kronos spike development [68]. We also provided Resolve Biosciences with the sequences of the lower-expressing homeolog to exclude them during probe specificity evaluations conducted against all the coding sequences of the wheat genome (Ref Seq v1.1, Data S1). Therefore, although probes were designed based on the sequence of the highest-expressing homeolog, they are not genome-specific and can detect both homeologs for each gene. Probes were compared with all the wheat expressed genes, and those that could amplify other genes were discarded. If no specific probes could be identified for a gene, it was excluded from the experiment.

Probes for the 99 genes were designed using Resolve Biosciences’ proprietary design algorithm and gene annotations from the Chinese Spring RefSeqv1.1 database. In brief, probe design was conducted at the gene level. For each targeted gene, all full-length protein-coding transcript sequences from the ENSEMBL database were utilized as design targets. To identify potential off-target sites, searches were confined to the coding regions. Each target sequence underwent a single scan for all k-mers, favoring regions with rare k-mers as seeds for full probe design. A probe candidate was generated by extending a seed sequence until reaching a certain target stability. Sequences known to cause problems experimentally were eliminated. Following these initial screens, each retained probe candidate was aligned with the background transcriptome, and probes with stable off-target hits were discarded. Specific probes were then scored based on the number of on-target matches (isoforms), giving preference to principal isoforms over others. A bonus was added if the binding site was within the protein-coding region. From the pool of accepted probes, the final set was selected from the highest-scoring probes.

#### 4.1.3. Imaging

Samples were imaged by Resolve Biosciences using a Zeiss Celldiscoverer 7 (CD7) with a 50× Plan Apochromat water immersion objective having an NA of 1.2 and a 0.5× magnification changer, resulting in a final magnification of 25×. Standard CD7 LED excitation light sources, filters, and dichroic mirrors were used in conjunction with customized emission filters optimized for detecting specific signals. The excitation time per image was fixed at 1,000 ms for each channel, 20 ms for DAPI, and 1 ms for Calcofluor White. A z-stack was captured at each region with a z-slice spacing according to the Nyquist-Shannon sampling theorem. A custom CD7 CMOS camera (Zeiss Axiocam Mono 712, 3.45 μm pixel size) was employed. For each region, a z-stack was acquired for each of two fluorescent colors in each imaging round. A total of 8 imaging rounds were performed for each position, resulting in 16 z-stacks per region. The entirely automated imaging process per round, which included water immersion generation and precise relocation of regions for imaging in all three dimensions, was implemented through a custom Python script utilizing the scripting API of the Zeiss ZEN software (Open application development). Imaging for the cell wall-specific stain, Calcofluor White, was conducted after the primary 8-round imaging.

#### 4.1.4. Spot segmentation and preprocessing

The algorithms for spot segmentation were written in Java and are based on the ImageJ library functionalities. Only the iterative closest point algorithm is written in C++ based on the libpointmatcher library (https://github.com/ethz-asl/libpointmatcher).

In the initial image processing steps, all images underwent background fluorescence correction. The allowed number of maxima, representing the brightest points in the images, was determined based on the slice’s area and an empirically optimized factor. Each image slice was processed independently. Maxima without neighboring counterparts in adjacent slices (referred to as z-groups) were discarded. An iterative filtering process adjusted thresholds for factors like absolute brightness, local background, and background of the periphery, all based on 3D image volume. Only maxima remaining in z-groups of at least 2 were retained. These maxima, with the highest absolute brightness within their z-groups, were documented as 3D-point cloud features.

For signal segmentation and decoding, the raw data images were aligned using transformation matrices derived from the feature point clouds. An iterative closest point cloud algorithm minimized error between point clouds. Point clouds from each round were aligned to the round one reference, and the corresponding point clouds were stored for downstream use. Aligned images were used to create pixel profiles with 16 values, representing two color channels across eight imaging rounds. Profiles were filtered for variance from zero, normalized by total profile brightness.

Matched pixel profiles were assigned IDs, and neighboring pixels with the same ID formed groups. These pixel groups underwent further filtering, considering group size, adjacent pixel count, and dimensions with a size of two pixels. Local 3D-maxima in groups were identified as potential transcript locations. Maxima were filtered based on expectations from raw data images. The remaining maxima were evaluated by fitting to corresponding codes, and they were documented as transcripts. Specificity, assessed by the ratio of experiment-matching signals to non-experiment-matching signals, was used to estimate false positives. This comprehensive process enabled precise image analysis and transcript identification. Final image analysis was performed in ImageJ using the Polylux tool plugin from Resolve BioSciences to examine specific Molecular Cartography^TM^ signals. Images, programs, and gene expression results are deposited in Zenodo [99].

#### 4.1.5. Cell segmentation, clustering, and cell trajectories

We identified cell boundaries from the DAPI-stained images using the cell segmentation program QuPath [20]. To account for different cell sizes and shapes, we defined different regions in each image, applying region-specific parameter sets. Across all regions we used the parameters pixel size = 0, background radius = 8 μm, intensity threshold = 5, and opening by reconstruction with the “include cell nucleus” parameter disabled. For regular, well-defined cells, “split by shape” was activated, and the cell expansion was set to 7 μm, while for irregular, elongated cells, “split by shape” was turned off, and the cell expansion was set to 5 μm. The cell walls, visualized with Calcofluor White in the same sections, served as controls. The six images were merged into a single composite image and converted into a format suitable for clustering using ImageJ.

Hybridization signals per cell and clustering were calculated using the Resolve Biosciences “Recognize” software with the following parameters: Min cell = 5, Min Features = 1, Min transcripts = 1, Dimensions = 20, Resolution = 0.75, and Selected genes = 99. “Recognize” utilizes Seurat v4 [100] as the backend, with the following methods: NormalizeData (normalization.method= “LogNormalize”, scale.factor= 1), FindVariableFeatures, ScaleData, RunPCA, FindNeighbors (dims= 1:20), FindClusters (resolution= 0.75), RunUMAP (dims= 1:20, n.components= 3, and FindAllMarkers. The R script to replicate the analysis using Seurat v5 can be found in Zenodo [99]. The average log2 fold changes of marker genes were generated locally using Seurat v5 [69], which implements a pseudocount at the group level instead of the cell level as in Seurat v4.

Cells from the three developmental stages and two replications were clustered together. Clusters are represented with the same color across stages and replicates. To facilitate visualization of the individual cell clusters, we also present them separately in Fig. S5. The Seurat object was deposited in Zenodo [99].

To find subclusters within scRNA-seq vasculature, we used the Seurat function “*FindSubCluster*” with parameter ‘*resolution* = 0.01*, <u>graph.name</u>* = “RNA_snn”, *algorithm* = 2. We then used the Seurat function “*FindMarkers*” to find differentially expressed genes between the subclusters including xylem and phloem markers.

We reconstructed cellular trajectories using the computer program Monocle 3 [101]. The 21 cell clusters from the spatial analysis and the 23 cell clusters from the scRNA-seq were leveraged to constrain two separate trajectory inferences. The initial cluster object was transferred into a Monocle SingleCellExperiment object using the as.cell_data_set function. Dimensionality reduction was carried out using the UMAP embeddings previously computed with RunUMAP(dims=1:30) into the Monocle object. Then function “cluster_cells” was applied using the default random seed and UMAP as the reduction method to preserve cluster consistency, followed by “learn_graph” to construct the principal graph, and “order_cells” to infer pseudotime trajectories, specifying meristematic cluster c8 as the root cluster in the spatial trajectory and sc14 in the scRNA-seq trajectory.

### 4.2. RNA-seq of different parts of the developing wheat spike

Kronos spikes at the glume primordia stage (W3.0) were sectioned into three parts (Fig. S24) and placed into 500 µl of DNA/RNA Shield solution in 1.5-ml Eppendorf tubes. Approximately 20 to 25 apices were used per replicate, and three replicates were collected for each section.

Samples in DNA/RNA Shield were stored at 4 °C prior to RNA extraction.

We followed the published protocol by Marina Millán Blánquez for RNA extraction (dx.doi.org/10.17504/protocols.io.36wgq7kj3vk5/v1) with minor modifications. Briefly, 400 µl of DNA/RNA Shield solution was removed, and the tissues were homogenized with plastic pestles in the remaining 100 µl of solution. RNAs were then extracted using TRIzol reagent, cleaned, and concentrated with the RNA Clean & Concentrator-5 kit from Zymo Research. RNA samples were shipped to Novogene for quality control, library preparation, and sequencing on the NovaSeq X Plus (PE150) platform.

Data have been deposited in bioproject number PRJNA1217025. The three replicates from each region were clustered together in a principal component analysis, indicating high reproducibility among biological replicates (Fig. S24). Of the 49,786 detected genes, 13,024 exhibited significant differences among regions in Tukey tests (Data S11). For each region we calculated the average TMM per region and performed Tukey tests. The genes showing significant differences and the corresponding Tukey tests are listed in Data S11. Scripts for RNA-seq mapping, principal component plot, and Tukey tests were deposited in Zenodo [99].

### 4.3. Single-cell RNA sequencing

#### 4.3.1. Protoplast preparation and 10x Genomics library construction and sequencing

We used a cell wall digestion method optimized to preserve meristem cells in maize inflorescence [66]. Briefly, the inflorescence spike was finely dissected (Fig. 1A) under a microscope and digested in a MOPS buffer-based enzyme mix (10 mM MOPS, pH 7.5; 10 mM L-Arginine HCl, pH 7.5; 0.65 M mannitol; 1.5% Cellulase RS; 1.5% Cellulase R10; 1.0% Macerozyme; 1.0% hemicellulose; 0.1% BSA; 5 mM β-mercaptoethanol; 1 mM CaCl₂) for 2 hours. Before adding CaCl₂, β-mercaptoethanol, and BSA, the solution was heated to 55°C for 10 minutes.

The digested cells and enzyme mix were passed through a 30 µm pluriStrainer (pluriSelect, Catalog No. 43-50030-50) for filtration. The filtered protoplasts were pelleted by centrifugation at 350g for 2 minutes at 4°C and washed with wash buffer (10 mM MOPS, pH 7.5; 10 mM L-Arginine HCl, pH 7.5; 0.65 M mannitol). The protoplasts were pelleted again by centrifugation at 350g for 2 minutes at 4°C. The clean protoplast pellets were gently resuspended in wash buffer, and viability was checked using trypan blue. High-viability protoplasts (≥ 70%) were immediately loaded into a 10x Genomics Chromium System using Next GEM Single Cell 3’ Kit v3.1 chemistry. The scRNA-seq libraries were sequenced on the Element Bio AVITI platform using the 10x Genomics recommended sequencing format, with 122 cycles used for Read 2 (Data S7).

#### 4.3.2. Single-cell RNA-seq analysis and clustering

Single-cell RNA-seq reads were mapped to the Kronos genome assembly v1.1. (annotation v1.0) [67] (https://doi.org/10.5281/zenodo.11106422) using STARsolo [102, 103]. Since scRNA-seq reads were generated from the 3’ end of the transcripts and not all Kronos genes have good annotations of 3’ UTR regions, we extended the 3′ end of all gene models by 500 bp.

We used the Seurat V5 pipeline [69] to cluster the cells. Briefly, genes expressed in at least three cells were retained. All cells from both stages were filtered to include those with 1,500 to 25,000 expressed genes and 8,000 to 300,000 unique molecular identifiers (UMIs).

Reads from individual samples were log-normalized, and 2,000 highly variable genes were selected for downstream analysis. The integrated matrix was scaled using the ‘ScaleData’ function. Principal component analysis (PCA) was performed with the ‘RunPCA’ function. Further integrative analysis of all W2.5 and W3.5 stage samples was conducted using the ‘IntegrateLayers’ function with the anchor-based RPCA integration method. The top 35 principal components (PCs) were used. A resolution of 1 was applied with the ‘FindClusters’ function to identify clusters. Cell cycle clusters were identified (Fig. S23), and the remaining clusters were re-normalized, scaled, integrated using the RPCA method, and re-clustered. The top 35 PCs were used to generate the final clusters, at a resolution of 1 (Fig. 5B-D). Scripts used for mapping reads and clustering analysis and the final Seurat object files were deposited in Zenodo [99].

#### 4.3.3. Correlations among cell clusters

Normalized expression data for the 99 genes in the smFISH study were used to calculate Pearson correlation coefficients between pairs of spatial clusters (Data S3). Normalized expression data for the 200 genes used in the spatial transcriptomics study in hexaploid wheat were kindly provided by Dr. Cristobal Uauy (John Innes Centre, UK) [21]. The hexaploid wheat data was provided separately for the four different stages included in their study, but we only used the three early stages (late double ridge, lemma primordia and terminal spikelet) that overlapped with the stages of spike development used in our scRNAseq study.

The average normalized gene expression across all cells for each single-cell cluster were obtained with the “AverageExpression” function in Seurat v5 in R after joining the six replications (three for each stage) of the scRNA-seq data. For this analysis, we used the final integrated data after removing the initial cell-cycle clusters and filtered the average normalized data per cluster using two criteria. First, we selected genes with coefficients of variation higher than 0.5 to eliminate genes with very similar expression levels across all clusters. We also eliminated genes with a total sum across clusters <0.1 or >20 to minimize the effect of genes with very high or very low expression levels (and multiple 0s). Using these criteria, we selected 14,903 informative genes that were used to calculate Pearson correlation coefficients between all possible pairs of single-cell clusters using their average normalized expression across all common genes (Data S8).

To calculate the correlations between the single-cell clusters and the spatial transcriptomics clusters from hexaploid wheat, we first identified the overlapping genes for each comparison. The coding regions of wheat homeologs are on average 97.5% identical, so most of the probes used in the wheat spatial transcriptomics experiment are expected to hybridize with the different homeologs. To make the single-cell data comparable, we summed the normalized expression of the three homeologs in hexaploid wheat, but separate correlation matrices were generated for the late double ridge (Data S13), lemma primordia (Data S14) and the terminal spikelet stage (Data S15).

Clustering of the correlation matrices was done by the function “heatmap” from R package [104] with default parameters (clustering_distance_rows = “euclidean”, clustering_distance_cols = “euclidean”, clustering_method = “complete”). Scripts used to extract cluster averages and to perform the correlation analyses are deposited in Zenodo [99].

#### 4.3.4. Imputation of scRNA-seq gene expression into spatial transcriptomics cells

To integrate the smFISH and scRNA-seq data without losing spatial information, we imputed the expression levels of all genes from the five nearest scRNA-seq cells into each cell from the spatial transcriptomic experiment using a recently published approach [70]. Briefly, we used SEURAT v5 [100] to calculate the cosine similarity (1 - cosine distance, varying from 0 for most dissimilar to 1 for the most similar) among the smFISH and scRNA-seq cells based on the expression of the 91 genes present in both studies (normalized counts per million). We identified the five nearest scRNA-seq cell neighbors to each smFISH-labelled cell and calculated a weighted average expression for each gene, using the cosine similarity as weight. The imputed gene expression was then incorporated into the smFISH-labelled cells.

We integrated the smFISH and scRNA-seq for each stage (W2.5 and W3.5) separately and then combined the imputed data. We imputed 70,878 genes for W2.5 and 72,475 for W3.5, resulting in 74,464 for the combined analysis. We also developed a website for visualization of the combined data (https://dubcovskylab.ucdavis.edu/imputed-genes). The scripts for data imputation, the final imputed data, and the source data of the website are included in the Zenodo depository (folder imputed_data) [99].

Using a similar procedure as in the imputation of scRNA-seq gene expression into spatial transcriptomics cells, after the integration, we found the five nearest smFISH cells for each scRNA-seq cell in W2.5 and W3.5 separately with one additional filtering: the nearest cell should have cosine similarity > 0.3 to exclude very different cells and at the same time keep enough number of cells for following analysis. We then assigned the scRNA-seq cell the most frequent smFISH cluster number within the 5 neighbors to the scRNA-seq cell. If there were no common smFISH cluster numbers (all 5 cluster numbers are unique), then this scRNA-seq cell had no smFISH cluster projection. After the projection, we calculated the frequency matrix between scRNA-seq original clusters and the projected smFISH clusters. The count matrix was normalized by the total number of cells used for each smFISH cluster (the frequency counts / total number of cells used in that smFISH cluster) to normalize for differences of cluster sizes.

#### 4.3.5. Annotation of scRNA-seq cell clusters

To annotate the scRNA-seq clusters, we used markers known to be expressed in specific cell types and/or previously associated with known functions in other species (Data S1) and validated their spatial expression in Kronos developing spikes using the imputation method described above. We supplemented this information with the correlations calculated between the single-cell and spatial expression clusters identified in the published spatial transcriptomics study in hexaploid wheat [21] (Data S13-15), and by the imputed expression of multiple differentially expressed genes in each scRNA-seq cluster using the methods described in the previous section.

Additional information to annotate the scRNA-seq clusters was extracted from the genes preferentially expressed in each cluster (Data S9). These genes were selected based on their low abundance in other clusters (pct.2<0.10) relative to their abundance in the studied cluster (pct.1>0.10), adjusted *P* values <0.001, and a greater than 2-fold difference in expression levels (log2Fold>1, Data S9). Clusters sc1 and sc22 had few enriched genes, so we relaxed the filtering parameters (average log2FC>1, pct.1 no filtering, pct.2<0.2, adjusted *P* value <0.01) (Data S9). We subsequently determined the pathways enriched in these preferentially expressed genes using KEGG analyses (Kyoto Encyclopedia of Genes and Genomes, Data S10) through the DAVID webtool (https://davidbioinformatics.nih.gov/tools.jsp) [105]. We converted the Kronos v1.0 gene IDs into Chinese Spring annotation v1.1 gene IDs, uploaded the CS gene IDs as “ENSEMBL_GENE_ID” (step 2: Select Identifier) and as “Gene List” (step 3: List Type), used the “*Triticum aestivum*” as background species, and started the analysis using “Functional Annotation Tool”. In addition, we examined the average log2Fold expression across clusters for a subset of 41 photosynthetic genes to identify clusters associated with photosynthetic functions, and for 45 ribosomal protein-encoding genes to identify cells with high protein production, typically associated with elevated cell division rates. These results were visualized using heat maps.

To incorporate additional spatial information into our cluster annotation, we used the RNA-seq data generated from the base, middle and top third of the spike described in section 4.2. (henceforth described as spike base, middle and tip). To determine if a cell cluster was enriched in any of these three spike regions, we identified genes with significant differences in Tukey tests among the preferentially expressed genes for each cluster and quantified the number of genes expressed at higher levels in each region. To determine if the distribution across the three regions was significantly different from a random uniform distribution, we performed chi-square tests for each cluster (Data S16).

#### 4.3.6. Detection of clusters affected by the protoplasting process

The protoplasting process may trigger a stress response in some cells, potentially influencing the clustering process. We employed two independent approaches to assess the potential effects of protoplasting on gene expression in the different clusters. First, we used a list of 713 maize genes previously identified as potentially affected by the protoplasting process [84], identified the closest Kronos homologs, and determined the percentage of preferentially expressed genes in each cluster overlapping with this list (totaling 215 unique genes). The second approach, which identified 235 unique genes, involved the identification of preferentially expressed genes in each single-cell cluster that were not present in two RNA-seq data from Kronos developing spikes (this study and [68]). A GO-term analysis of the unique genes identified in both analyses revealed enrichment in stress-related annotations, particularly among the genes not detected in the two RNA-seq datasets (Data S19). These results suggest that many of these genes may have been induced by the protoplasting process and are typically not expressed under normal conditions in the developing wheat spike. The percentages of genes identified by these two independent methods across clusters were highly correlated (*R*= 0.813, *P*< 0.0001) suggesting a similar underlying phenomenon. Both methods identified clusters sc3 and sc22 as having the highest risk of being affected by the protoplasting process (Data S16).

#### 4.3.7. scRNA-seq co-expression analysis

Using the cell x gene normalized count matrix, we calculated Pearson correlation coefficients (*R*) between genes of interest with known spatial distribution (*AGL6, FZP, TCP24*, *ULT-A1*) and all other detected genes in the scRNA-seq across all cells (26,009 cells. We focused on co-expressed genes with correlation coefficients higher than the average plus four standard deviations. The script used for co-expression analysis was deposited in Zenodo [99].

#### 4.3.8. Network analysis

To better understand the interactions between *TCP24* and *FZP*, we performed a gene regulatory network analysis using the program GENIE3 [96]. The analysis included the 988 genes co-expressed with *FZP* (Data S20) and/or *TCP24* (Data S21) in the scRNA-seq dataset. We extracted the top-ranked 500 links to generate and visualize the network using Cytoscape 3.10.3 (https://cytoscape.org/). In Data S22, we summarized the first neighbors of *TCP-A24*, *TCP-B24*, *FZP-A*, and *FZP-B* and identified shared and unique transcription factors among them.

#### 4.3.9. Generation of loss-of-function mutants of FZP and SPL14 using CRISPR-Cas9

To validate the specific functions of *FZP* and *SPL14,* we used CRISPR-Cas9 to edit both homeologs of each target gene in the tetraploid wheat variety Kronos. We designed two guide RNAs for *FZP* (FZP-gRNA-1 and FZP-gRNA-2, Table S1) and one for *SPL14* (SPL14-gRNA, Table S1). Guide RNAs for each gene were cloned into a binary vector (Addgene Plasmid #160393) containing the *Cas9* gene and a *GRF4-GIF1* chimera, which enhances wheat regeneration efficiency during *Agrobacterium*-mediated transformation [106]. Transgenic plants were generated at the UC Davis Plant Transformation Facility using methods described previously [106]. Independent transgenic plants were screened for mutations using next-generation sequencing (NGS) as previously described [107] using primers described in Table S1. We identified eight independent *T_0_* lines with mutations in both *FZP* homeologs and further confirmed the edits by Sanger sequencing using primers listed in Table S1. For *SPL14*, we obtained 10 individual *T₀* transgenic plants. This line was backcrossed with wildtype Kronos to segregate out the *Cas9* transgene, and we identified one line in the *T₄* generation with successful edits in both the A and B genomes and without the transgene. We developed CAPS markers to track the *spl14* mutations across generations using primers described in Table S1.

## ACKNOWLEDGEMENTS

The authors thank Chaozhong Zhang and Qiujie Liu (UC Davis) for their help with the selection of the genes included in the smFISH experiment, Samik Bhattacharya from Resolve Biosciences for his continuous help with the spatial transcriptomics project, Cristobal Uauy from the John Innes Centre UK for his advice with the analysis of the spatial transcriptomics data and early access to his spatial transcriptomics data, Rüdiger Simon and Björn Usadel for their help with the expression imputation method, and Ksenia Krasileva from UC Berkeley for early access to the Kronos genome. JD acknowledges funding from the Howard Hughes Medical institute and the United States Department of Agriculture (USDA) – National Institute of Food and Agriculture (NIFA) grants 2022-68013-36439 (WheatCAP) and 2022-67013-36209. X.X acknowledges USDA-NIFA HATCH project (CA-D-PLB-2850-H) and UC Davis new faculty start-up funding.

## AUTHORS CONTRIBUTIONS

XX, HL, and JD conceived the project. Experimental work was performed by HL, XX, FP, KL, CT, and YL. JZ, GB, HL, CL, XX and JD performed data analyses. JD and XX obtained funding and directed the project. HL, XX and JD generated the first draft, and all authors contributed to the revision of the manuscript. XX, CL and JD provided supervision.

## DATA AVAILABILITY

Data and scripts used in the different analyses are available in Supplemental Data and in Zenodo https://doi.org/10.5281/zenodo.14872904 [99]. The ImageJ version with the Polylux tool plugin, the high-resolution images for the six sections used in the paper (Calcofluor White Figs. S3, S4 and DAPI), the hybridization results for the 99 genes for each image, and instructions for download and installation are also available at our web site (https://dubcovskylab.ucdavis.edu/content/spatial-transcriptomic) and in Zenodo [99]. We also developed a web tool to visualize individual cell clusters and heatmaps of the 99 genes in the six sections https://dubcovskylab.ucdavis.edu/JD99-wheat-spike-smFISH and a similar tool to visualize the imputed genes https://dubcovskylab.ucdavis.edu/imputed-genes. These tools can be also downloaded from the same Zenodo link and run on individual computers. The raw RNA-seq data of the three regions of the Kronos spike at W3.0 were deposited in NCBI’s Sequence Read Archive (SRA) under BioProject number PRJNA1217025. The processed data, scripts used for mapping and feature counts, and a Readme file with instructions are deposited in Zenodo [99].

The scRNA-seq data were deposited in NCBI’s Sequence Read Archive (SRA) under BioProject accession number PRJNA1199502. The final single-cell clusters can be accessed via the Shiny App (https://dubcovskylab.ucdavis.edu/content/kronos-integrated-clusters). Additionally, a 3D-UMAP (https://dubcovskylab.ucdavis.edu/content/kronos-scrna-3d-clusters) was generated to visualize cluster distributions. The Kronos genome assembly v1.1 (annotation v1.0) is available in Zenodo https://doi.org/10.5281/zenodo.11106422. Processed single-cell data, Seurat objects and R scripts are also available in Zenodo https://doi.org/10.5281/zenodo.14872904.

Data and scripts for cluster correlations, the Seurat object produced by Resolve BioSciences “Recognize” online tool, and the R script to replicate the online analysis locally can be also found in Zenodo [99].

