## Supplemental Table S1 for "Spatial and single-cell expression analyses reveal complex expression domains in early wheat spike development"

**Supplemental Table S1**. Guide RNA sequences and primers used in the functional validation of *SPL14* and *FZP*.

| Guide/Primer Name | Sequence | Purpose |
|---|---|---|
| FZP-gRNA-1 | GCTCTCCATGAAGGGCGCGC | Crispr gene editing |
| FZP-gRNA-2 | TTCCTCGCGCCGTTCCACGC | Crispr gene editing |
| FZP_Geno_NGS-F1 | TCCTCTGTCACGGAAGCGCTCTCACTCCTTCTAGCTTCTCC | NGS |
| FZP_Geno_NGS-R1 | TTTAGCCTCCCCACCGACTCGAAGGTGCCGAGCCAG | NGS |
| FZP_Geno_NGS-F2 | TCCTCTGTCACGGAAGCGAGCCCGGGCGCTTCCT | NGS |
| FZP_Geno_NGS-R2 | TTTAGCCTCCCCACCGACCGCCGTGATCCGCGGCA | NGS |
| FZP-Sang-F2A | CCAACCTCACTTCACTTC | Sanger Sequencing |
| FZP-Sang-R2A | TACGGCATGGCCGAGGACGCG | Sanger sequencing |
| FZP-Sang-F2B | CCAGCATCACTTCAGTTG | Sanger sequencing |
| FZP-Sang-R2B | TACGGCATGGTGGAGGAC | Sanger sequencing |
| SPL14-gRNA | GCTTCTGCCAGCAGTGCAGC | Crispr gene editing |
| SPL14-NGS-F | TCCTCTGTCACGGAAGCGCACAAGGTGTGCTCCATG | NGS |
| SPL14-NGS-R | TTTAGCCTCCCCACCGACAGAATACGTCAAGAAATTG | NGS |
| SPL14A-F | GTGTGCTCCATGCACACCAAGGag | CAPS marker |
| SPL14A_PvuII_R | GAAGGTGACCAAGTTCATGCTGGTGGGAGGAGAAATGGTGACCaGCT | CAPS marker |
| SPL14B-F | GCGCCAAGCAATACCATTC | CAPS marker |
| SPL14B-PvuII-R | GAAGGTGACCAAGTTCATGCTGGGGAGGAGCAATGGTGACCaGCT | CAPS marker |
