## Supplemental Figures for "Spatial and single-cell expression analyses reveal complex expression domains in early wheat spike development"

**TABLE OF CONTENTS**

**1. Spatial expression and cell segmentation**
Fig. S1 Page 3
Fig. S2 Page 4
Fig. S3 Page 5
Fig. S4 Page 6
Fig. S5 Page 7
Fig. S6 Page 10
Fig. S7 Page 11
**1.1. The basal region below the spike**
Fig. S8 Page 12
Fig. S9 Page 13
Fig. S10 Page 14
**1.2. The transition zone between leaf and spikelet meristems**
Fig. S11 Page 15
**1.3. The central region of the spike**
Fig. S12 Page 16
Fig. S13 Page 18
**1.4. The region proximal to the spikelet meristem (SM) and *FZP* validation**
Fig. S14 Page 19
Fig. S15 Page 20
Fig. S16 Page 21
Fig. S17 Page 22
**1.5. Spikelet meristem (SM) and floret meristem (FM) regions**
Fig. S18 Page 23
**1.6. Changes within the inflorescence meristem (IM) and *SPL14* validation**
Fig. S19 Page 24
Fig. S20 Page 25
Fig. S21 Page 26
**1.7. Pseudotime trajectories**
Fig. S22 Page 27
**2. Comparison initial and final scRNA-seq clustering**
Fig. S23 Page 28
**2.1. RNA-seq of bottom, middle, and top of spike at W3.0**
Fig. S24 Page 29
**2.2 Cell cycle enriched clusters**
Fig. S25 Page 30
**2.3. Epidermal clusters**
Fig. S26 Page 31
Fig. S27 Page 33
**2.4. Vascular clusters**
Fig. S28 Page 35
Fig. S29 Page 38

**2.5. Central spike clusters**
Fig. S30 Page 39
Fig. S31 Page 41
Fig. S32 Page 42
**2.6. Transition zone between leaves and spike**
Fig. S33 Page 43
**2.7. Suppressed bract cluster**
Fig. S34 Page 45
**2.8. Boundary clusters**
Fig. S35 Page 46
Fig. S36 Page 47
Fig. S37 Page 48
**2.9. Cortex cells below developing spikelets**
Fig. S38 Page 49
Fig. S39 Page 51
**2.10. Meristem clusters**
Fig. S40 Page 52
Fig. S41 Page 55
**2.12. Co-expression analysis**
Fig. S42 Page 56
Fig. S43 Page 57
**2.13. scRNA-seq trajectory analysis**
Fig. S44 Page 58

**Fig. S1. Microscopy images of Kronos developing spikes. A, B** Stereo microscope images. **C, D** scanning electron microscope images. **A** Shoot apical meristem (SAM) producing leaf primordia. **B** Early stage of spike development showing initial meristem elongation and a lateral meristem before its differentiation into upper and lower ridges. IM= inflorescence meristem (red) and LM= lateral meristem (blue) **C** Double ridge stage (W2.5). SM= spikelet meristem (yellow) and repressed lower ridge (also known as leaf-ridge, violet). **D** Floret primordia stage (W3.5). Spikelet primordia are formed in a distichous organization. The glumes are indicated in green, the lemmas in blue, the floret meristem (FM) in pink, and the SM in yellow. In this spike the IM has already transitioned to a terminal spikelet showing two glume primordia, one lemma primordium with an incipient FM, and the terminal SM. Note the rotated orientation of the terminal spikelet relative to the lateral spikelets, with the glume and lemma primordia originating in the same orientation as previous SMs. **E** Putative section plane at W3.5 explaining the absence of SM cells adjacent to the large FM in the central spikelets of Fig. 1. The large FM may displace the SMs outside the section plane.

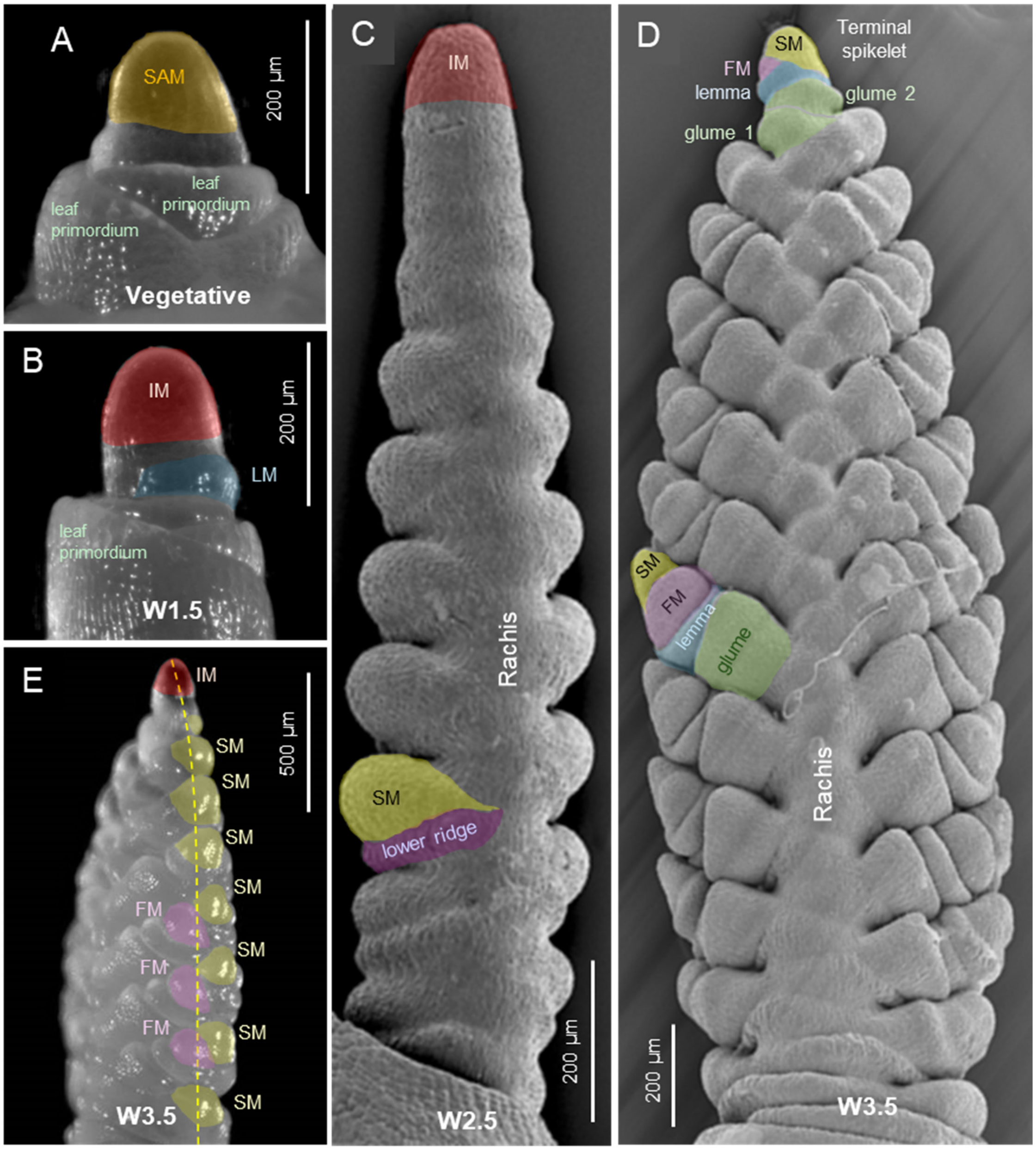

**Fig. S2. Additional sections showing cell segmentation and clustering.** Cell clusters based on the expression of 99 genes studied by smFISH at three different stages of wheat spike development. **A** W1.5, initial transition from vegetative to reproductive stage (section D1-2, 4123 cells,). **B** W2.5, late double ridge (section B2-4, 7,491 cells). **C** W3.5, floret primordia stage (section A2-2, 11,742 cells). Same colors represent same cell clusters in all sections in Fig. 1 and S2. The inset in panel **B** is a vegetative bud at the base of the section (truncated to fit the three figures in the page). IM= inflorescence meristem, SM= spikelet meristem and FM= floret meristem.

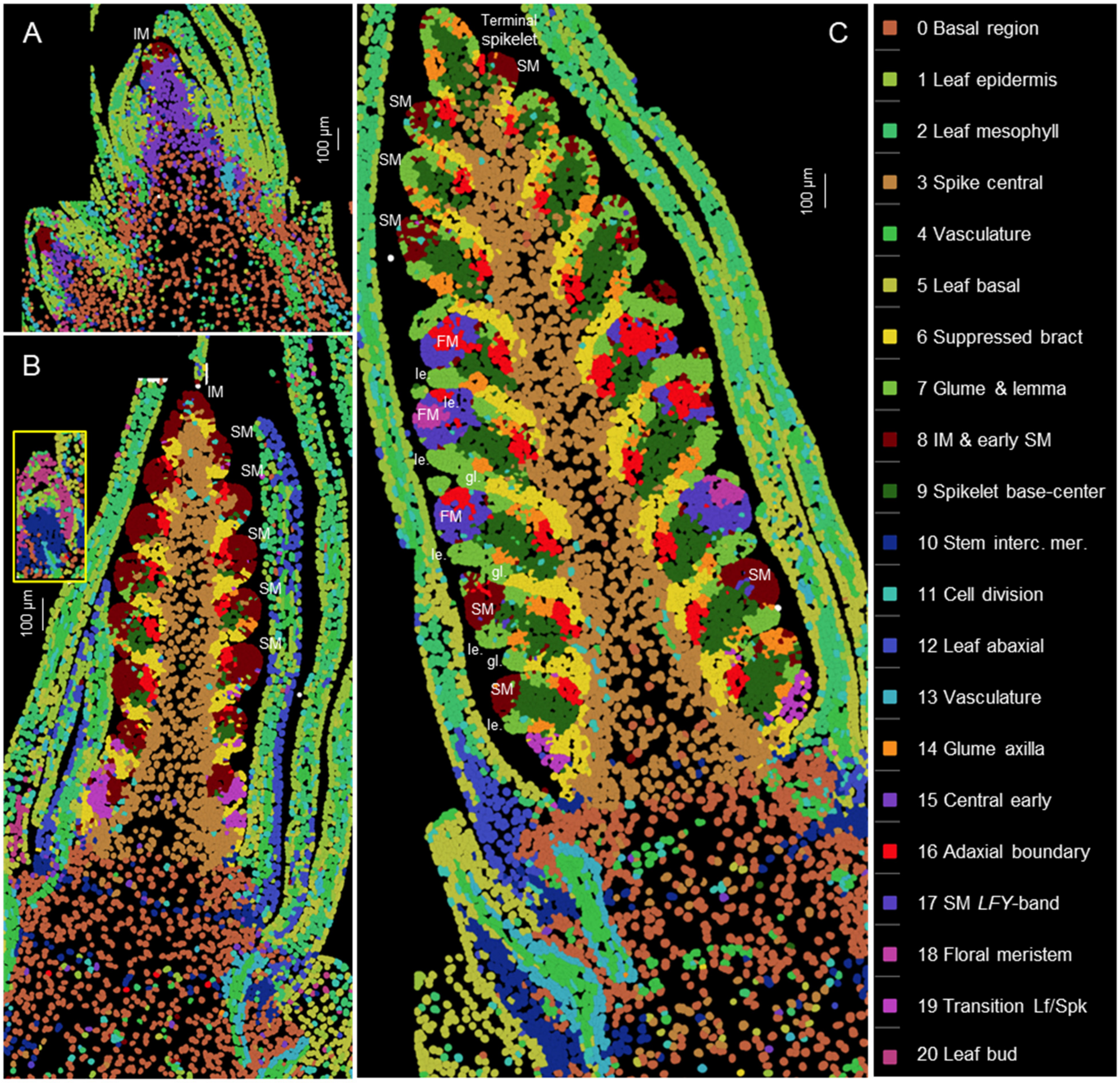

**Fig. S3. Calcofluor-white images of cell segmentation sections presented in Fig. 1. A** W1.5, initial transition from vegetative to reproductive stage. **B** W2.5, late double ridge. **C** W3.5, floret primordia stage.

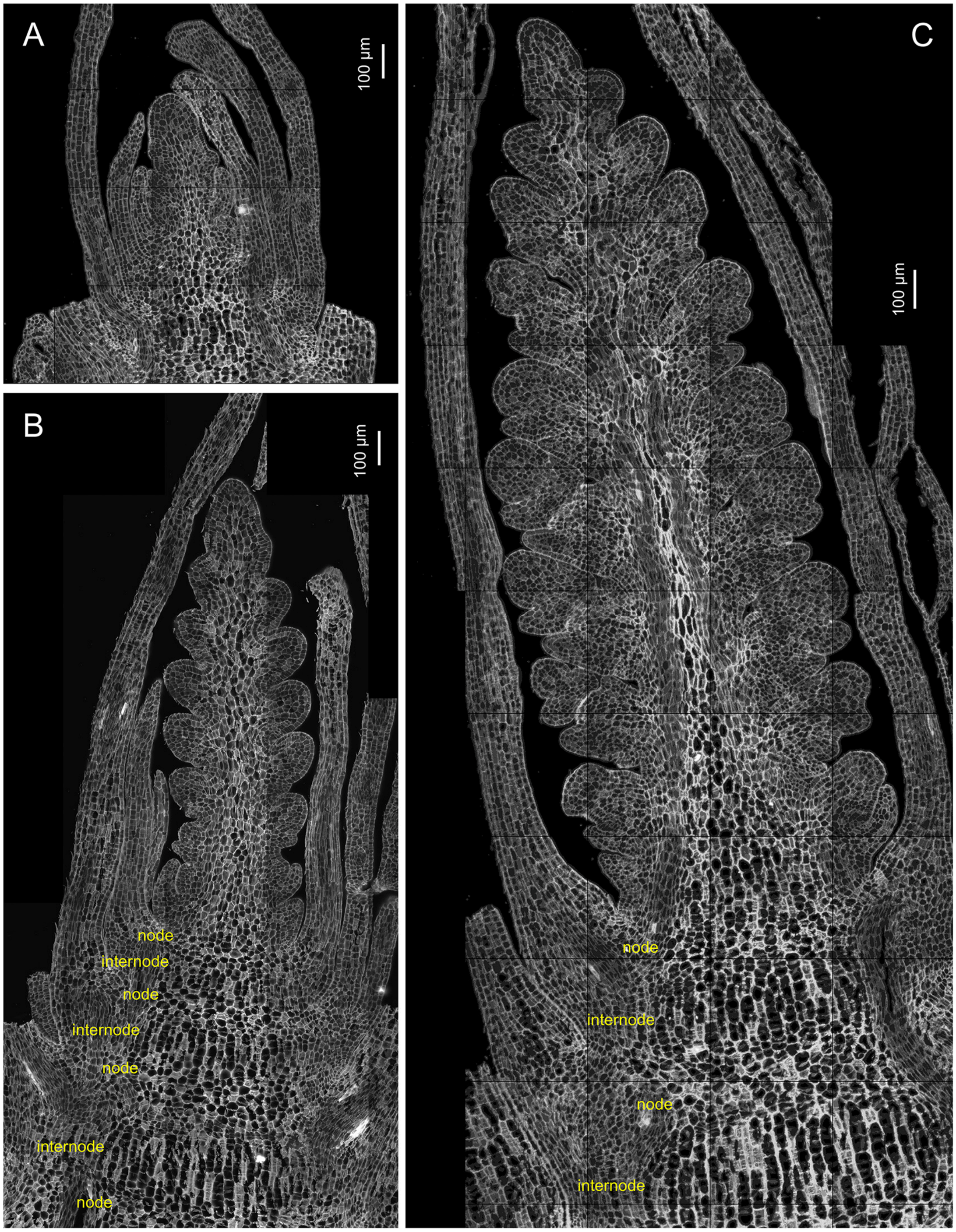

**Fig. S4. Calcofluor-white images of cell segmentation sections in Fig. S2. A** W1.5, initial transition from vegetative to reproductive stage. **B** W2.5, late double ridge. **C** W3.5, floret primordia stage. The inset in panel B is a vegetative bud at the base (left) of the section, which was truncated to fit the three figures in the page.

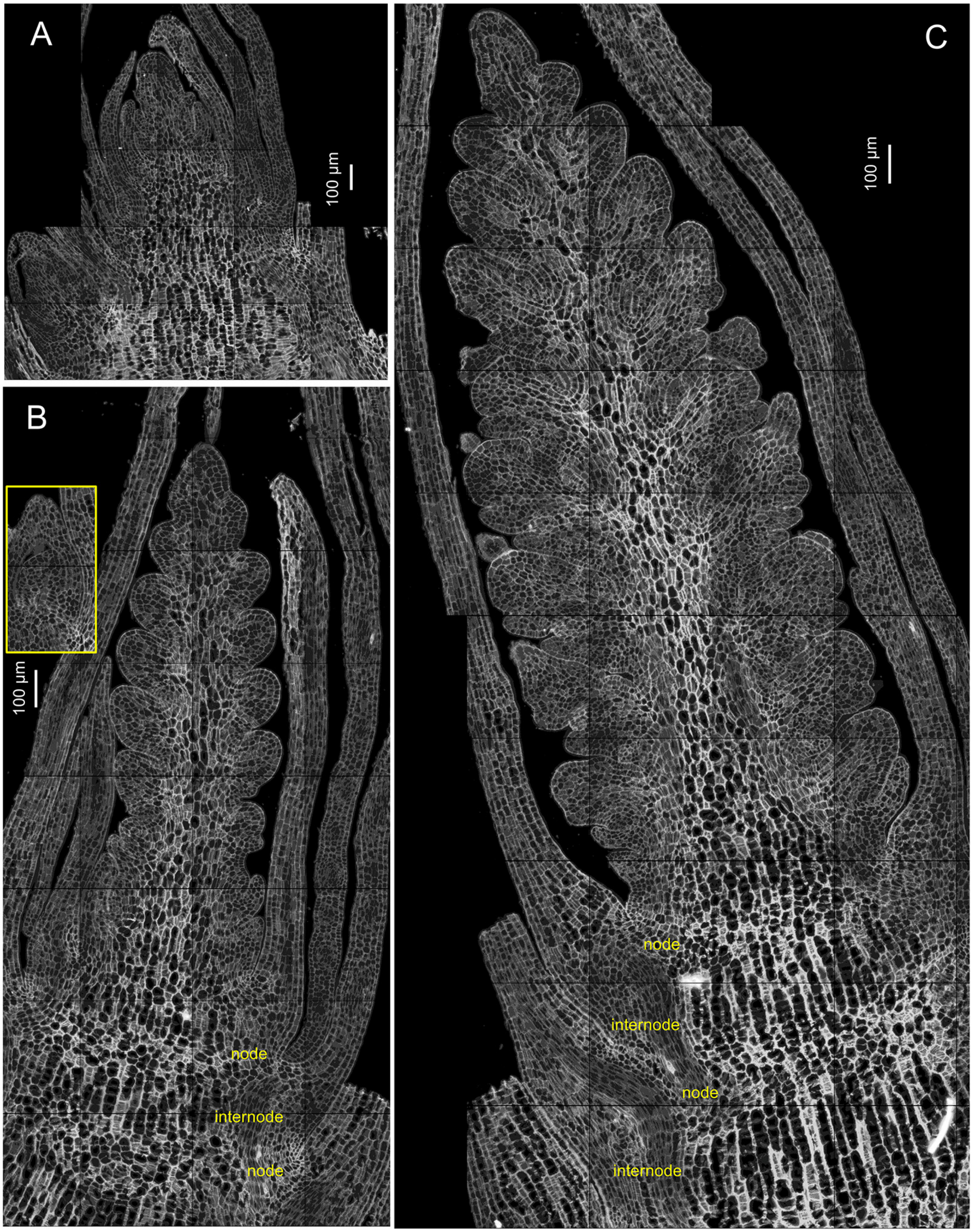

**Fig. S5. Individual cell clusters identified at three spike development stages** (W3.5, W2.5 and W1.5). Cell clusters are numbered from 0 to 20 based on the expression of 99 genes studied by smFISH. Cells from all sections were clustered together. Cluster numbers are followed by the preferentially expressed genes in white and the preferentially not expressed genes in red.

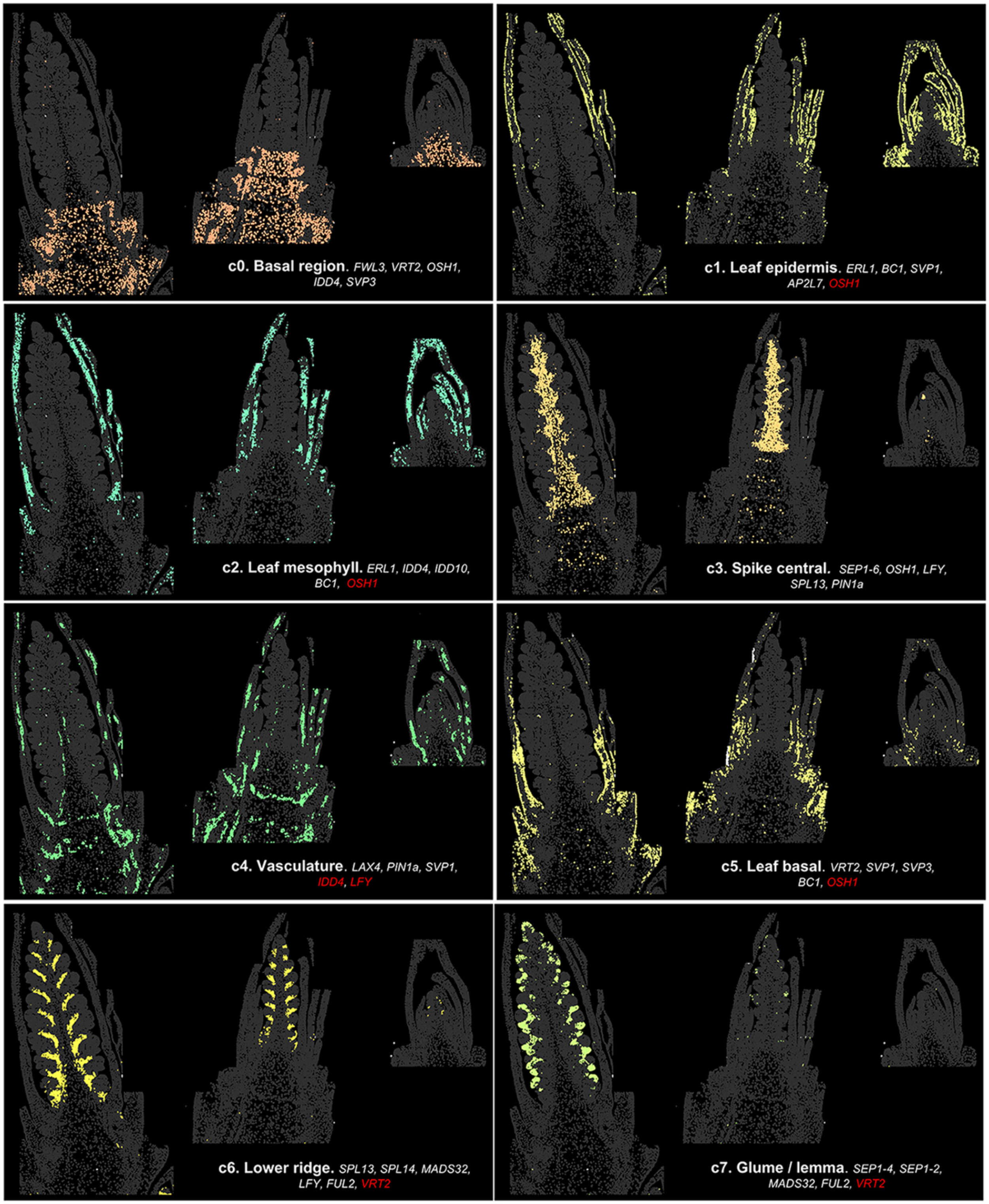

**Fig. S5. Continuation**

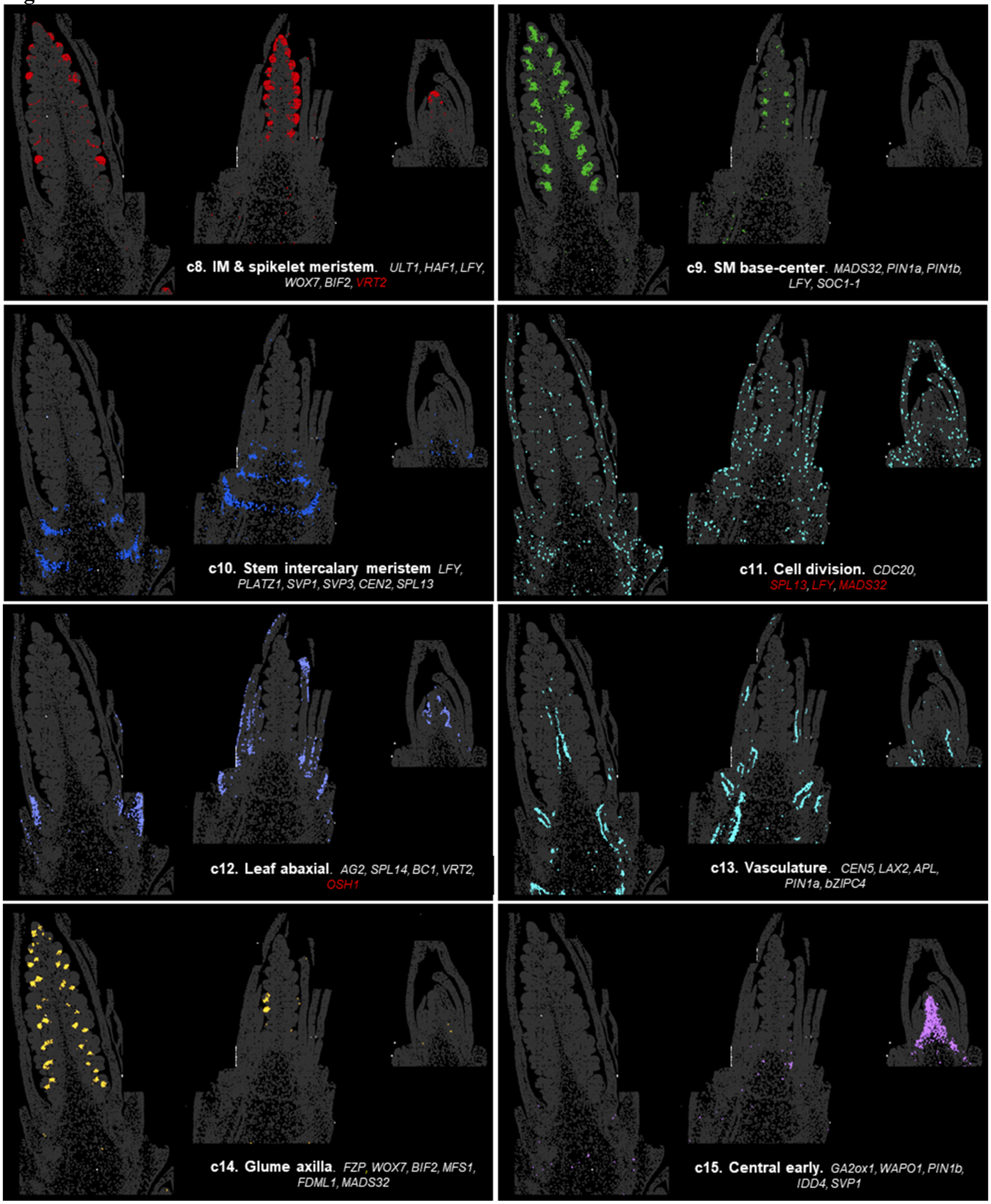

**Fig. S5. Continuation**

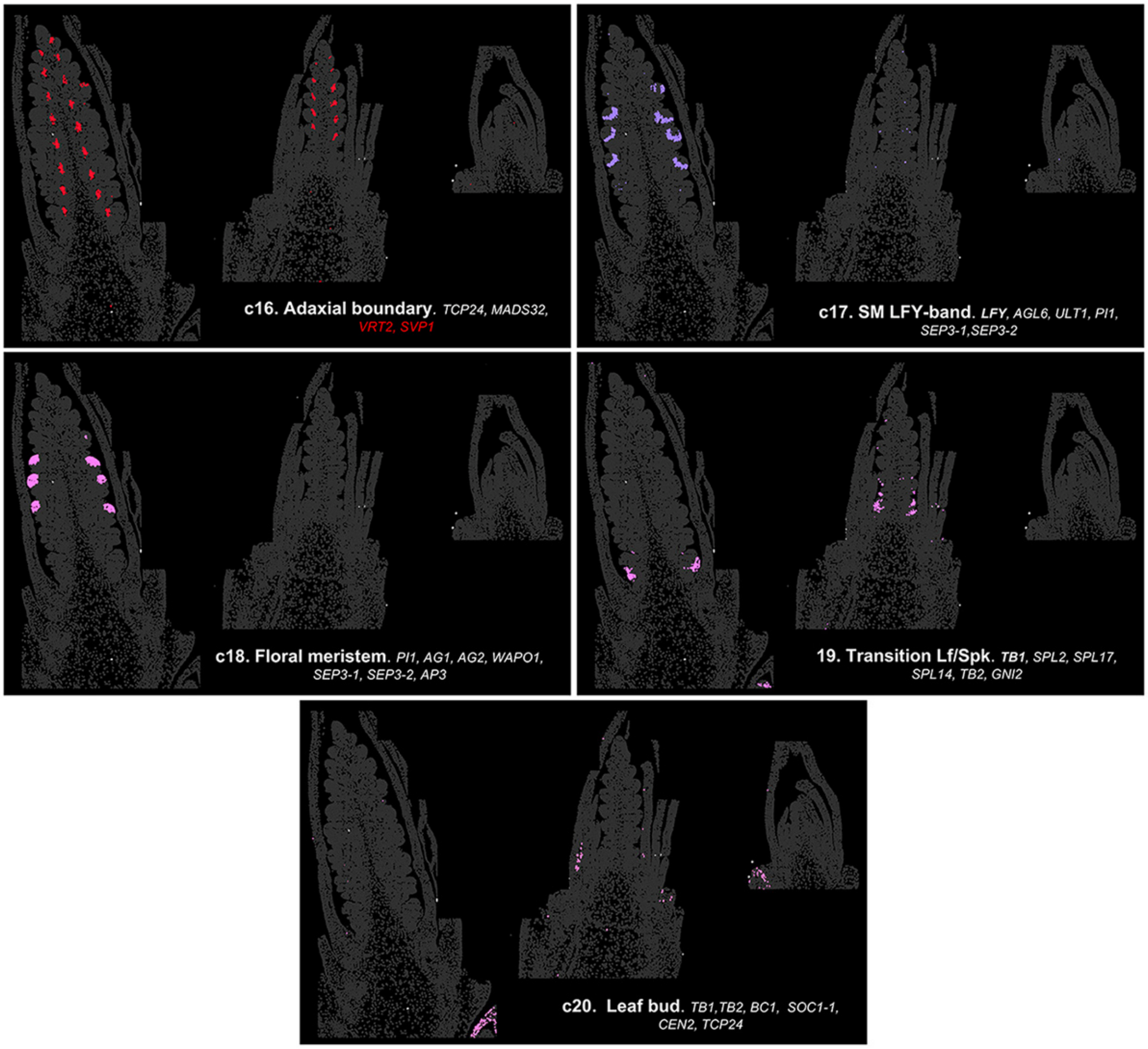

**Fig. S6. Validation of smFISH profiles.** Comparison of Molecular Cartography expression profiles and *in situ* hybridization profiles previously obtained in our laboratory.

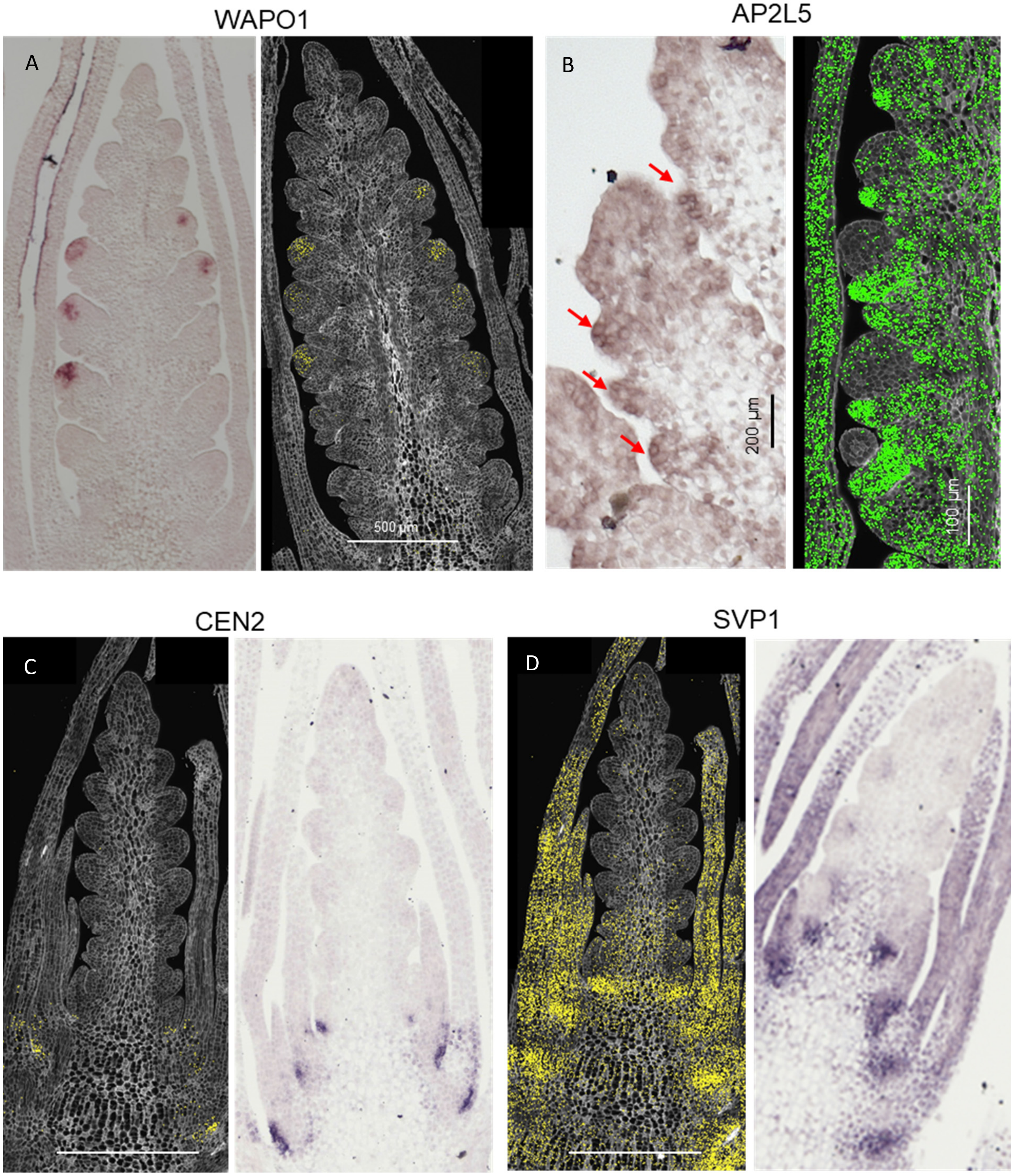

**Fig. S7. Validation of smFISH profiles.** Comparison of Molecular Cartography expression profiles and MERFISH results in hexaploid wheat spikes at similar stages.

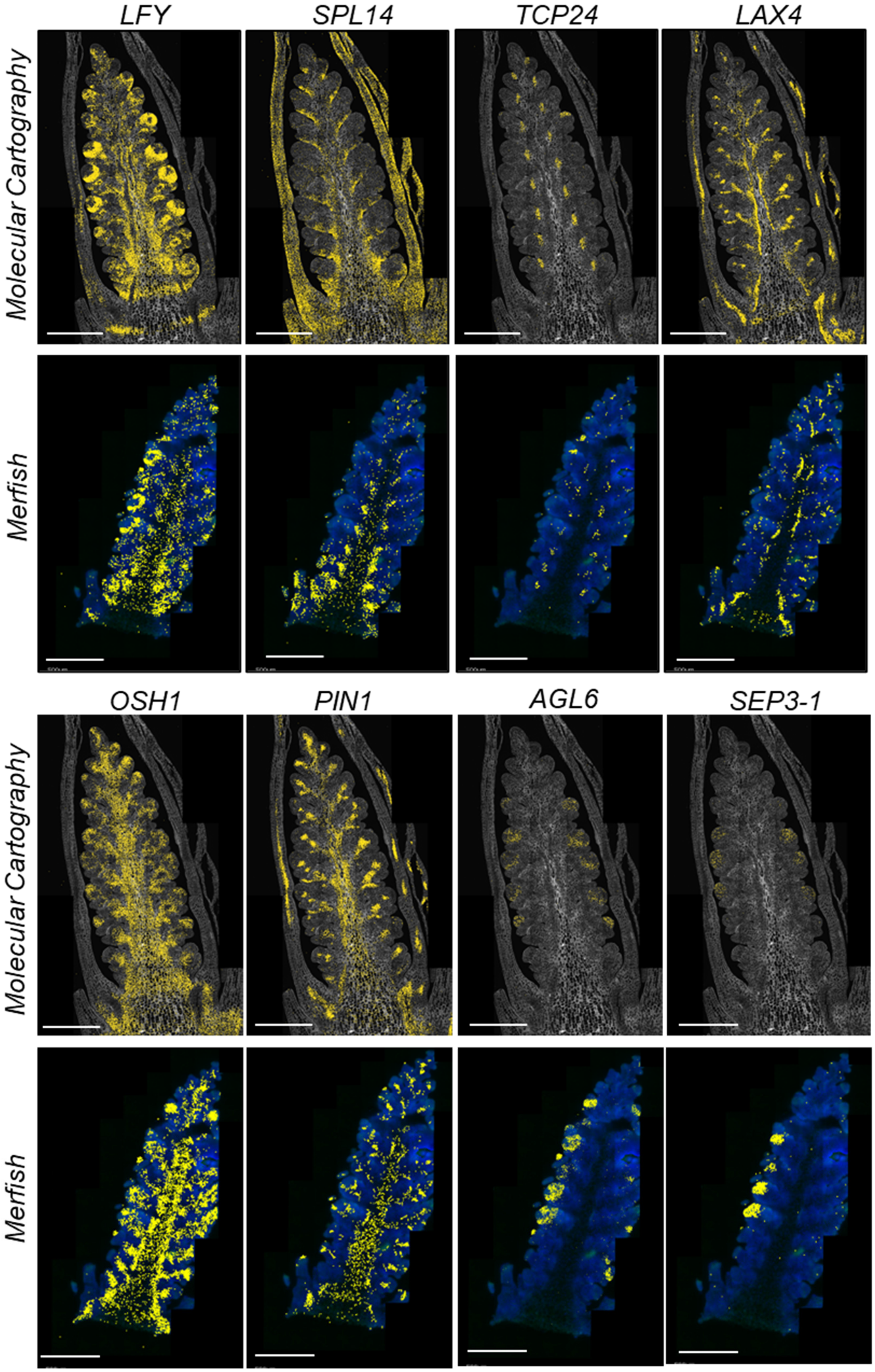

**Fig. S8. Basal region below the spike.** Cell walls are stained with calcofluor-white. **A-B** W3.5. **C-D** W2.5. **A** *LFY, PLATZ1, ULT1*, and *SVP1*. **B** *LFY* and *NAL1*. **C** *LFY, PLATZ1, ULT1* and vascular markers *LAX4* and *APL*. **D** *LFY, NAL1*, and *SVP1*.

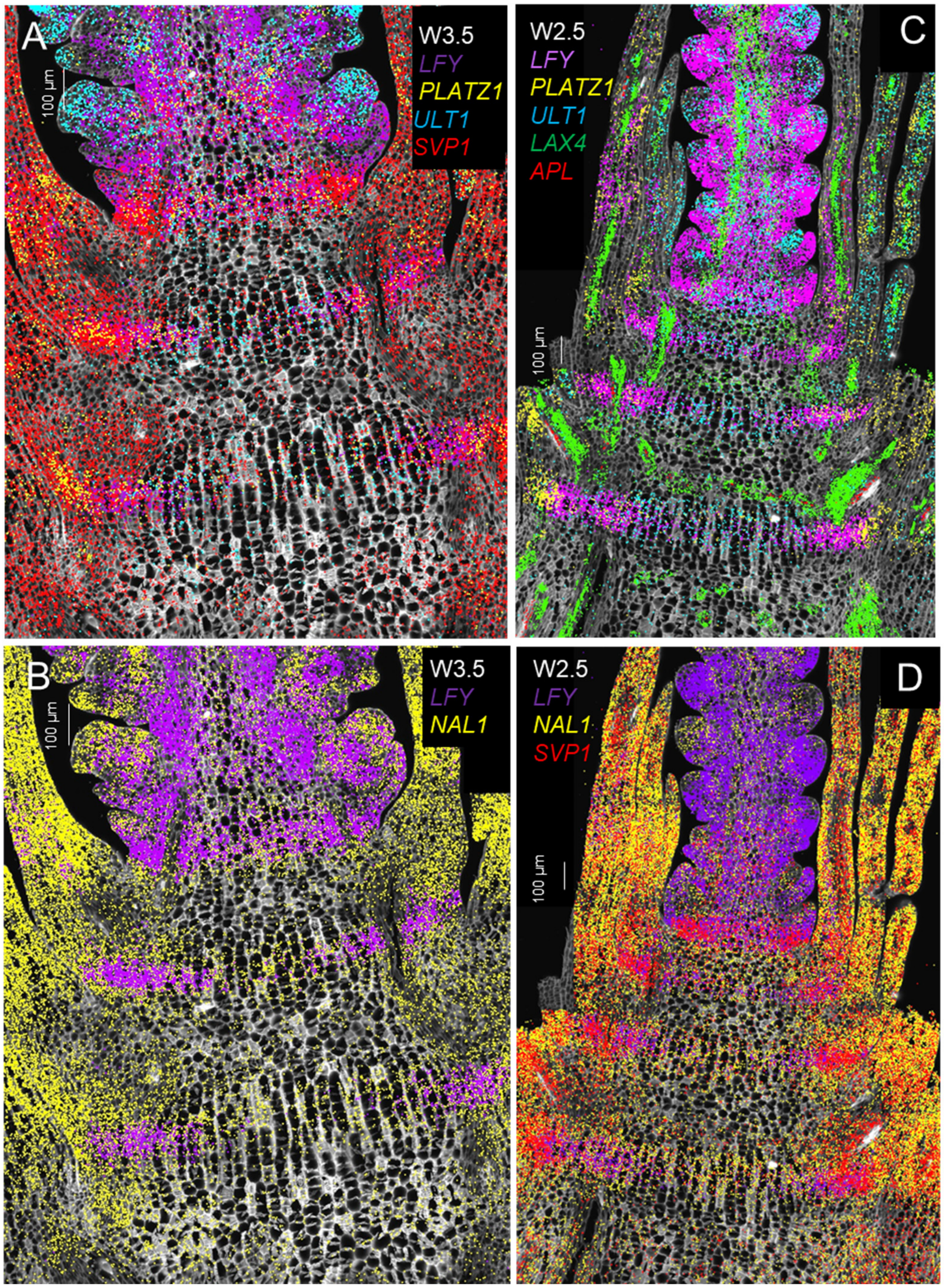

**Fig. S9. Functional validation of *LFY* role in the intercalary meristem.** Effect of combined loss-of-function mutations in both *LFY* homeologs on **A** node diameter in mm, **B** internode diameter in mm, and **C** hollow or solid stem internodes. Internodes and nodes are numbered starting from the peduncle downward. Bars are averages of 5 wildtype-plants and 12 *lfy* mutant plants. Error bars are S.E.M. *= $P<$ 0.05, **= $P<$ 0.01, and ***= $P<$ 0.001 using two-tailed *t*-Tests. Sections were stained with Toluidine Blue. Raw data and statistical analyses are available in Data S4.

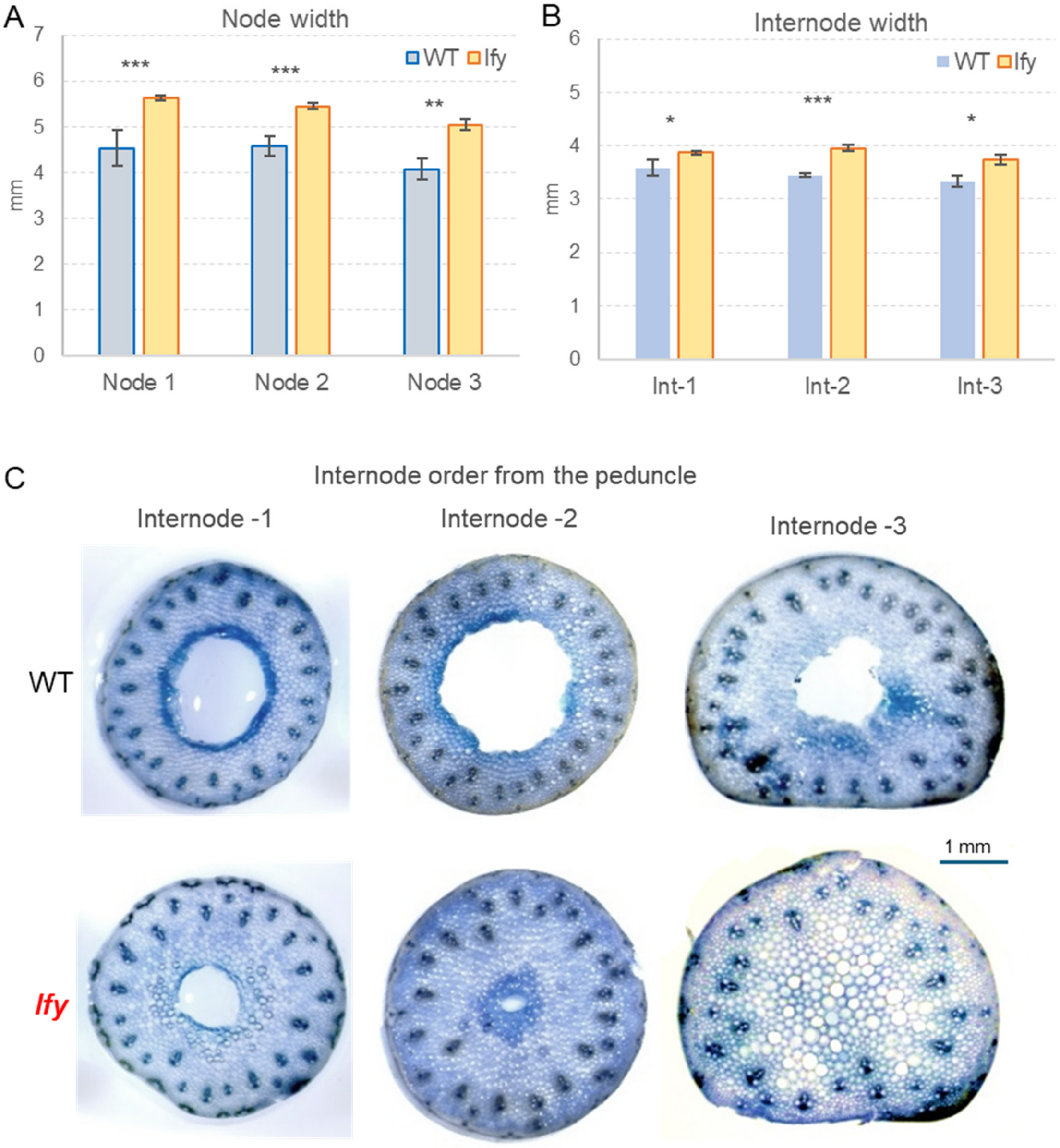

**Fig. S10. Expression of genes involved in the biosynthesis (*GA20ox1* and *GA20ox2*) and degradation (*GA2ox1*) of gibberellin in the basal region below spike.** Cell from cluster c10 (stem intercalary meristem) are marked in blue. **A** W1.5. **B** W2.5. **C** W3.5. *GA20ox1* (yellow), *GA20ox2* (green), and *GA2ox1* (red).

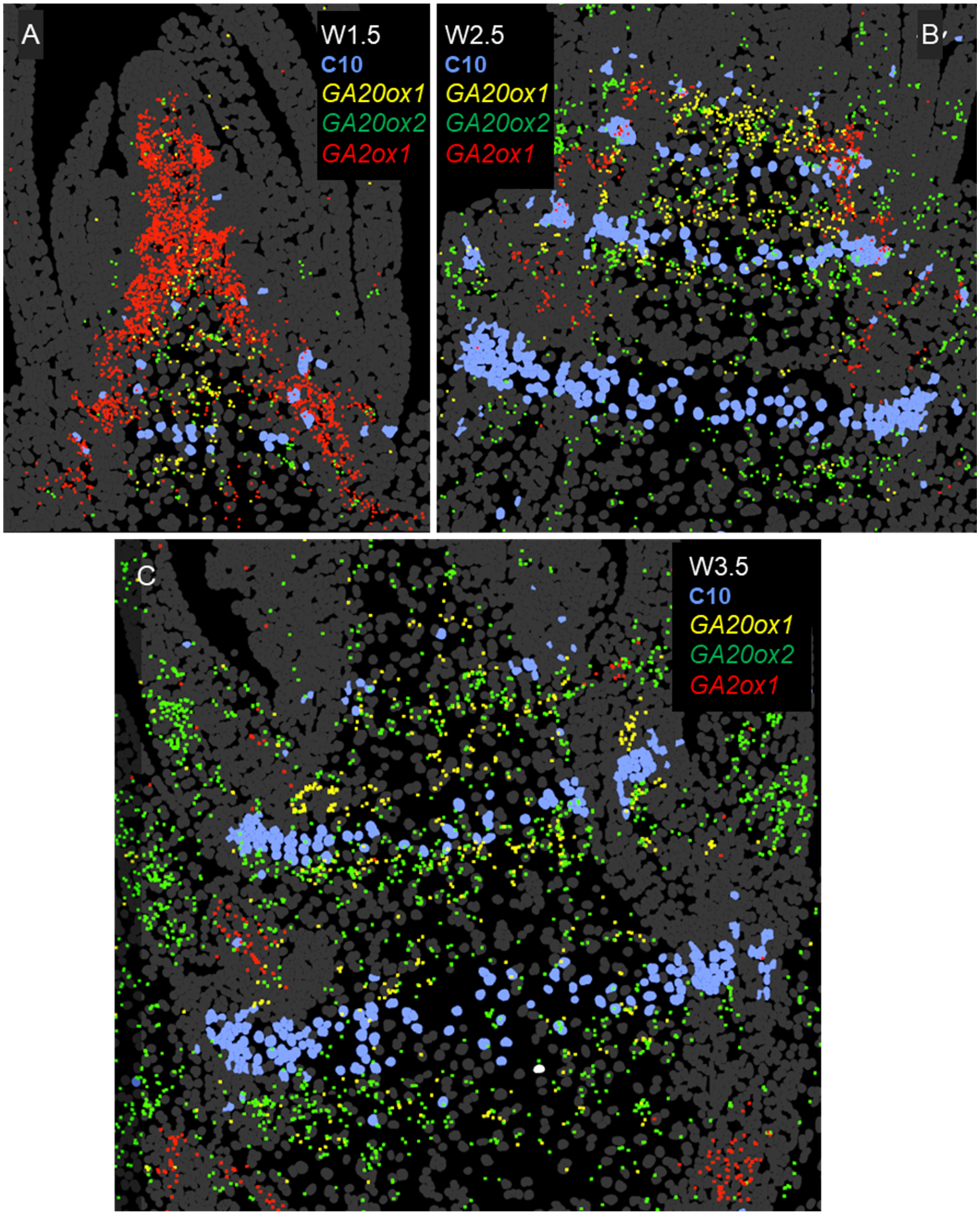

**Fig. S11. Differentially expressed genes in the transition zone between leaves and spike (cluster c19). A-B** Cell included in cluster c9. Differentially expressed genes in c19 include **C-D** *SPL2, SPL14* and *SPL17* and **E-F** TB1 and TB2.

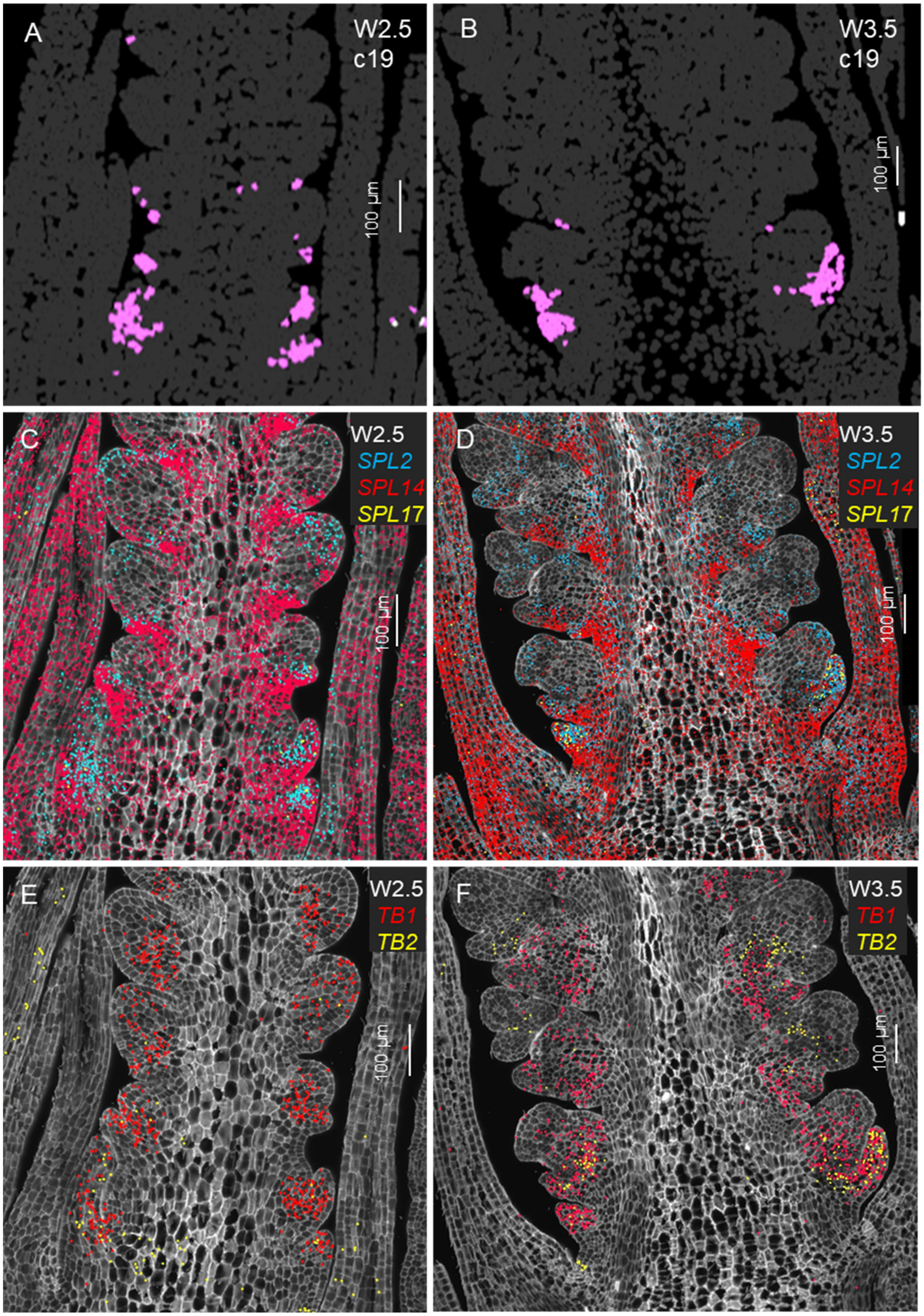

**Fig. S12. Spike central region with preferentially expressed genes.** **A, D** W1.5 initial transition from vegetative to reproductive stages. **B, E** W2.5 late double ridge stage. **C, F** W3.5 floret primordia stage. Cells are stained with calcofluor-white in D-F. **A-C** *AP3* in yellow and *bZIPC4* in pink. **D-F** *IDD4* in yellow and *OSH1* in red. *AP3* genes are also expressed in stamen primordia in the central spikelets at W3.5.

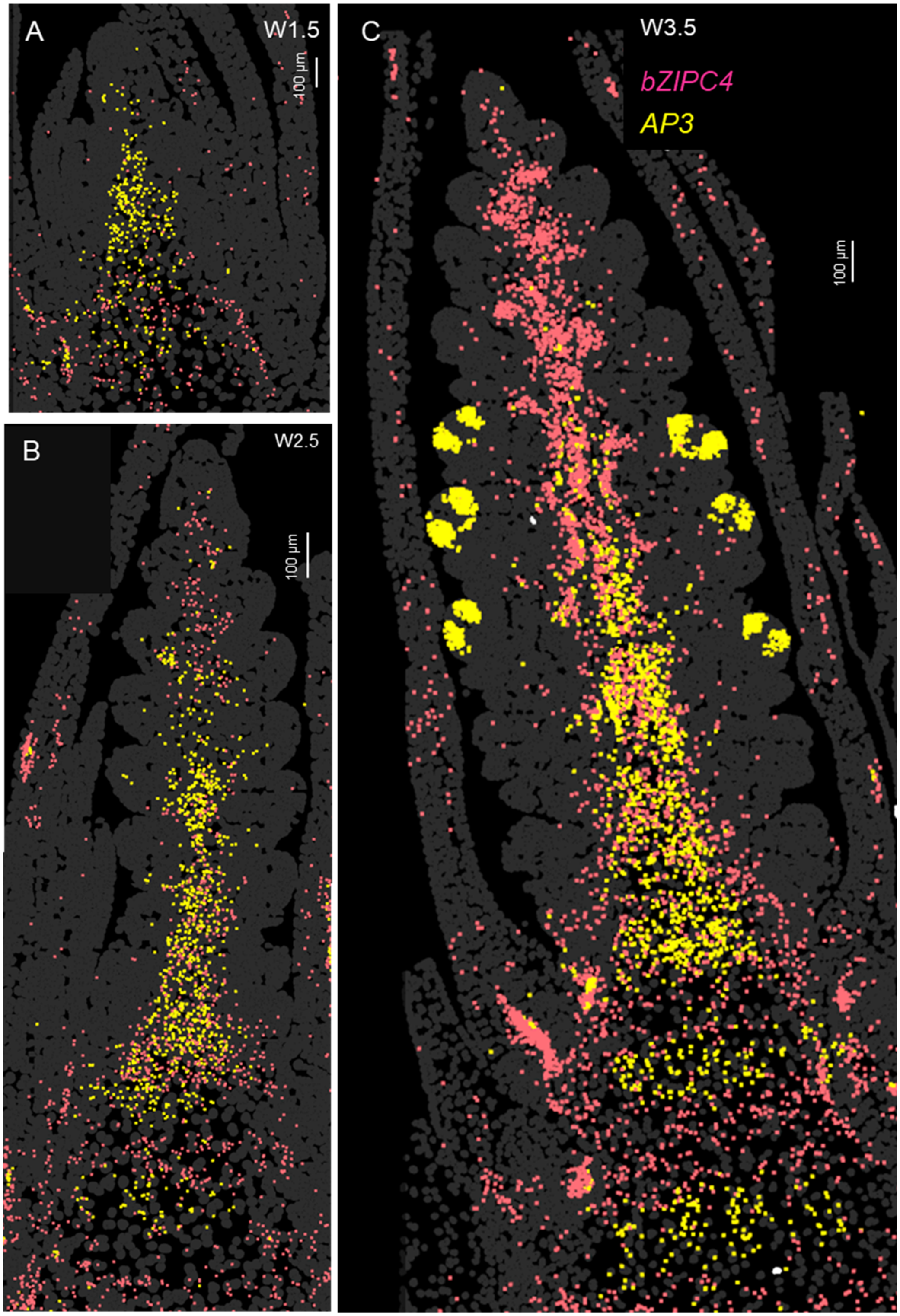

**Fig. S12. Spike central region differential genes. Continuation.**

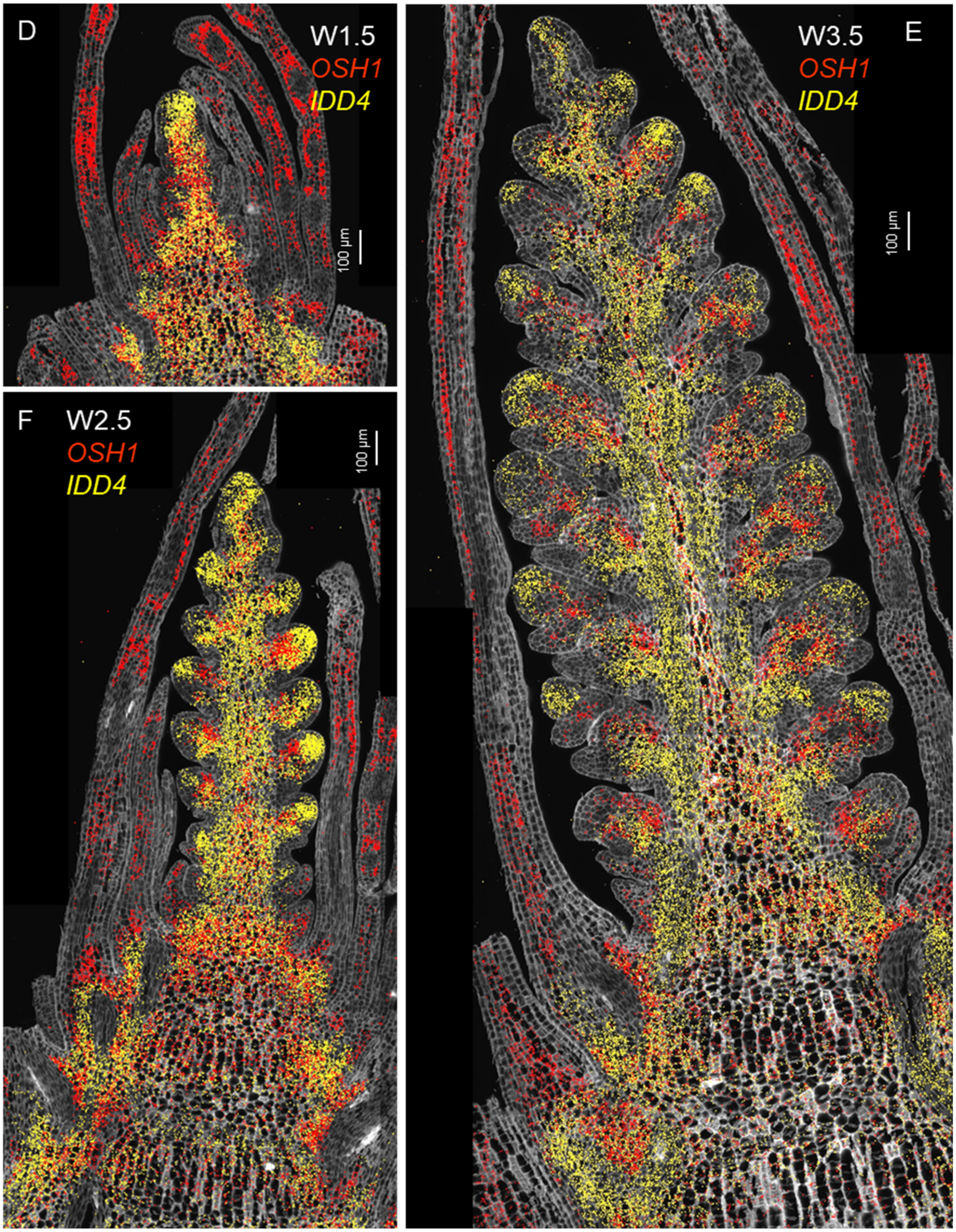

**Fig. S13. Genes preferentially in vasculature cell clusters sc4 and sc13 at W3.5. A** Vasculature clusters c4 (green) and c13 (violet). Same cells with overlapping gene expression: **B** *APL* is a phloem marker located mainly outside c4 and c13, whereas *CEN5* overlaps with c13, **C** *LAX4*, **D** *PIN1a*.

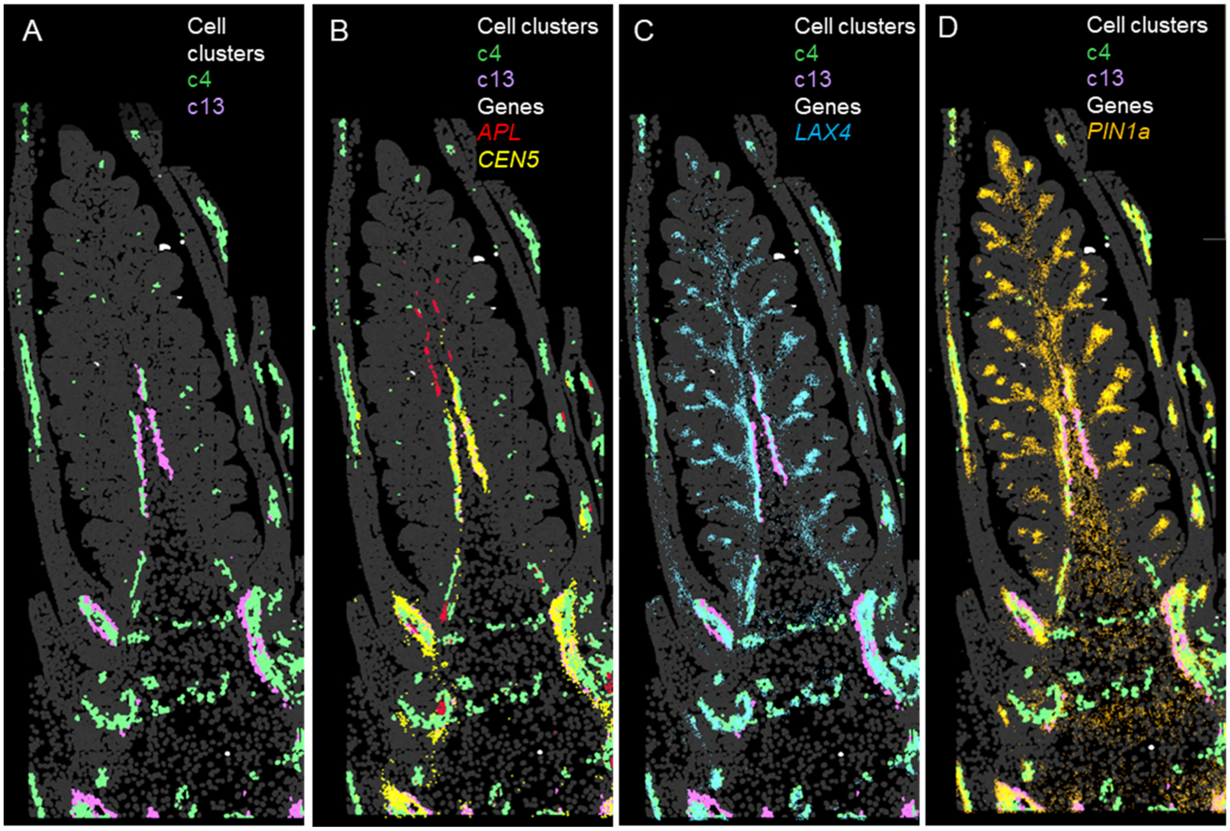

**Fig. S14. Genes preferentially expressed in the suppressed bract region (cluster c6). A, C** W2.5 late double ridge stage. **B**, **D-F** W3.5 floret primordia stage. **A-B** *SPL14*. **C-D** *SPL13*. **E-F** Heatmaps: base of the spike at W3.5. **E** *FUL2*. **F** *VRN1*. **G** Combined *vrn1 ful2* mutant showing bract outgrowth (lost repression of lower ridge), indeterminate IM and vegetative lateral meristems).

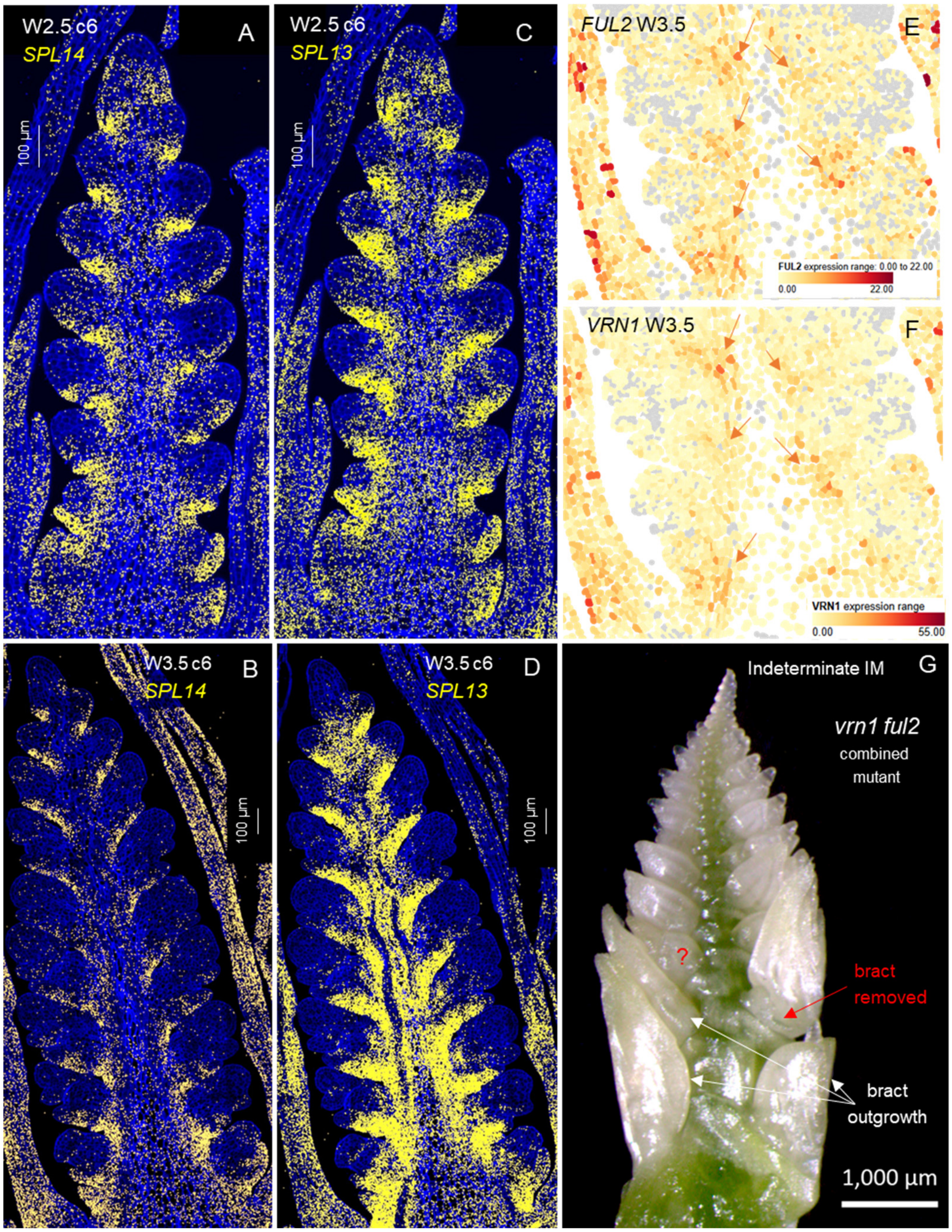

**Fig. S15. Expression profiles of diagnostic genes for cell clusters at the base of the developing spikelet.** **A** W2.5 late double ridge stage. **B** W3.5 floret primordia stage. *SPL14* is preferentially expressed in the suppressed bract region (c6). *PIN1a* marks the developing spikelet vasculature (c9). *FZP* is expressed at the glume axilla (c14). *TCP24* is preferentially expressed at the adaxial boundary of the SM basal region (c16). *SEP1-4* is preferentially expressed in glumes and lemmas (stronger in glumes) and *SEP1-2* in lemmas at W3.5 (c7). Cell walls are stained with calcofluor-white.

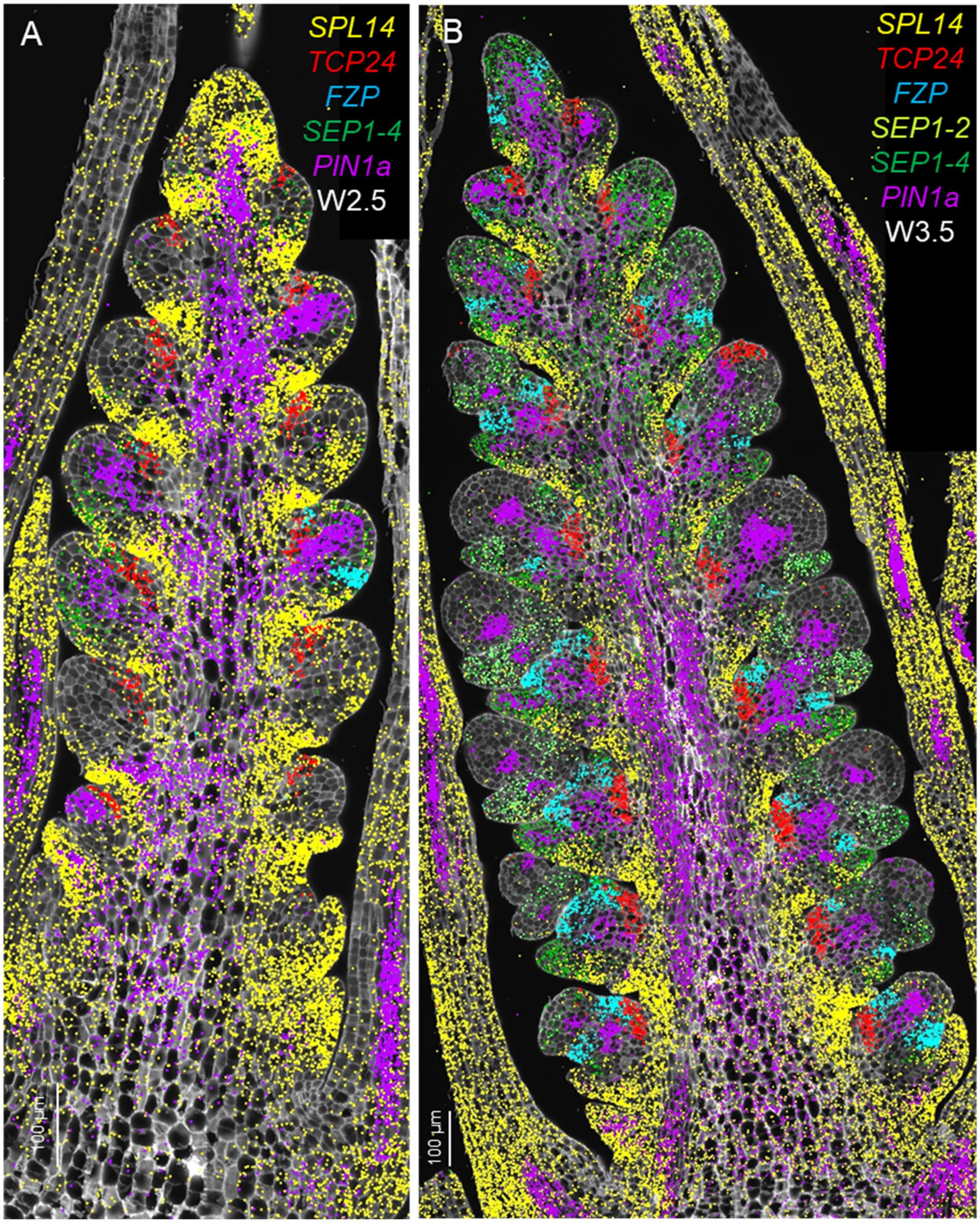

**Fig. S16. Expression profiles of genes preferentially expressed in c14 at W3.5 (in addition to *FZP*).** **A-B** Cells from cluster c14 (*FZP*) are marked in blue and those from c16 in violet (*TCP24*). **A** *WOX7*. **B** *BIF2*. These two genes overlap well with the cells of c14 but are also present in other cells outside c14. **C** Detail of W3.5 stained with calcofluor-white: *FZP* in blue, *BIF2* in yellow and *WOX7* in red.

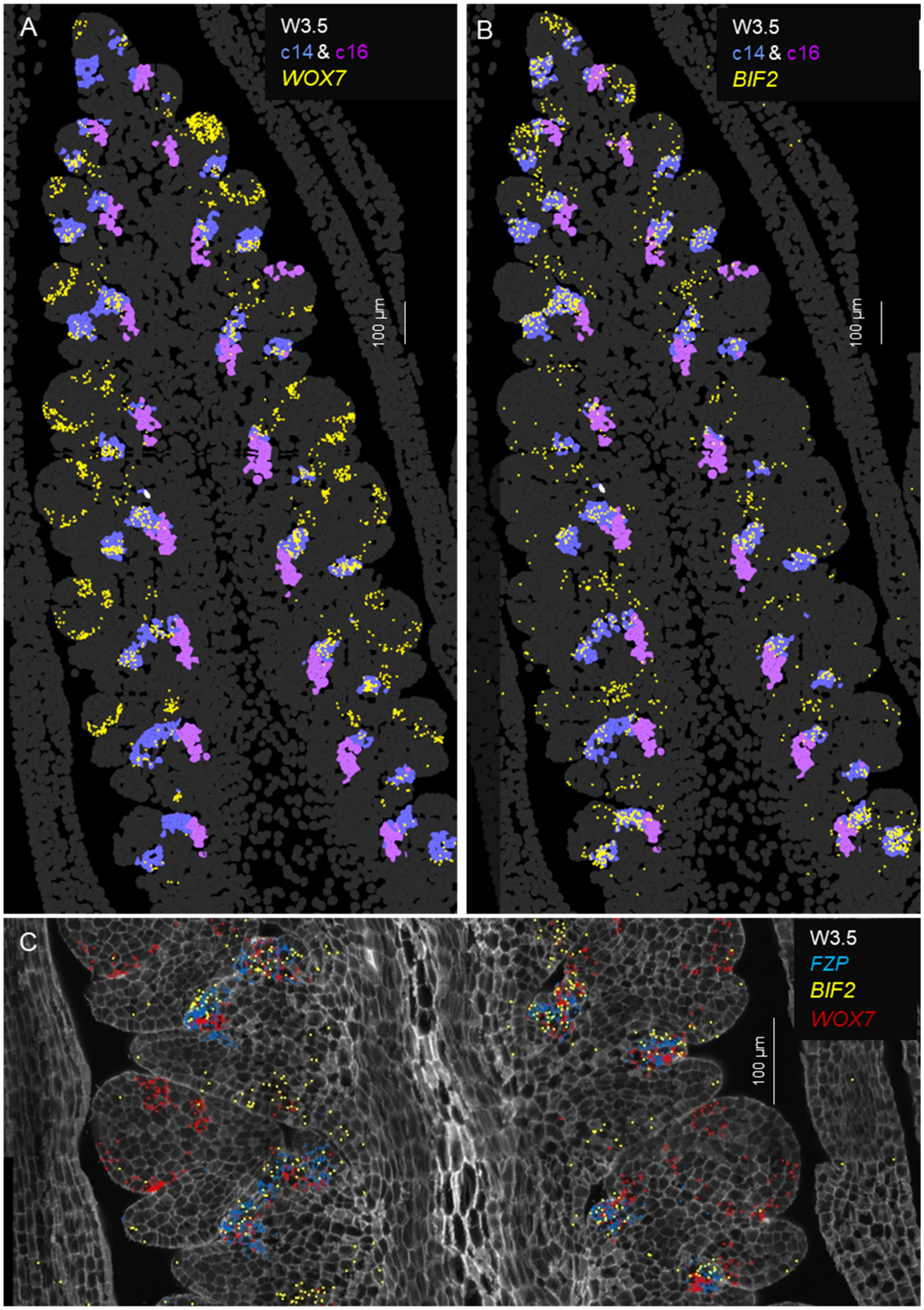

**Fig. S17. Functional validation of *FZP*.** Effect of combined loss-of-function mutations in *fzp-A* and *fzp-B* (*fzp*) on spike development. **A** Schematic representation of *FZP* gene structure and location of the CRISPR induced deletions. **B-C** Spike of **B** wildtype and **C** *fzp*. **D-E** Detail of a spike node with a spikelet in **D** wildtype and **E** *fzp*. **F** Dissection of recurrent glumes at *fzp* node indicated in panel D by a red square. **G** Abnormal floral organs observed among the recurrent glumes. **H** Rachilla at the same *fzp* node with abnormal flowers. **I** Scheme of the developmental steps leading to the recurrent glumes: In each glume a new axis emerges from the axillary meristem and produces additional glumes that repeats the same process. At each cycle, fewer and smaller glumes are formed, until the SM is exhausted. The axes at different cycles are indicated by different colors (black → blue → green). Abnormal flowers are indicated by red circles. **J** Dissection of a wildtype Kronos spikelet (only the first floret is shown.). **K** Scheme of a wildtype Kronos spikelet.

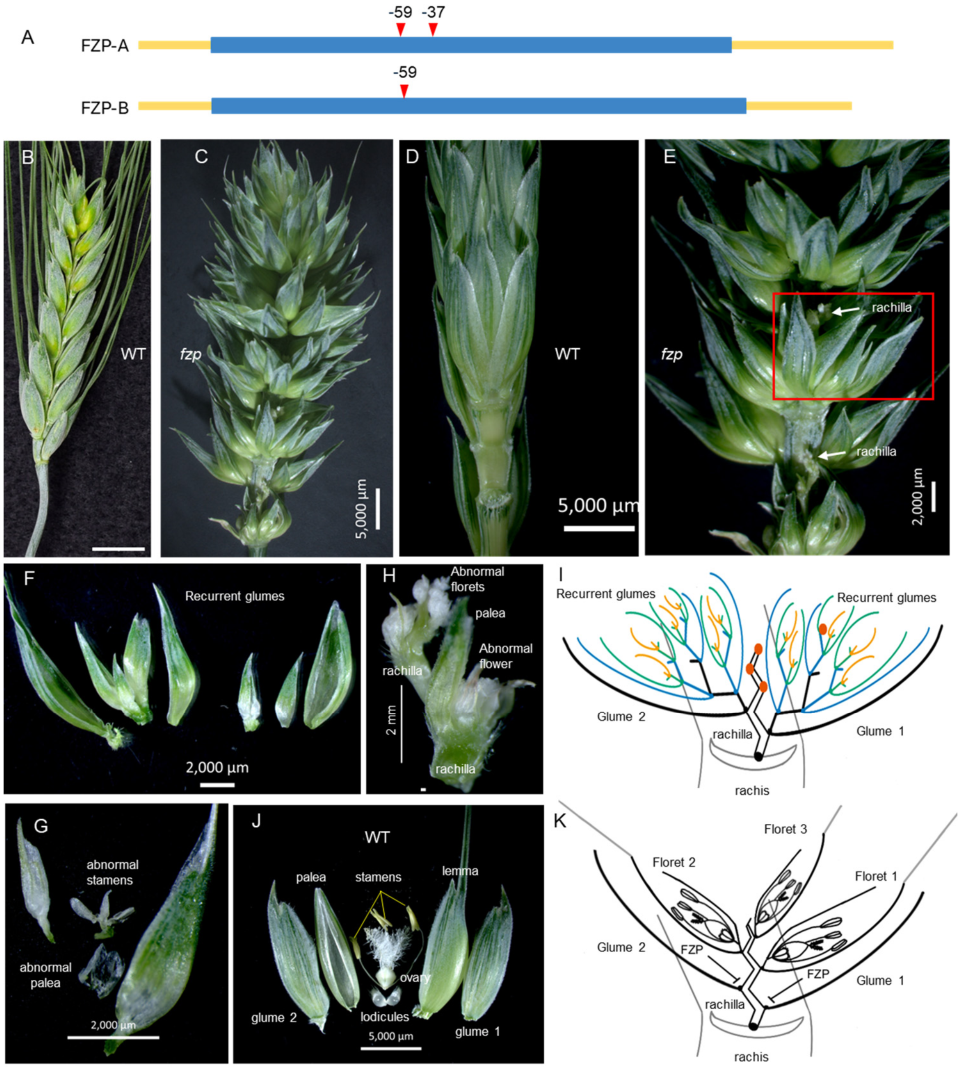

**Fig. S18. Expression profiles of genes preferentially expressed in the spikelet meristem (SM). A** *HAF1* and *BIF2*. **B** *HAF1* and *AGL6* **C** *ULT1* and *WOX7*. **D** *ULT*, *WOX7* and *BIF2*. **A-C** W2.5 late double ridge stage. **B-D** W3.5 floret primordia stage. c8 = early SM development, c18 = floret meristem FM. **E-G** Floral homeotic genes expressed the FM (c18). **H** *WAPO1* is expressed at the FM in c18, whereas *LFY* is highly expressed in a band between the lemma primordia and the FM (c17).

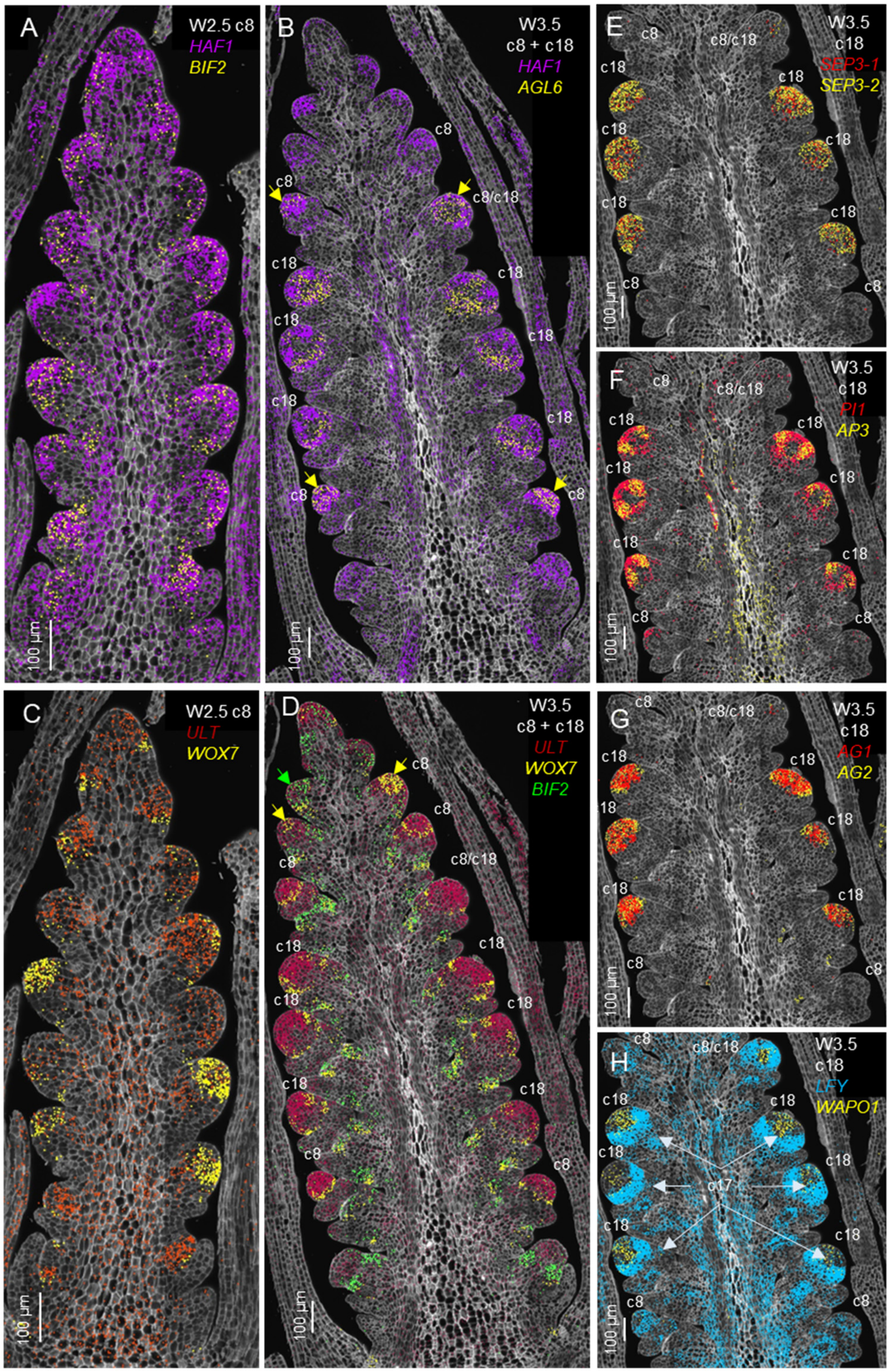

**Fig. S19. Selected regions in the inflorescence meristem.** Example for the changes in hybridization density in the terminal and first two lateral meristems between W2.5 (IM+2LM) and W3.5 (SM + 2 glume primordia). **A** *SPL14* at W2.5. **B** *SPL14* at W3.5. **C** *FZP* expression at W3.5 is included as a control to show that the IM has already initiated the transition to a terminal spikelet.

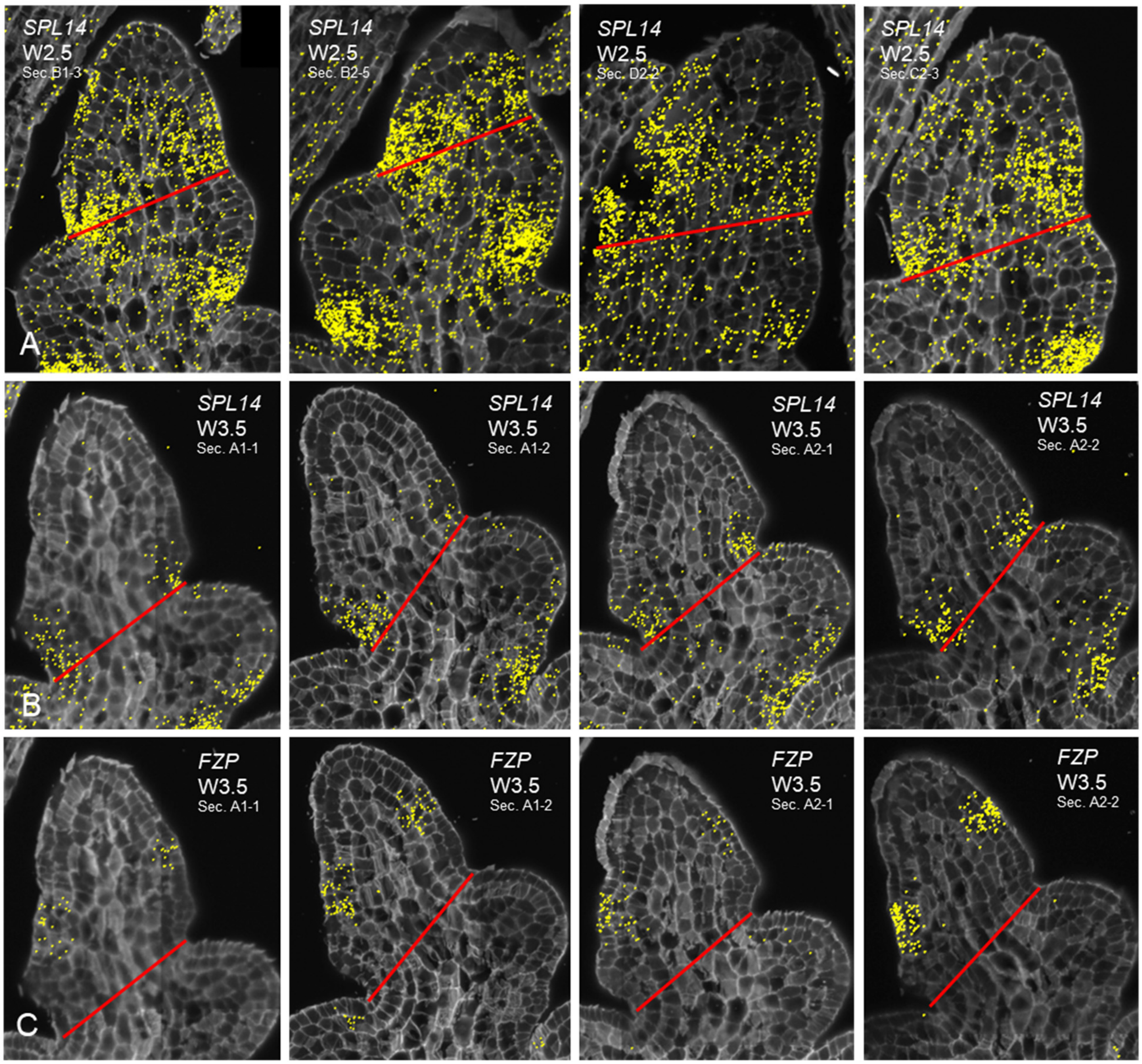

**Fig. S20. Genes with significant differences in hybridization density in the terminal region between W2.5 and W3.5.** The selected region included the terminal meristem and the two youngest lateral meristems. At W2.5 the TM= IM and LM= SM, whereas at W3.5 TM= SM and LM= glume meristems. Genes downregulated at W3.5: **A** *SPL14*, and **B** *HAF1*. Genes up-regulated in W3.5: **C** *BC1* **D** *FUL2*, **E** *SOC1-2*, **F** *SEP1-4*, **G** *bZIPC1*, **H** *bZIPC3*, **I** *FDL2*, **J** *FDL6* **K** *PIN1b*, and **L** *CRN*. Four different sections at W2.5 and two at W3.5 (each with two subsamples) were used in the analyses. * $P < 0.05$, ** $P < 0.01$, and *** $P < 0.001$ based on two-tailed *t*-Tests. *HAF1* was marginally not significant $P < 0.12$) but was included since there were not many downregulated genes. Raw data and statistical analyses are presented in Data S5.

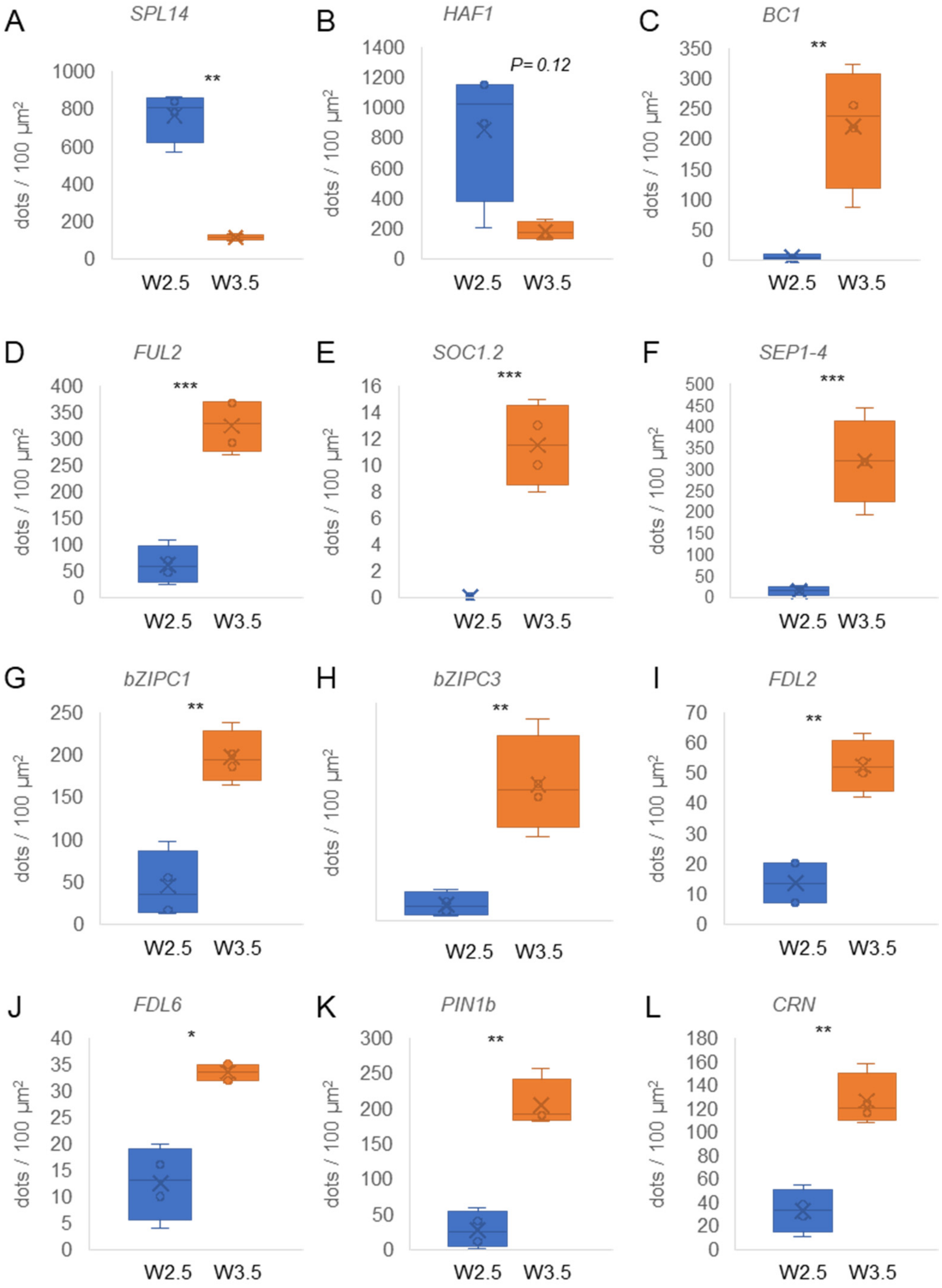

**Fig. S21. Functional validation of *SPL14.*** **A** *SPL-A14* and *SPL-B14* gene structure and location of the CRISPR induced mutations **B** Conservation across different monocot species of the region including the frame shift insertion and premature stop in *spl-A14* and the C121 deletion in *spl-B14*. **C-H** Effect of the combined *spl-A14 spl-B14* mutations on **C & E** spikelet number per spike, **D &F** plant height, **G** heading time, and **H** leaf number. Plants are $T_2$ transgenic lines homozygous for mutations in both homeologs and without the CRISPR-CAS9 vector. Bars are the averages of 12 different $T_1$ homozygous edited plants and error bars are s.e.m. ***= $P<0.0001$. Raw data is available in Data S6.

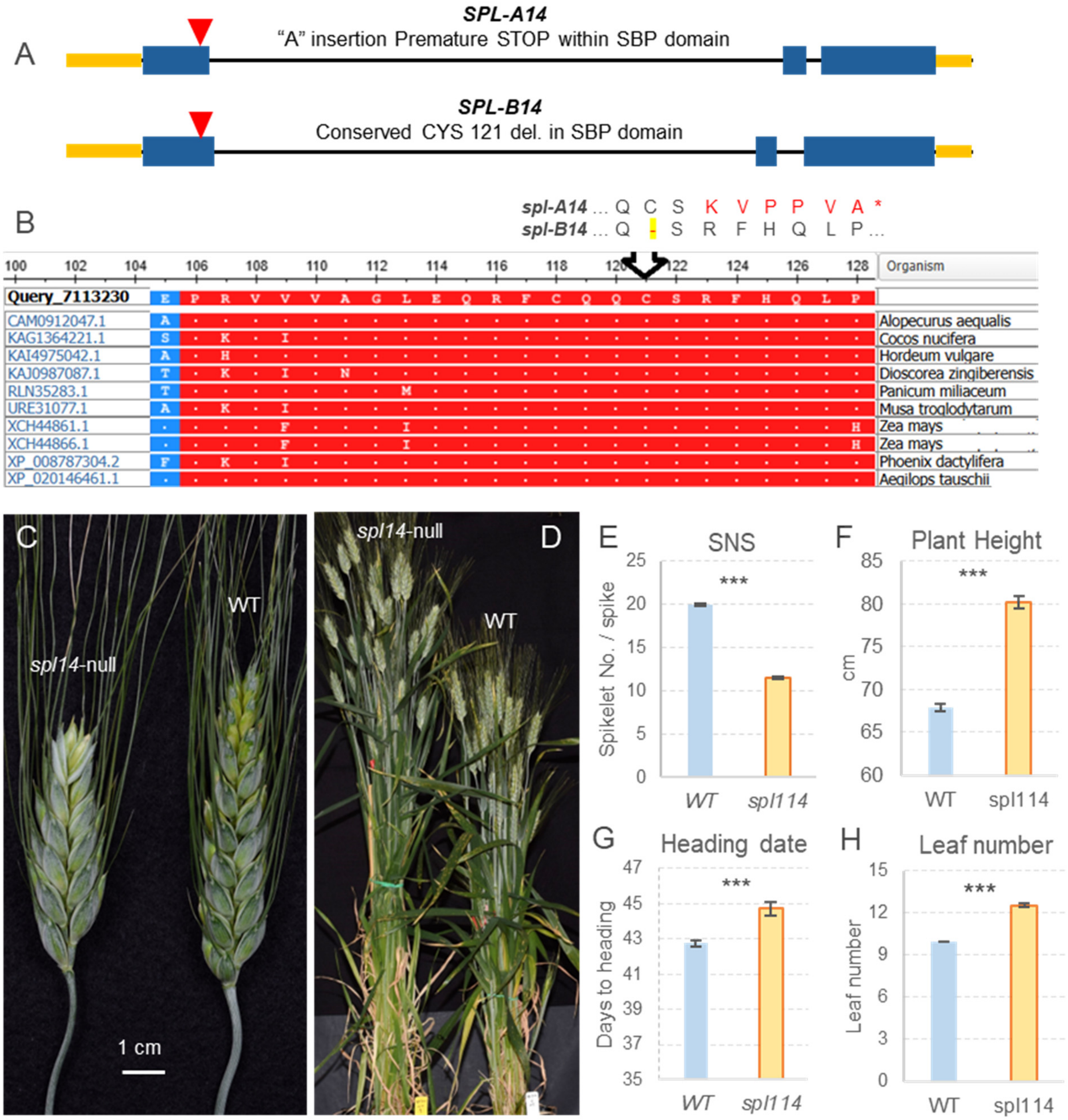

**Fig. S22. Trajectory analysis.** The 21 cell clusters from the smFISH analysis were leveraged to constrain trajectory inferences generated with program Monacle 3. The cluster numbers and colors are the same as in Fig. 1. Cluster c8, which includes both inflorescence meristem (IM) and spikelet meristems (SM) was used as root. Starting from c8 one trajectory went towards the floral meristem (c17 and c18), and another one in the opposite direction towards the transition zone (c19) and the repressed bract (c6). A third trajectory trifurcated into the glume and lemma primordia (c7), the central cluster associated with the spikelet vasculature, and the boundary clusters including a trajectory that bifurcates into the glume axilla (c14) and the adaxial boundary (c16). The ground tissue at the center of the spike (c3) is connected by trajectories to both the meristem (c8) the suppressed bract (c6) and the leaves. Cluster c11, associated with dividing cells, was placed close to the spike meristems (c8) and the leaf buds (c20). Leaf clusters were not analyzed in this study.

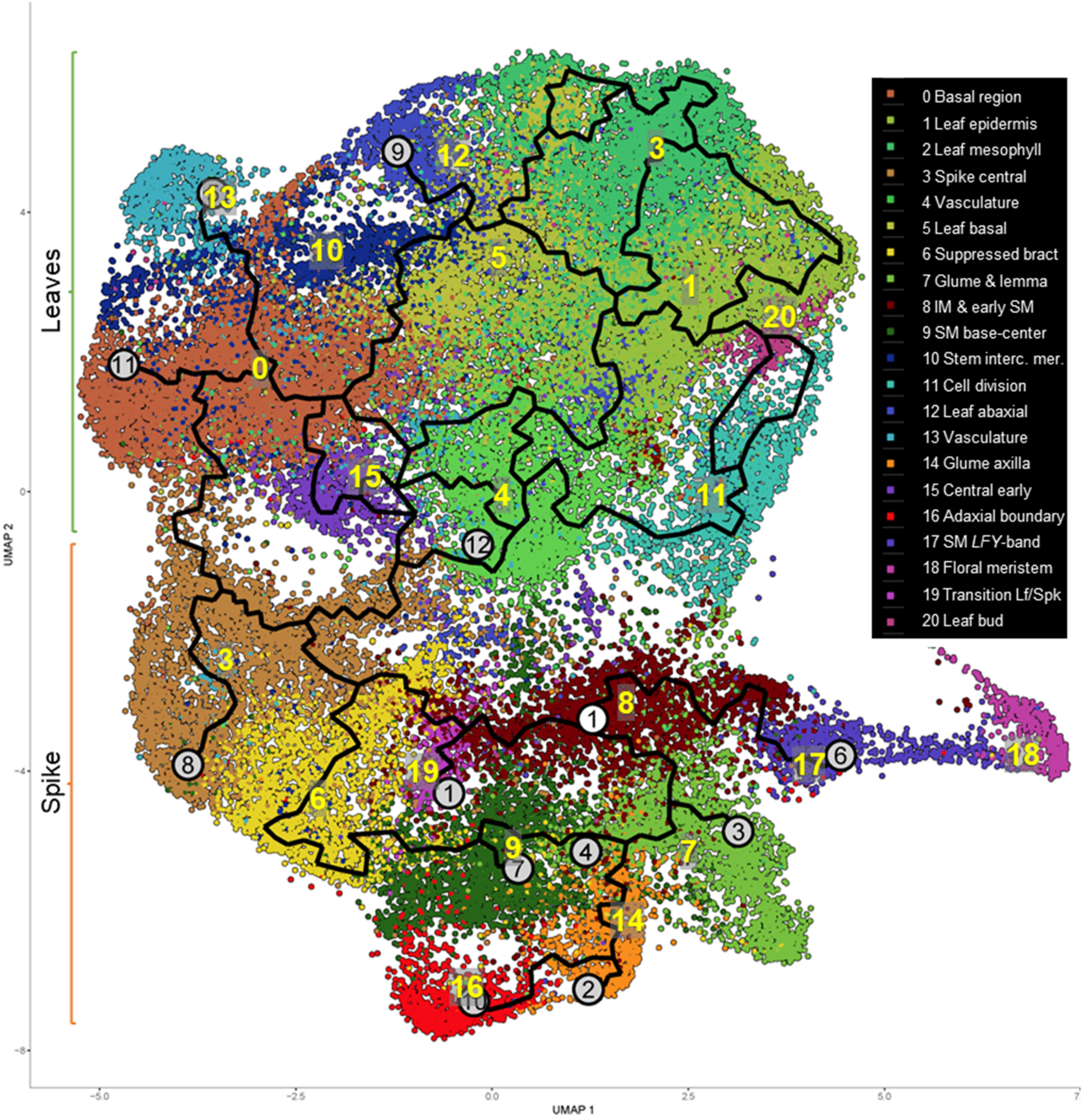

**Fig. S23. Comparison between initial and final clustering. A** Initial 25 clusters including 9 cell-cycle clusters showing good integration of the three replications and the two stages. **B** Identification of cells in the S (*HISA1*) and G2/M (*CYCB2*) phases in the UMAP. **C** Bubble-plots for *CYCB2* and *HIS2A1*. **D** Final 23 clusters used in this study showing good integration of the three replications and the two stages.

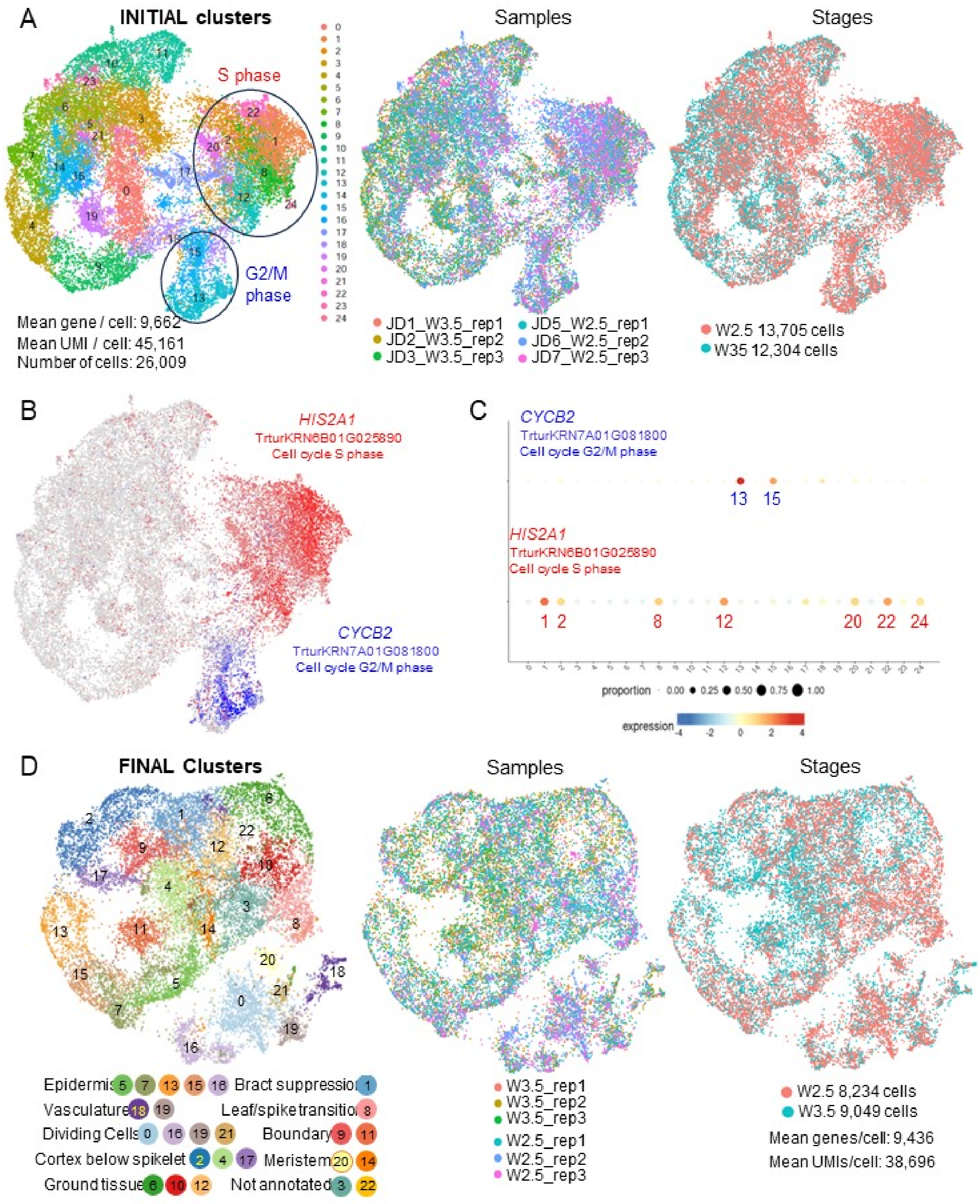

**Fig. S24. RNA-seq for three different sections of the spike.** **A** Representative Kronos spike at W3.0 (glume development stage) and the three regions where the developing spikes were sectioned (dotted red lines). **B** Venn Diagram indicating the number of differentially expressed genes among the three different regions based on Tukey test (Data S11). **C** Principal component analyses for the nine samples. Red color indicates the basal region, green the mid region and blue the tip. Different geometric figures indicate the three replications.

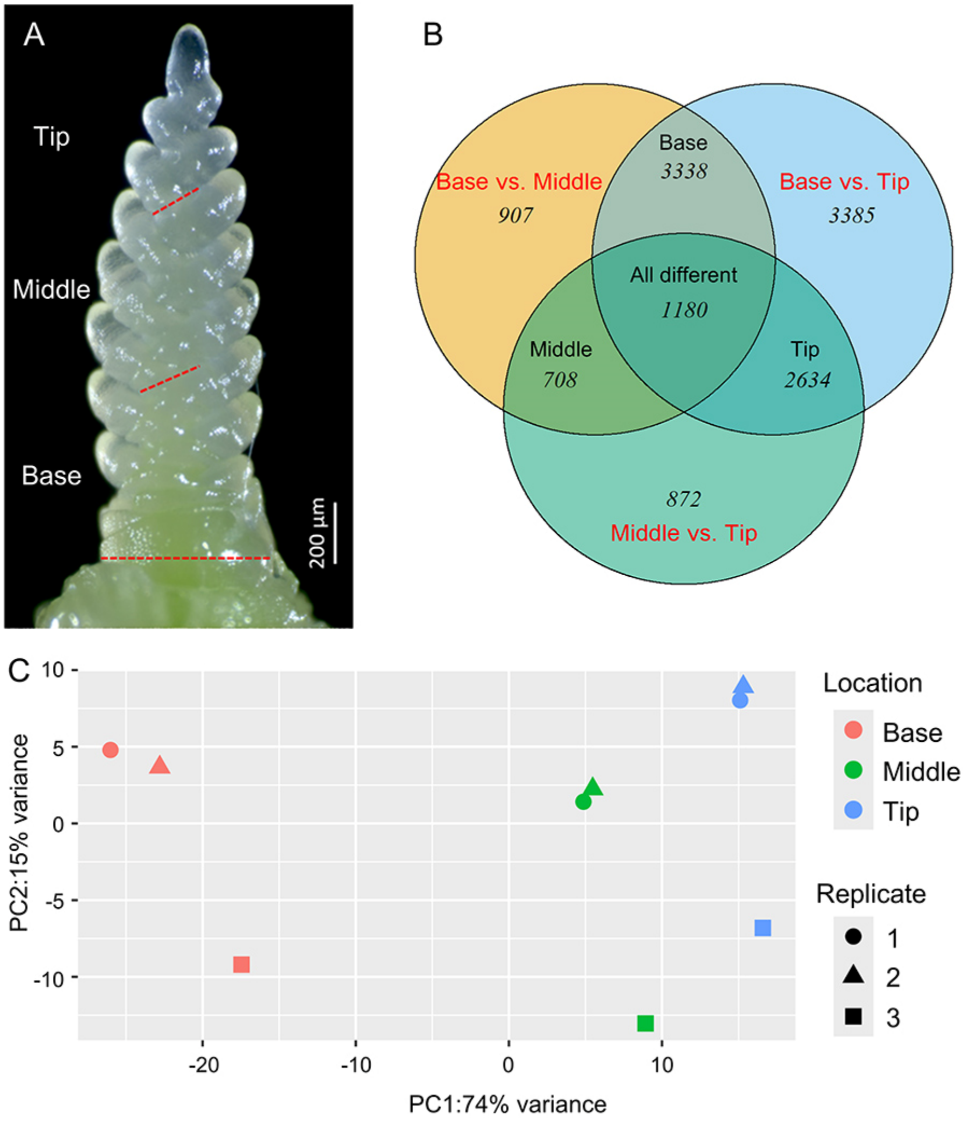

**Fig. S25. Cell cycle clusters. A-F** UMAPS and **G** bubble plots of known cell-cycle marker genes for clusters sc0, sc16, sc19 and sc21. Annotation of the marker genes and references are available in Data S1. Additional genes preferentially expressed in cell-cycle clusters are available in Data S9.

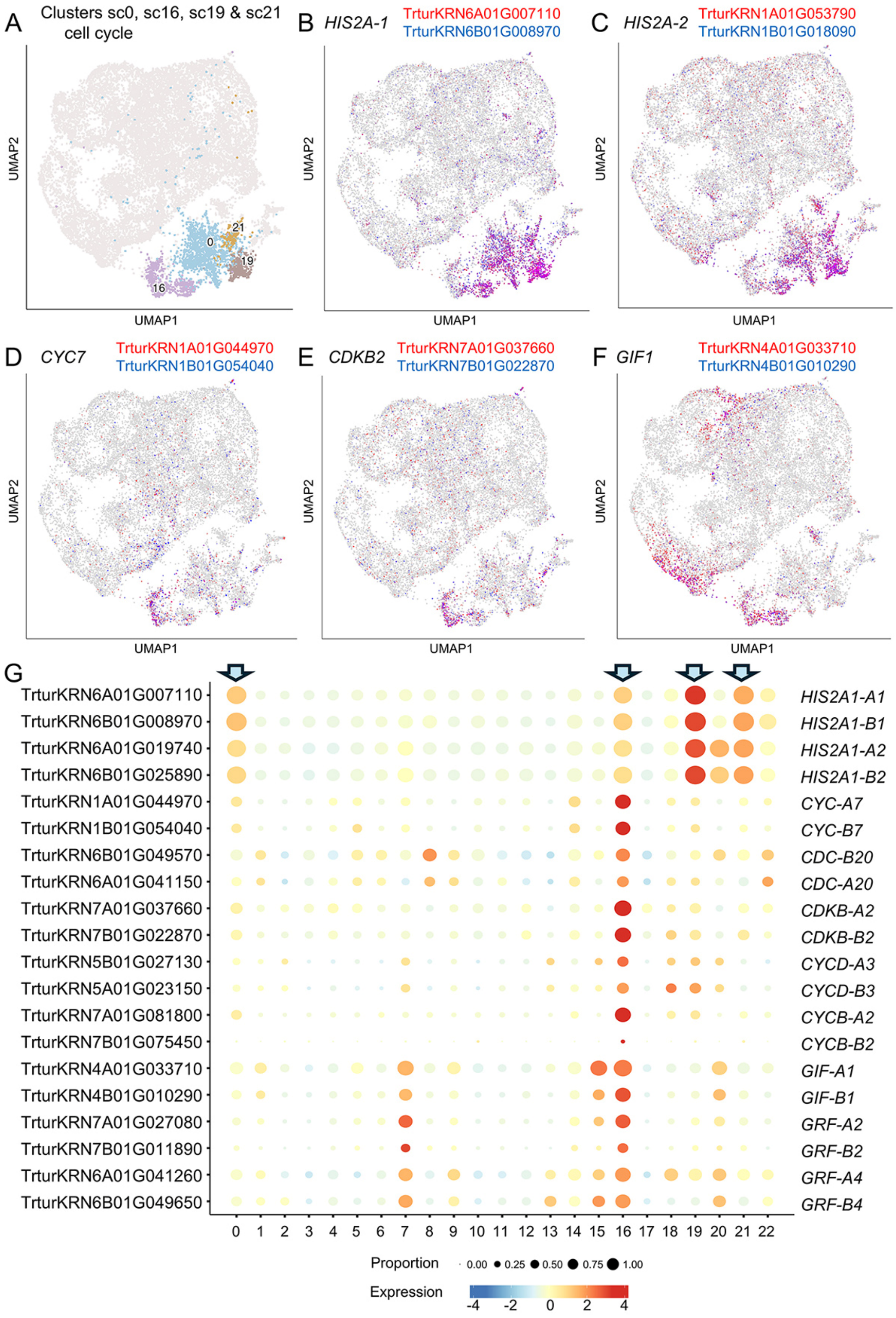

**Fig. S26. Common markers for single-cell epidermal clusters. A-F** UMAPS and **G** bubble plots of known epidermal marker genes for clusters sc5, sc7, sc13, sc15 and sc16. Epidermal clusters showed relatively low expression of the homeodomain-box gene *OSH1*. **H** Spatial pattern of epidermal genes using imputed expression. A-genome homeologs: fill color using yellow-red scale, B-genome: border color using green-blue scale. Annotation and references for marker genes are available in Data S1. Additional genes preferentially expressed in epidermal cell clusters are provided in Data S9.

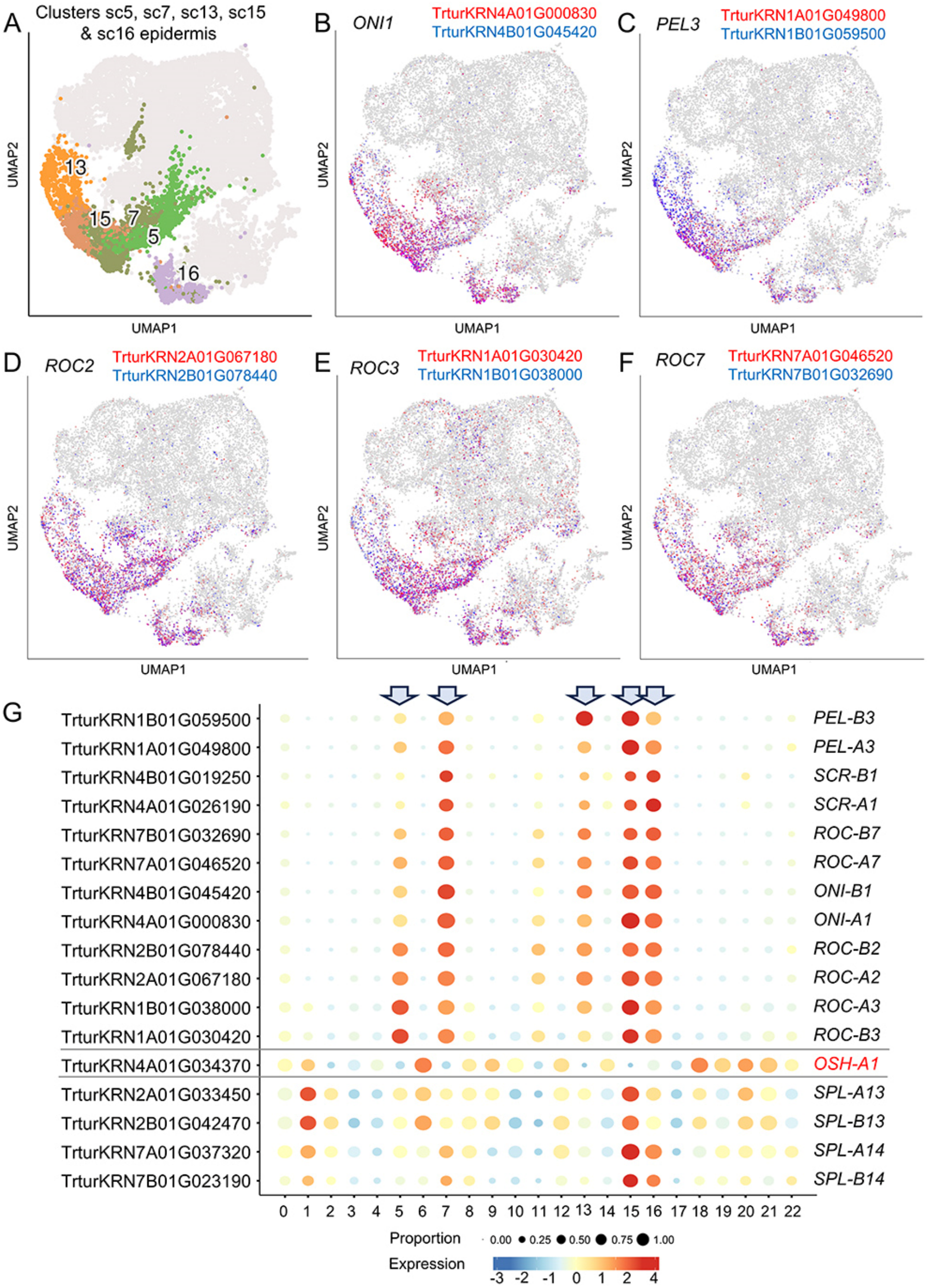

**Fig. S26. Continuation.**

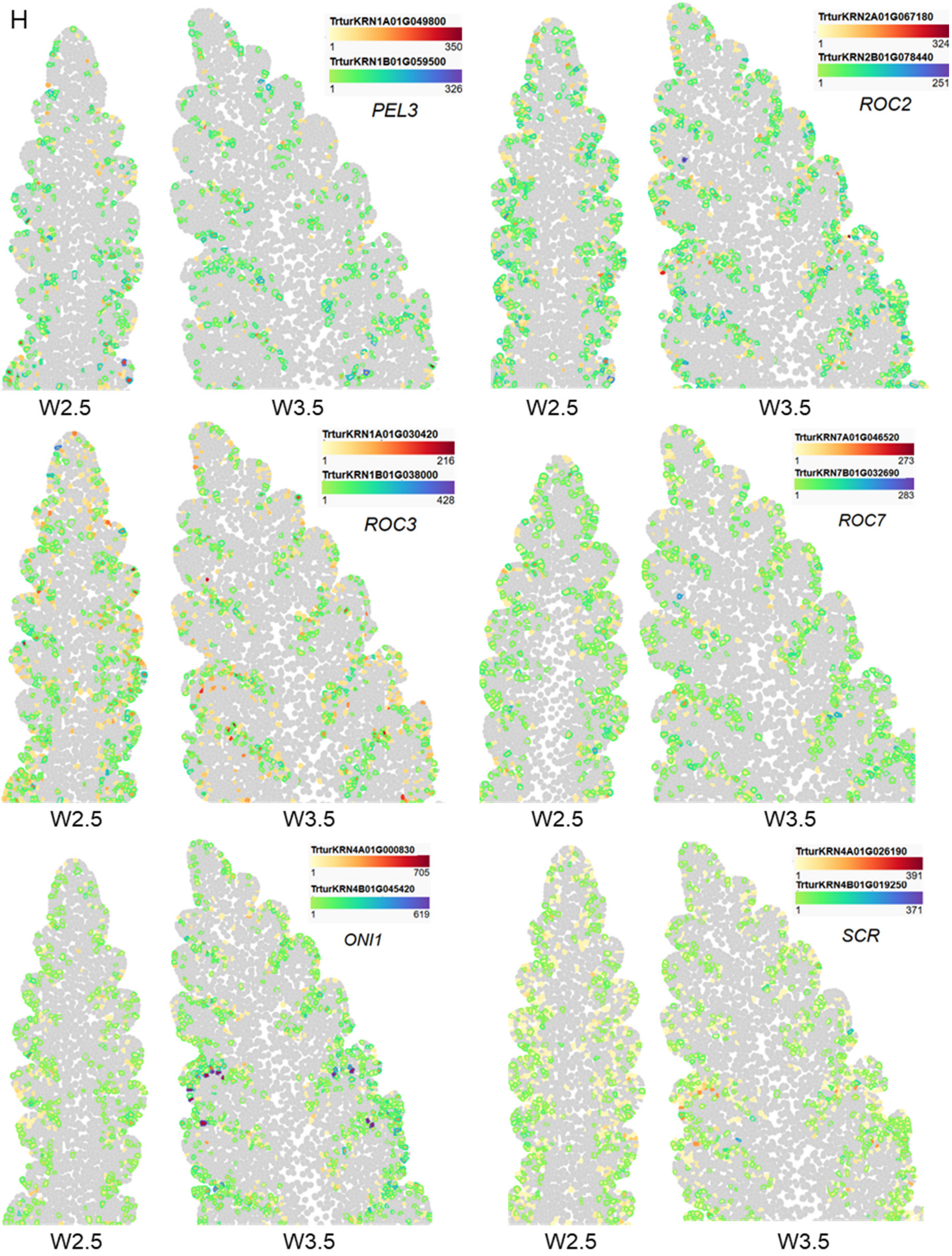

**Fig. S27. Differential genes among single-cell epidermal clusters. A-I** UMAPS of markers that differentiate the epidermal clusters. Annotation and references for marker genes are available in Data S1. UMAPS for epidermal – cell-cycle cluster sc16 are provided in Fig. S25. **A-C** Markers for sc7. **A** *AGL6*, **B** *WOX3b* and **C** *YAB3*. **D-E** Markers for sc15. **D** *SPL13* and **E** *SPL14*. **F-G** Markers for sc13. **F** *CUC3* and **G** *LAX1*. **H-I** Markers for sc5. **H** GSTF13 and **I** CHT6. **J-P** Imputed expression. **J** *WOX3b*. **K** *CLE33* (meristem sc7). **L** Lateral organs *YAB3* and *YAB4*. **M-P** Imputed expression of boundary genes expressed in cluster c14 and c16 at W3.5 including epidermal cells. **M** *TCP22*. **N** *CUC3*. **O** *LAX1*. **P** *DP1*. Only one homeolog is shown using a green-blue scale for the cell-border color. Gene identification numbers and annotations are in Data S1. Additional genes preferentially expressed in the different epidermal cell clusters are in Data S17.

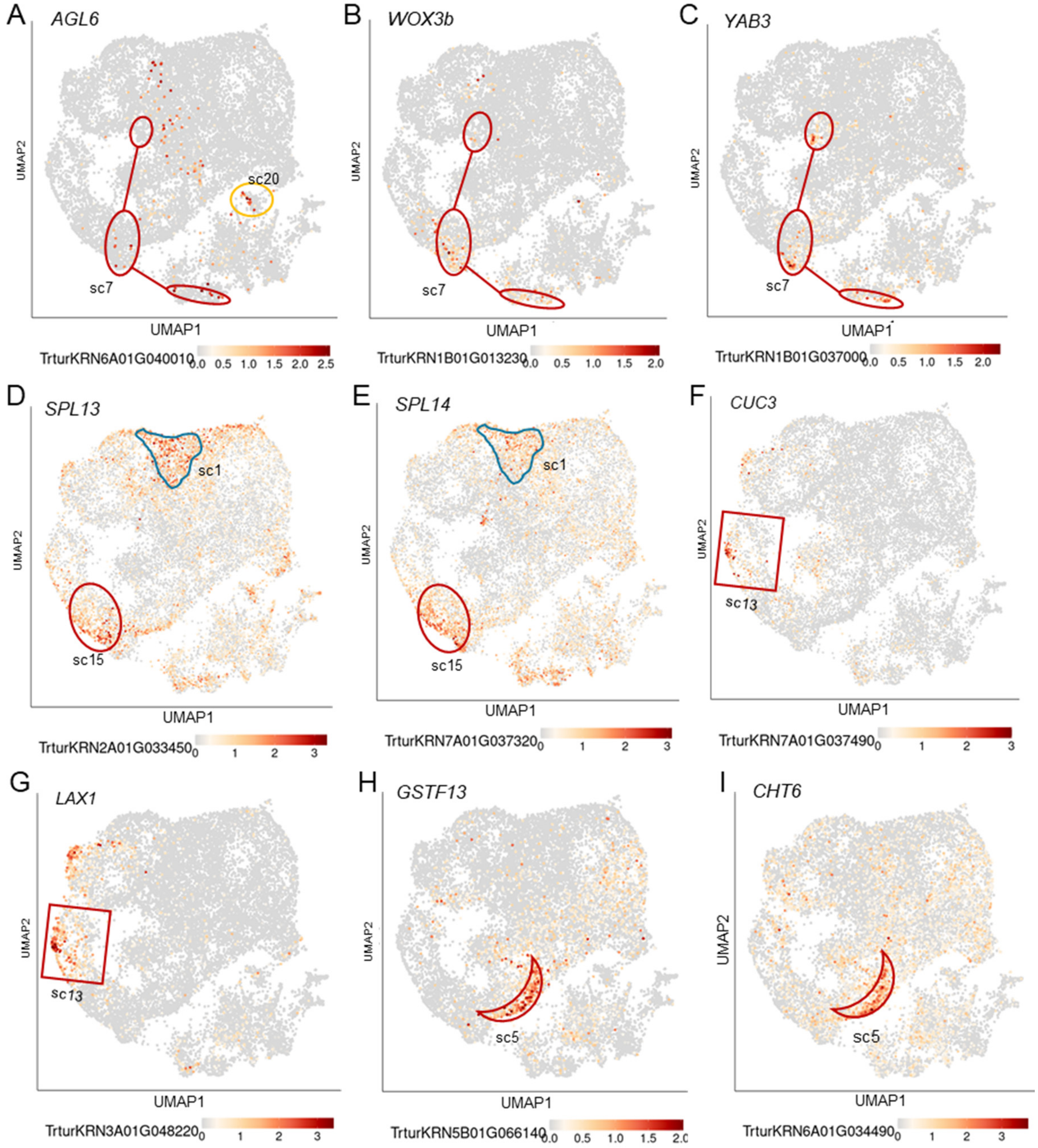

**Fig. S27. Continuation.**

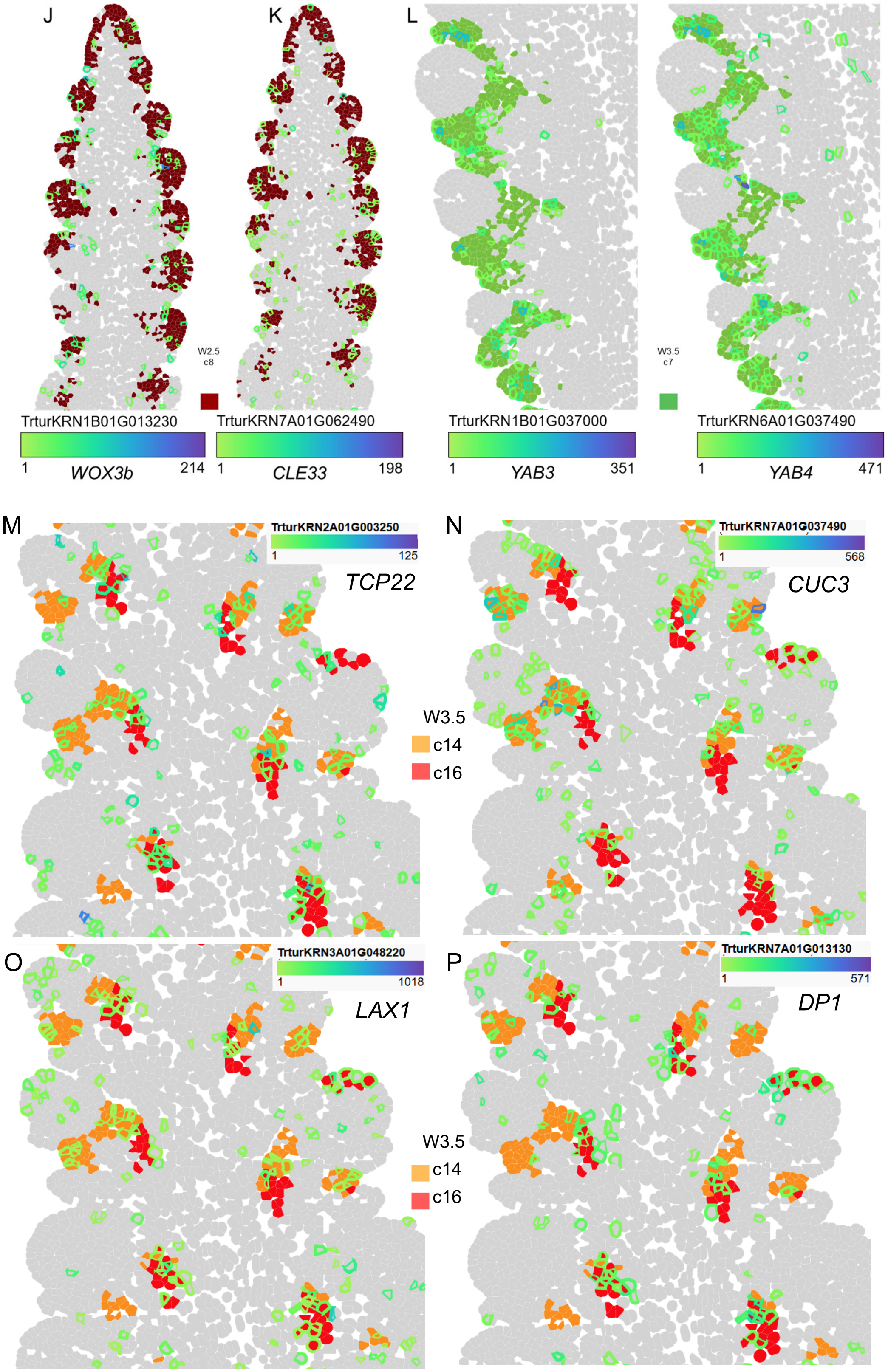

**Fig. S28. Vasculature clusters.** **A** UMAPS showing cells from clusters sc18 and sc19. **B-C** Phloem markers *APL* and *JUL1*. **D** Procambium marker *LAX4*. **E-F** Xylem markers *WAT1* and *XCP1*. **G** Bubble plots of known vascular marker genes from clusters sc18 and sc19. **H-I** Spatial pattern of vascular genes using imputed expression at W3.5. A-genome homeologs: fill color using yellow-red scale, B-genome: border color using green-blue scale. **H** Vascular markers (phloem marker *APL* and procambial marker *LAX4* included in the smFISH study are included for reference). **I** Hormone-associated vascular markers. Annotation of the marker genes and references are available in Data S1. Additional genes preferentially expressed in vascular and pro-vascular clusters sc18 and sc19 are provided in Data S9, and sub-clustering of sc18 in Data S18. Cluster sc19 includes both cell cycle and vasculature markers.

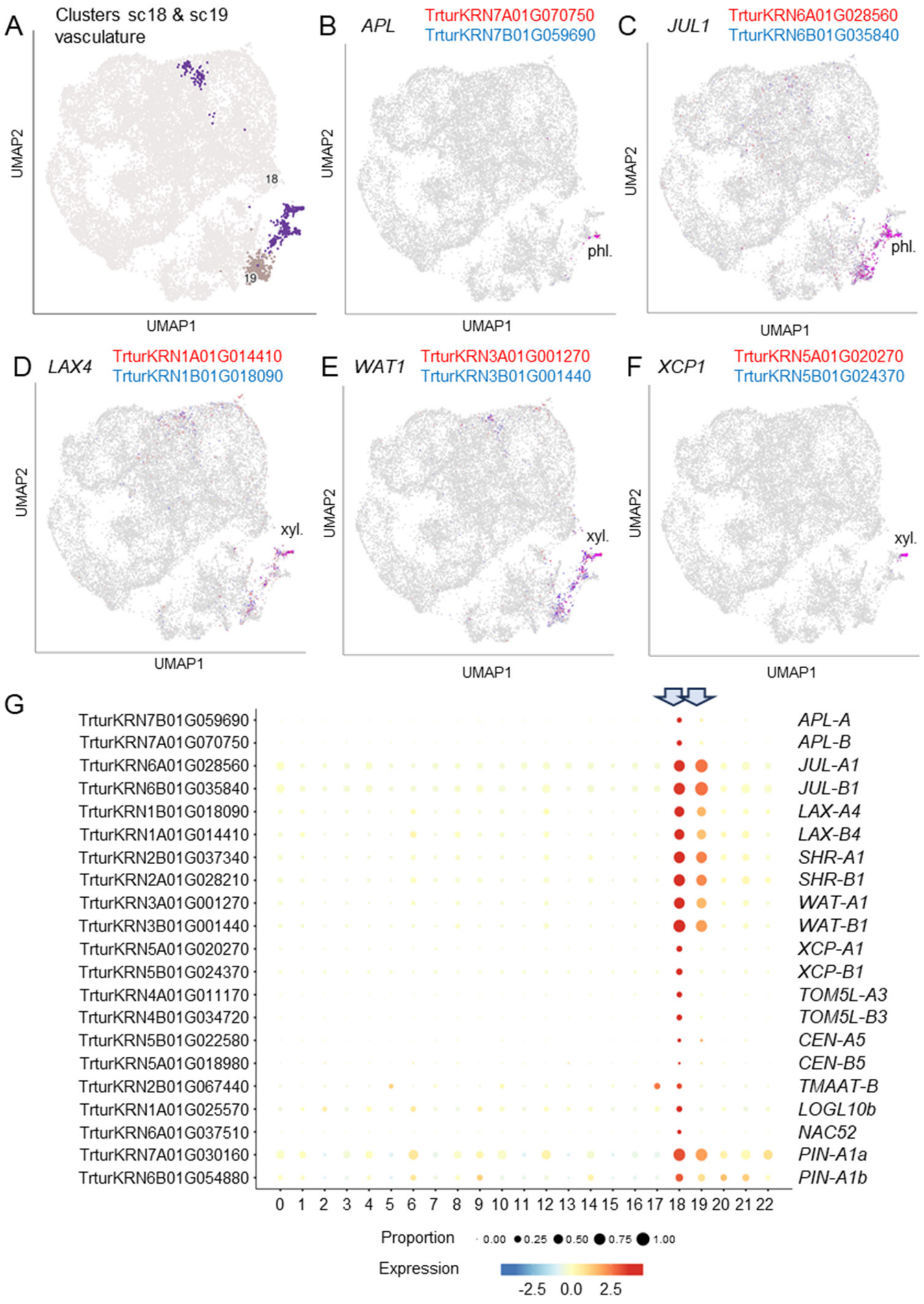

**Fig. S28. Continuation**. Genes expressed in the vasculature at W3.5. Vasculature: *SHR1*, procambial: *LAX4*, phloem: *JUL1*, *DOF19*, and *APL*, early xylem: *WAT1*, xylem: *TMO5L1, TMO5L3*, *TMAAT*, *XCP1*, *LOG10b*, and *NAC52*.

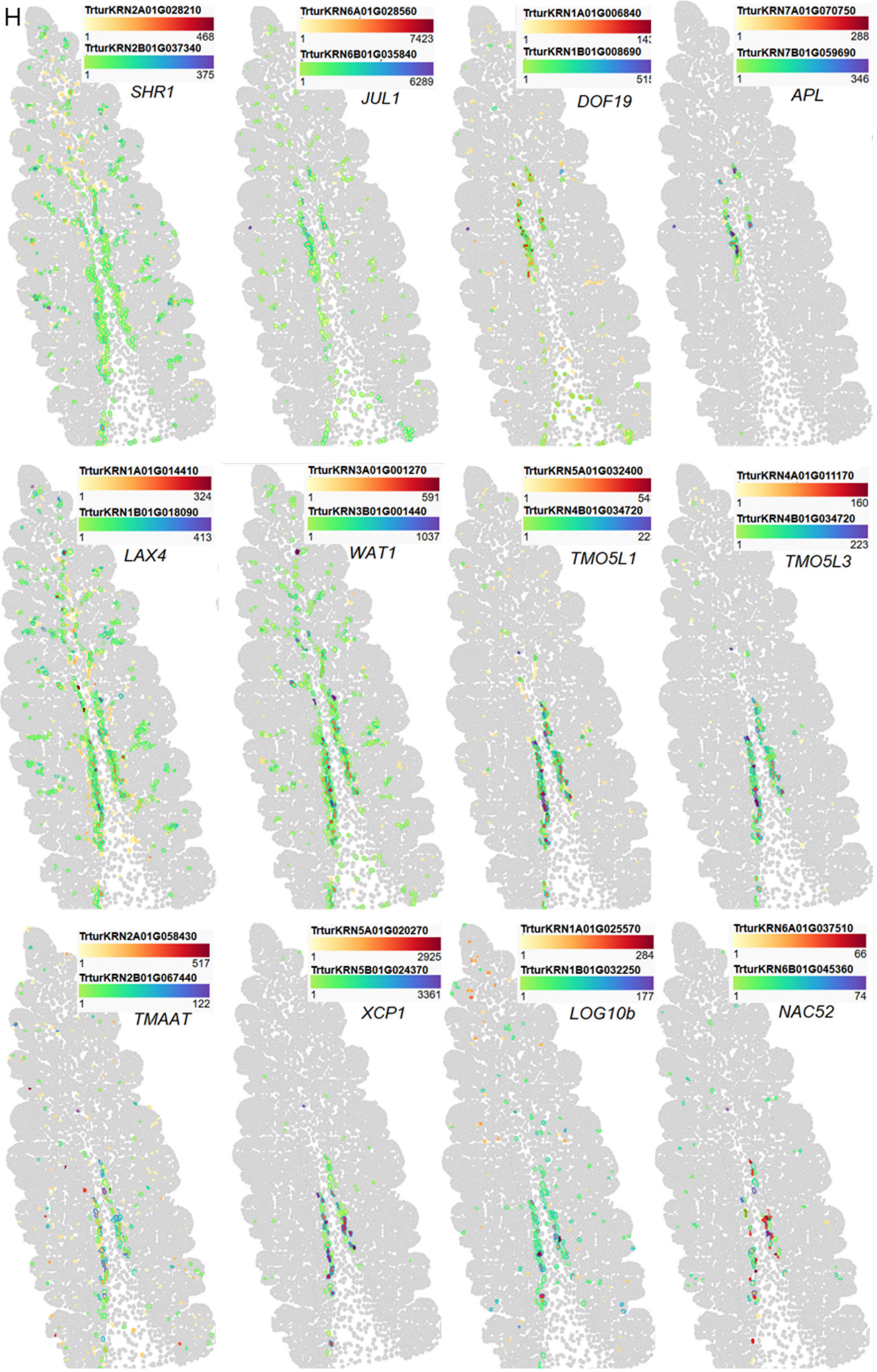

**Fig. S28.** Continuation. Hormone-related vascular genes. Auxin: *ARF1*, jasmonic acid: *OPR3*, gibberellin: *CPS1*, cytokinin: *CKX*3, *CKX5*, *CKX11*. W3.5.

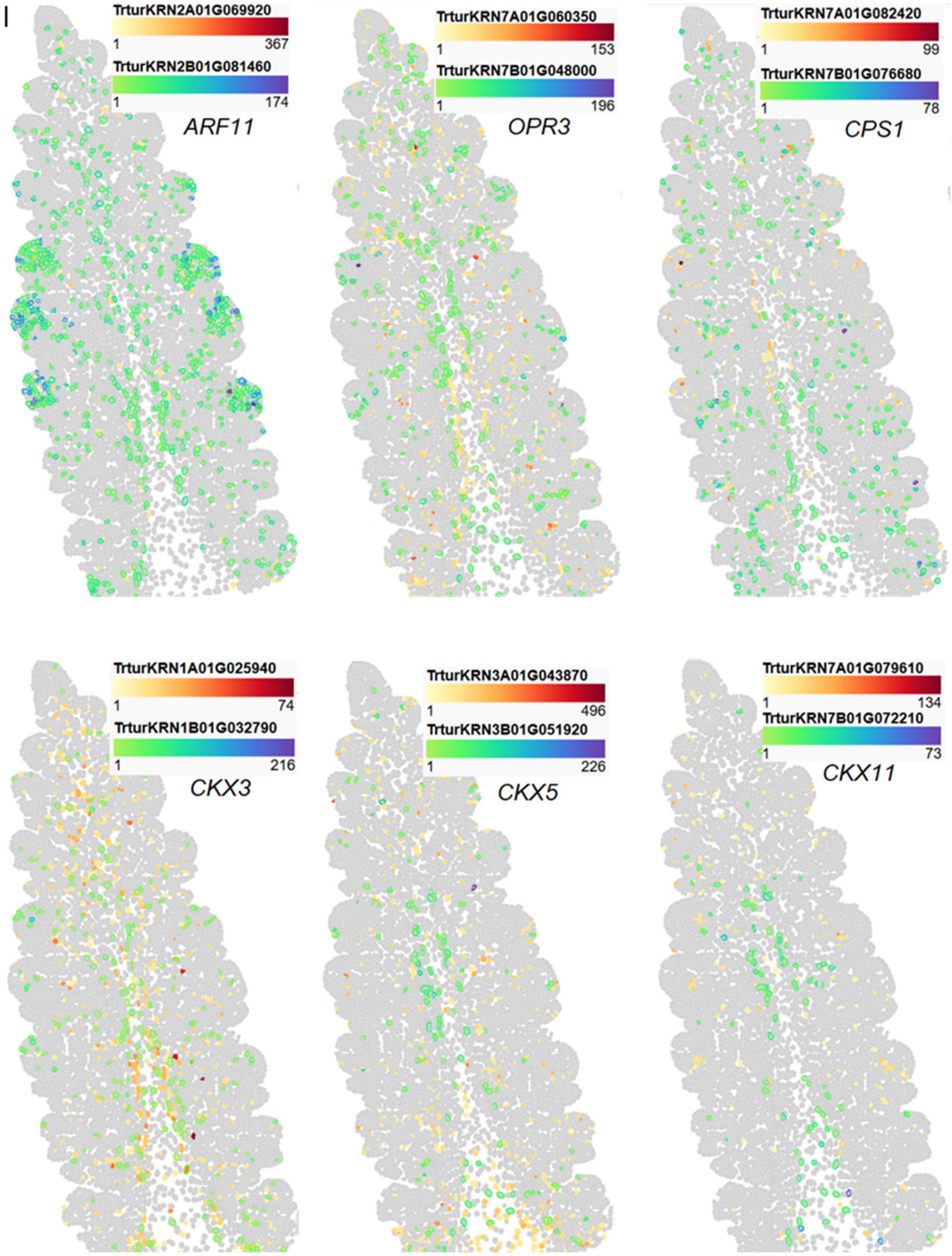

**Fig. S29. Co-expression of *APL* phloem marker with B-class MADS-box genes *PI1* and *AP3*. A-B** UMAPS showing expression of *AP3* and *PI1* in vasculature cluster sc18. **C-E** Co-localization of *APL, AP3* and *PI1* in the vascular tissue of the leaves at W1.5 **C** and W2.5 **D**, and at the center of the spike at W3.5 **E**. Cell walls in E are stained with calcofluor-white.

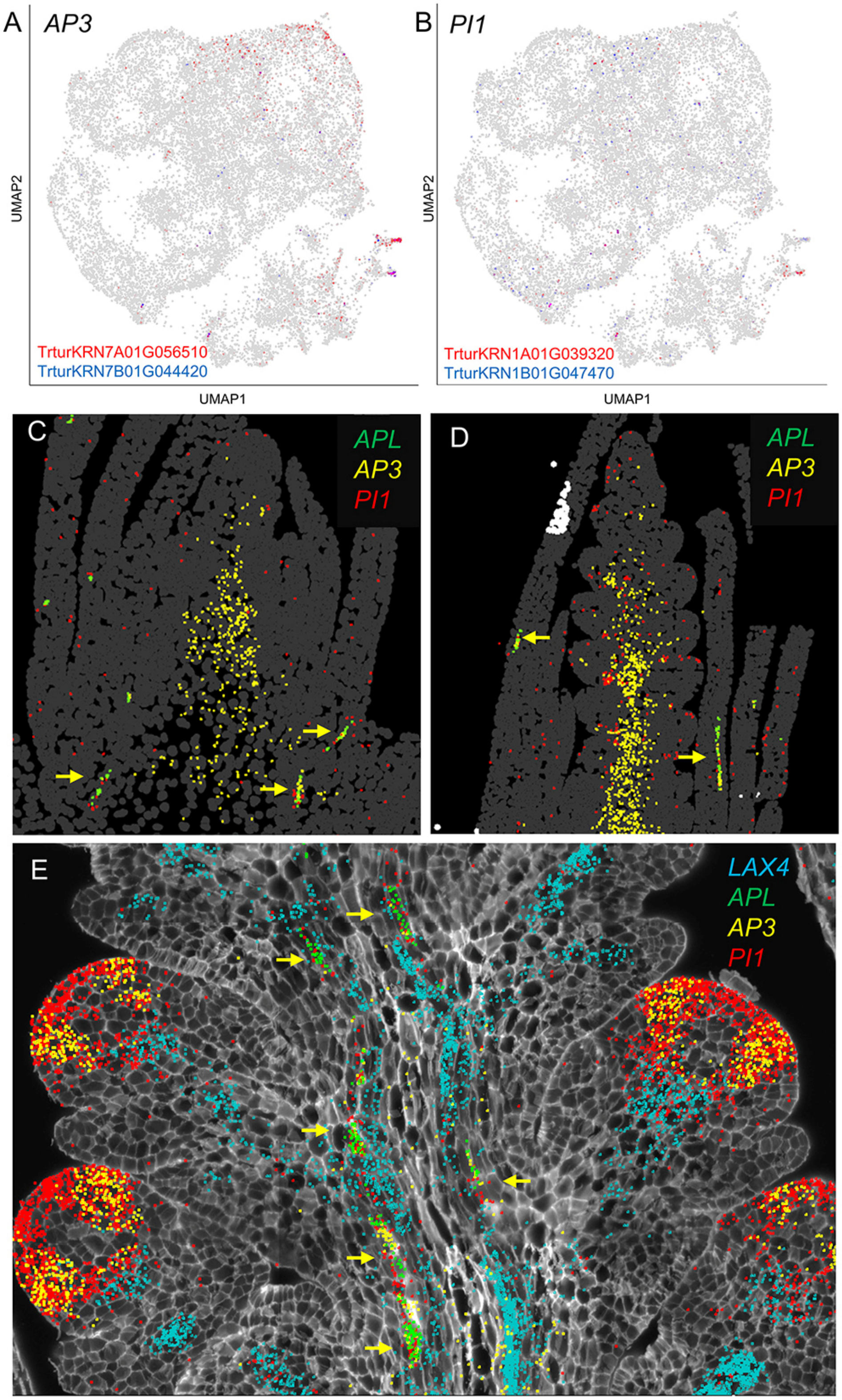

**Fig. S30. Central spike photosynthetic ground tissue. A-F** UMAPS and **G** bubble plots of known central spike ground tissue (scRNA-seq cluster sc6). **H** Imputed expression of central spikelet ground tissue genes *CAP2R* and *LHCB1.1* genes expressed in smFISH cluster c3 (brown). Only one homeolog is shown using the green-blue scale for the cell-border color. Annotation of the marker genes and references are available in Data S1. Additional genes preferentially expressed in cluster c6 are provided in Data S9.

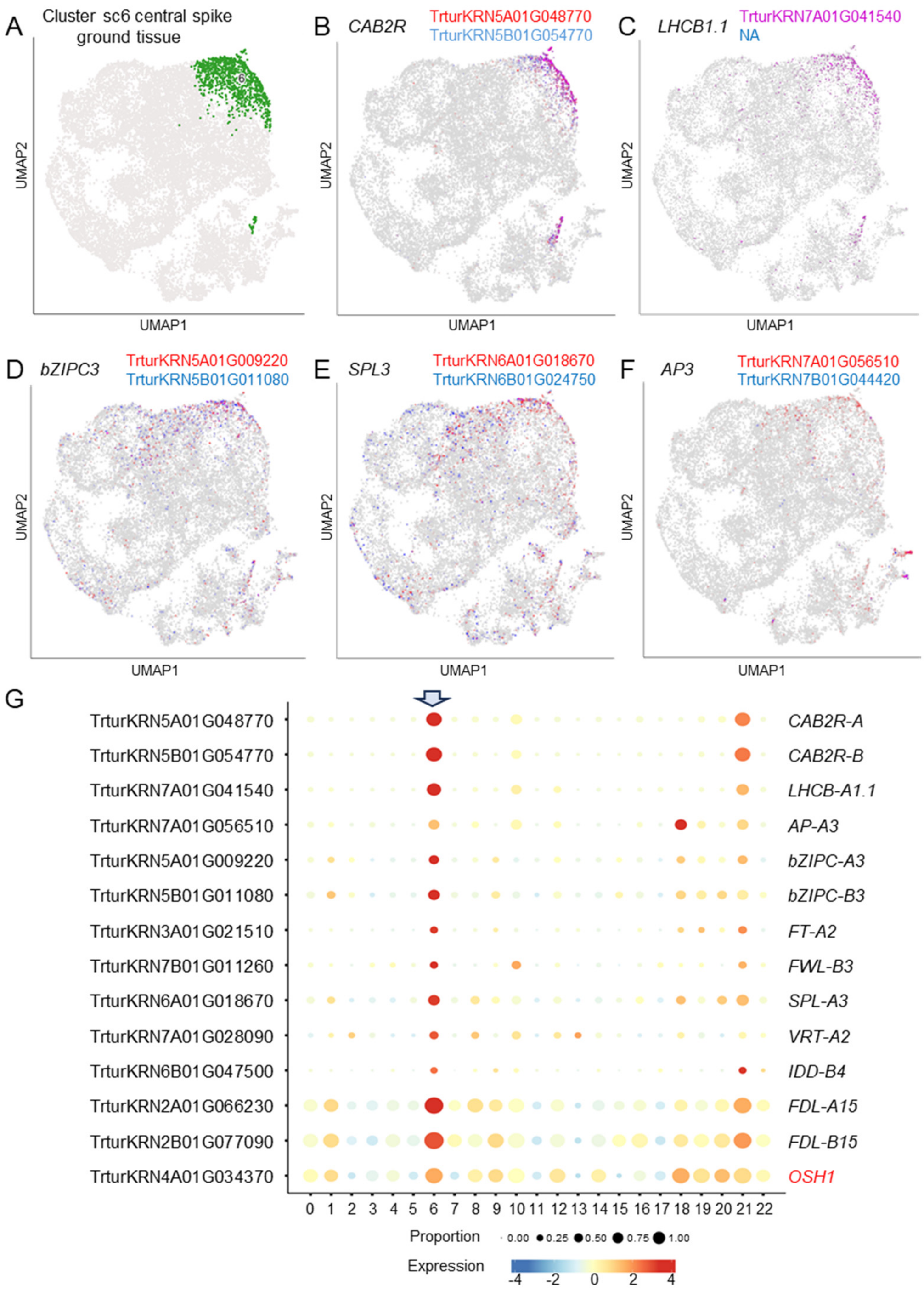

**Fig. S30. Continuation.**

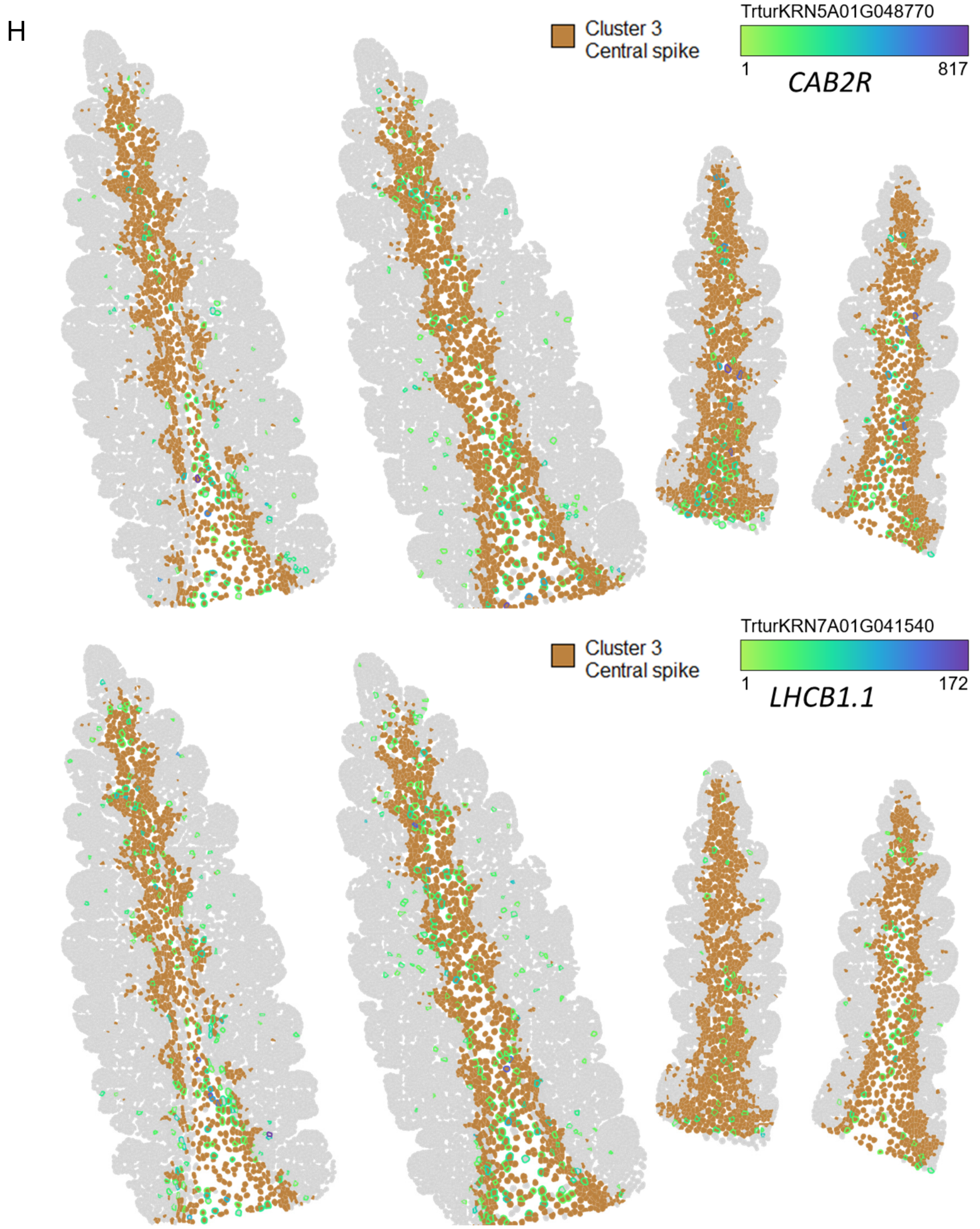

**Fig. S31. Heat map for genes with photosynthesis related annotations.** These genes were selected from the highest preferentially expressed genes in sc6 with photosynthesis related annotations to test the presence of these genes in other clusters. Cluster sc21 was enriched in both cell cycle and photosynthetic markers.

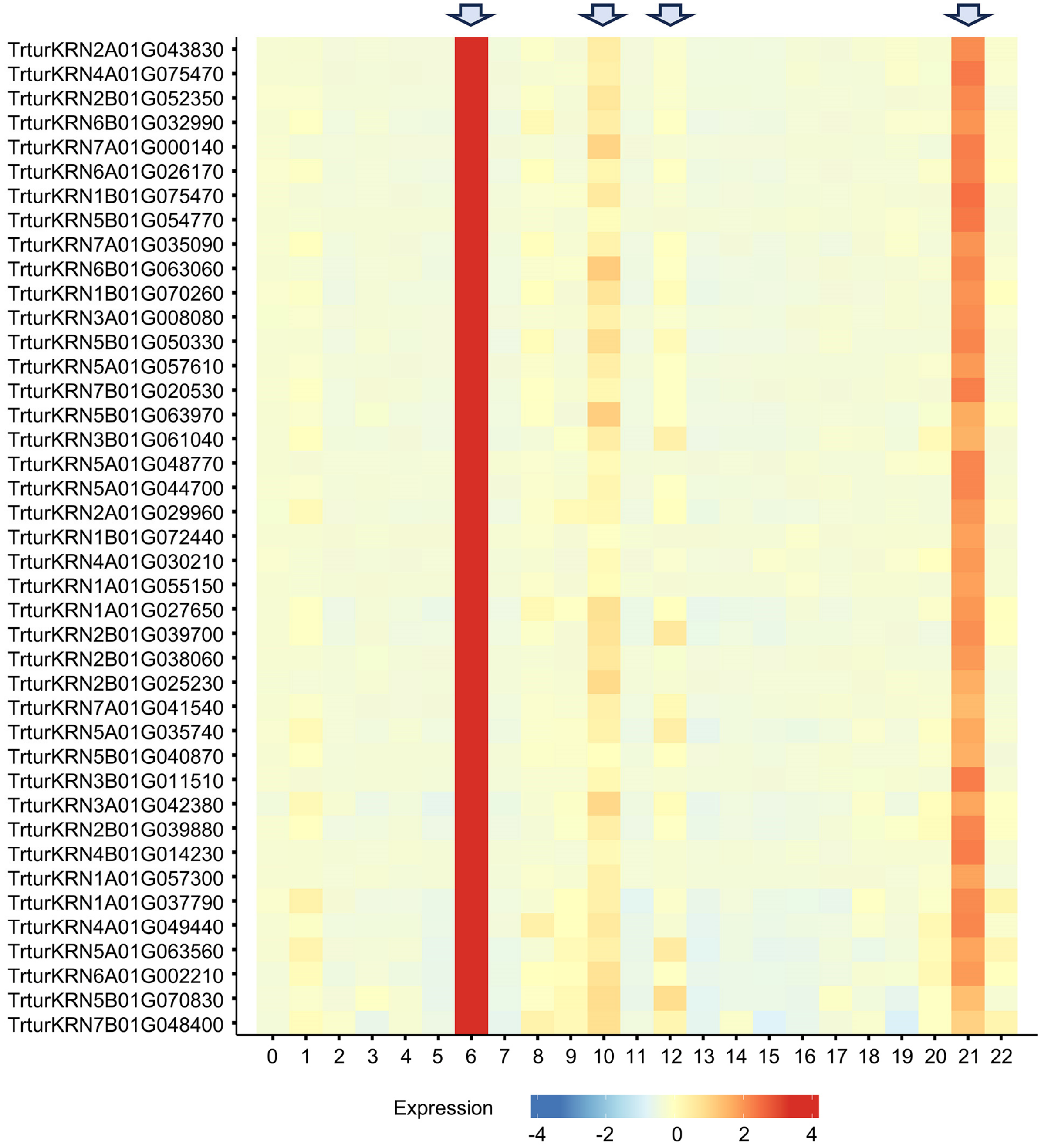

**Fig. S32. Ground tissue. A-F** UMAPS and **G** bubble plots of ground tissue marker genes (sc10 and sc12). Annotation of the marker genes and references are available in Data S1. Additional genes preferentially expressed in ground tissue clusters sc10 and sc12 are provided in Data S9.

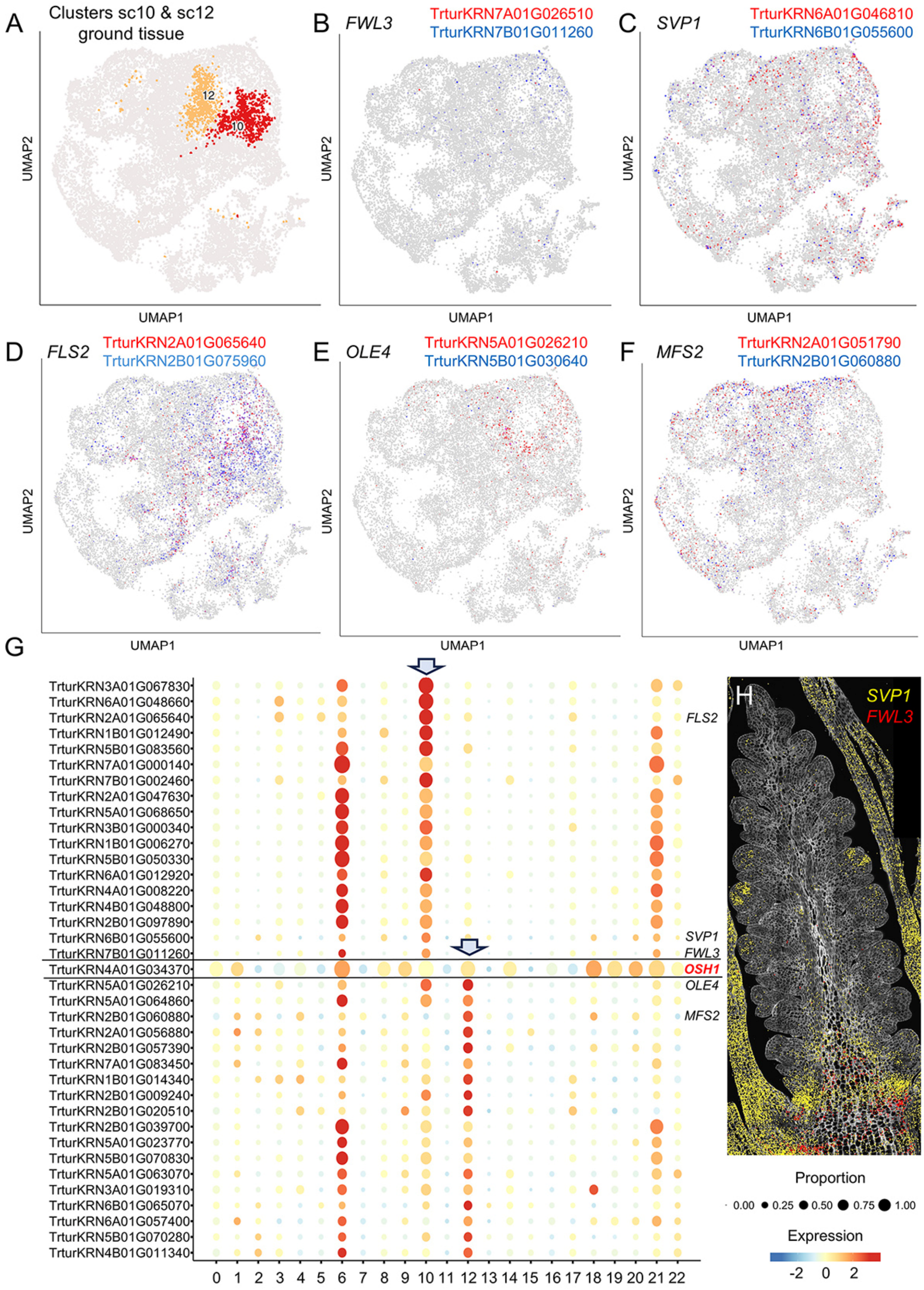

**Fig. S33. Transition leaf-spike. A** smFISH for genes enriched in the transition between leaves and spike at W2.5 (left) and W3.5 (right). **B-D** UMAPS and **E** bubble plots of marker genes for the transition region between the leaves and the spike (sc8). **F** Imputed expression of transition zone genes *LEC1* and *TRD1* at W3.5, JA-related genes *MYC2* and *JAZ8*, and reference genes *TB1* and *SPL17* (smFISH cluster c19). Annotation of the marker genes and references are available in Data S1. Additional genes preferentially expressed in cluster sc8 are provided in Data S9.

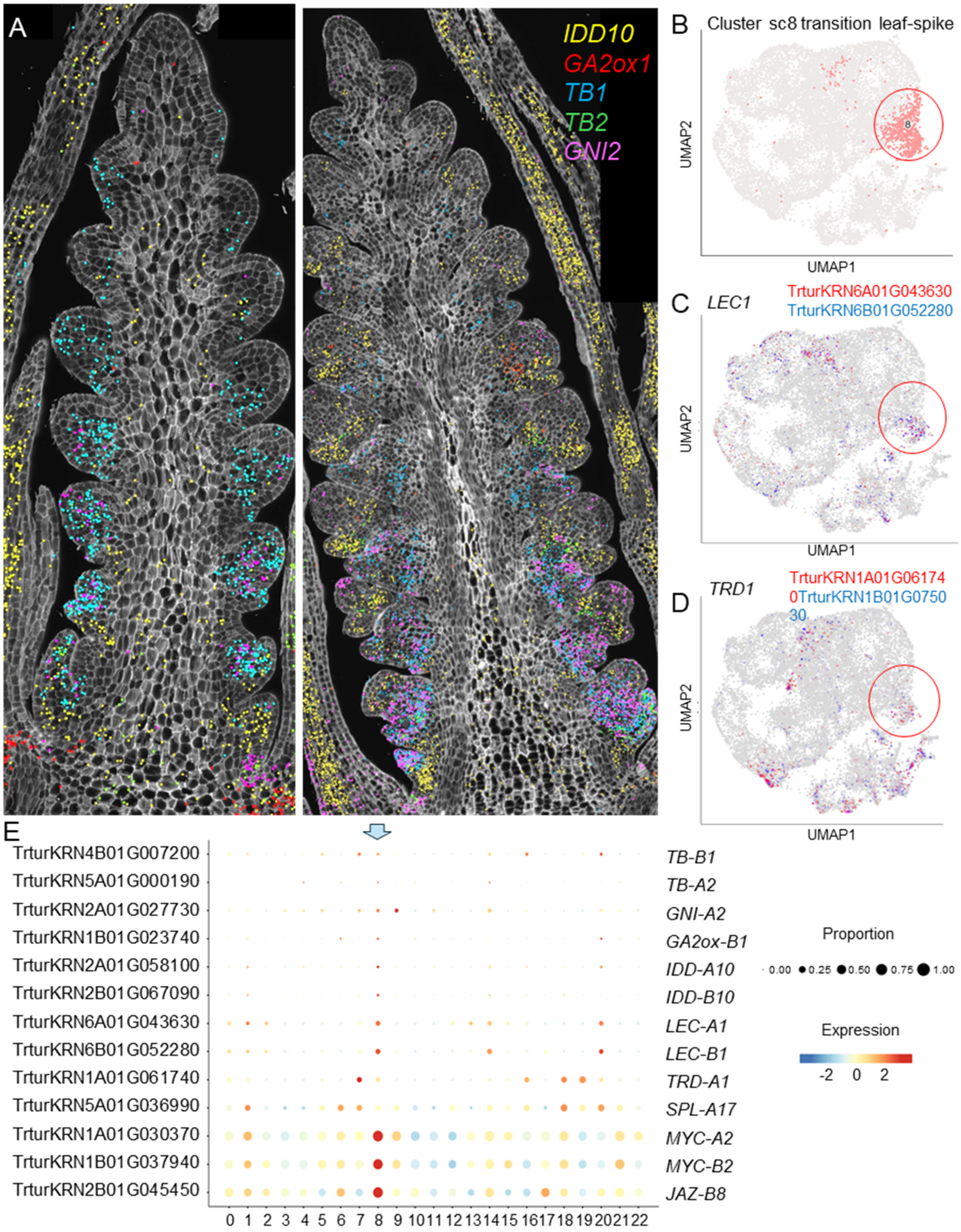

**Fig. S33. Continuation**.

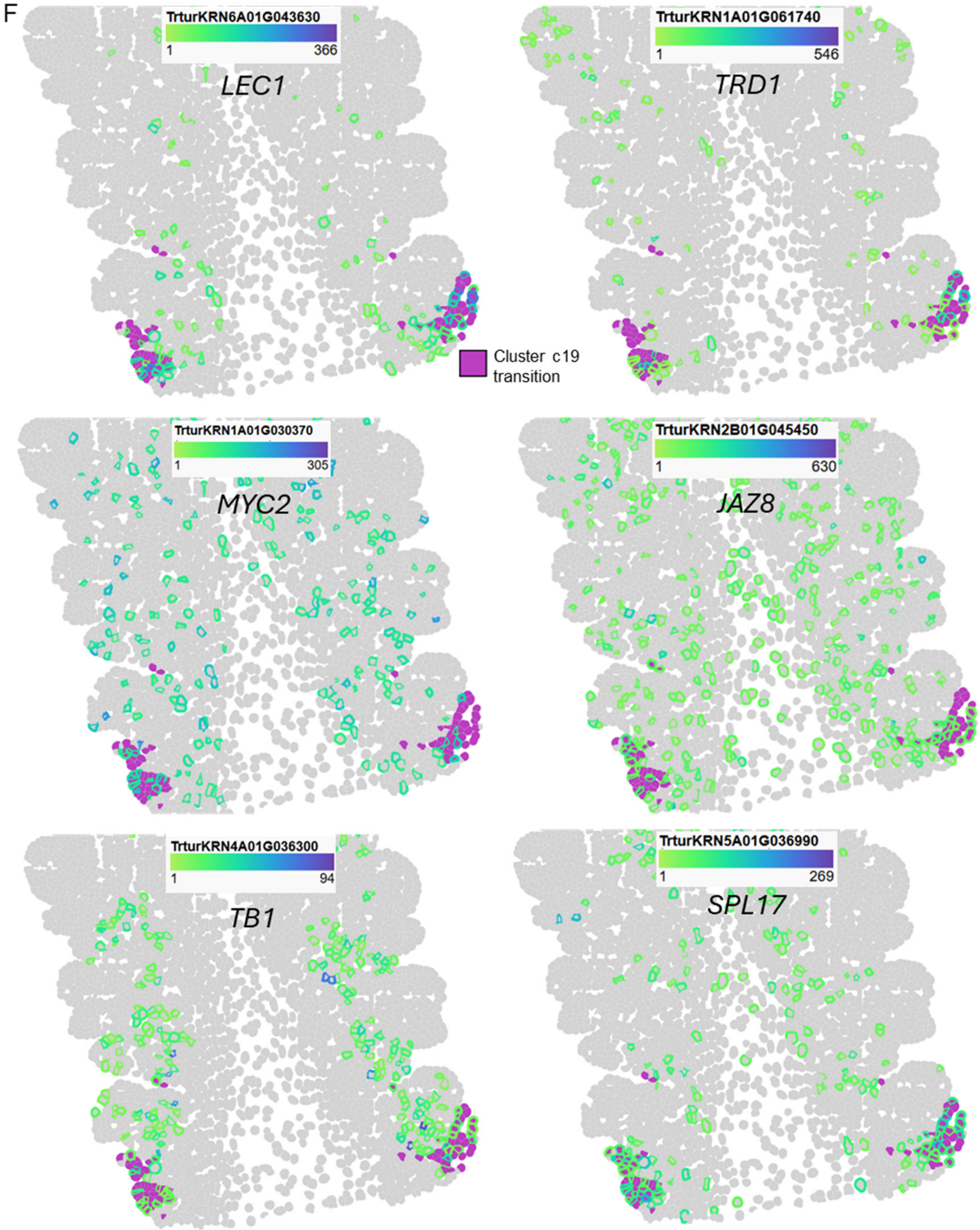

**Fig. S34. Suppressed bract. A-F** UMAPS and **G** bubble plots of marker genes for the suppressed bract cluster sc1. Annotation of the marker genes and references are available in Data S1. Additional genes preferentially expressed in cluster sc1 are provided in Data S9.

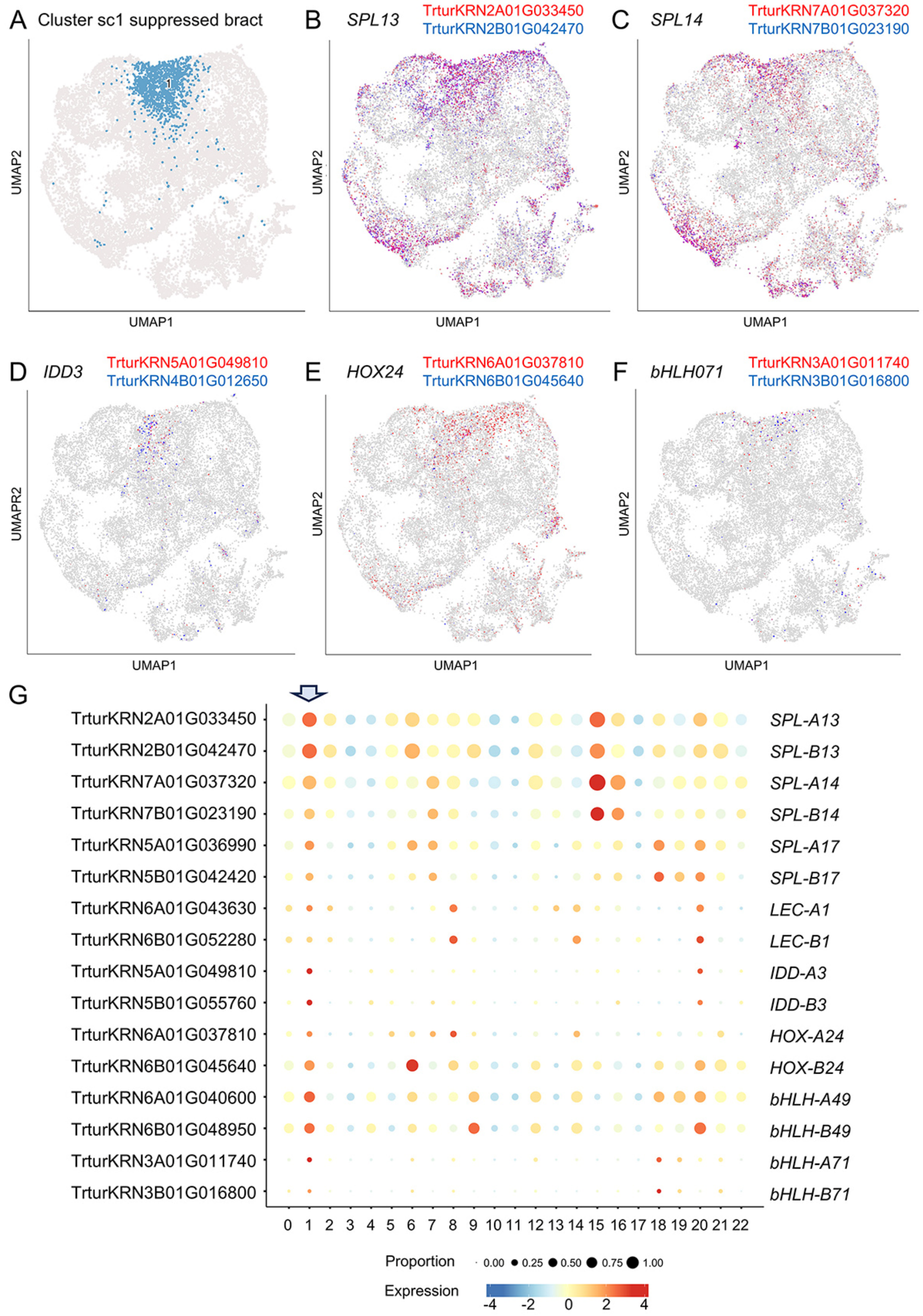

**Fig. S35. Validation of co-expression results using imputed expression and smFISH data. A-B** Imputed expression of boundary genes in developing spike at W3.5 showing clusters c14and c16. Only one homeolog is shown using the green-blue scale for the cell-border color. **A** *WOX9c.* **B** *MFS1.* **C** smFISH expression of *MFS1*, *FZP*, and *TCP24* in spikelet at W3.5. **D** UMAP expression of *MFS1* in single-cell clusters. **E** *CLE33* imputed expression at W3.5 (W2.5 in Fig, S27K). **F** *RA2* imputed expression.

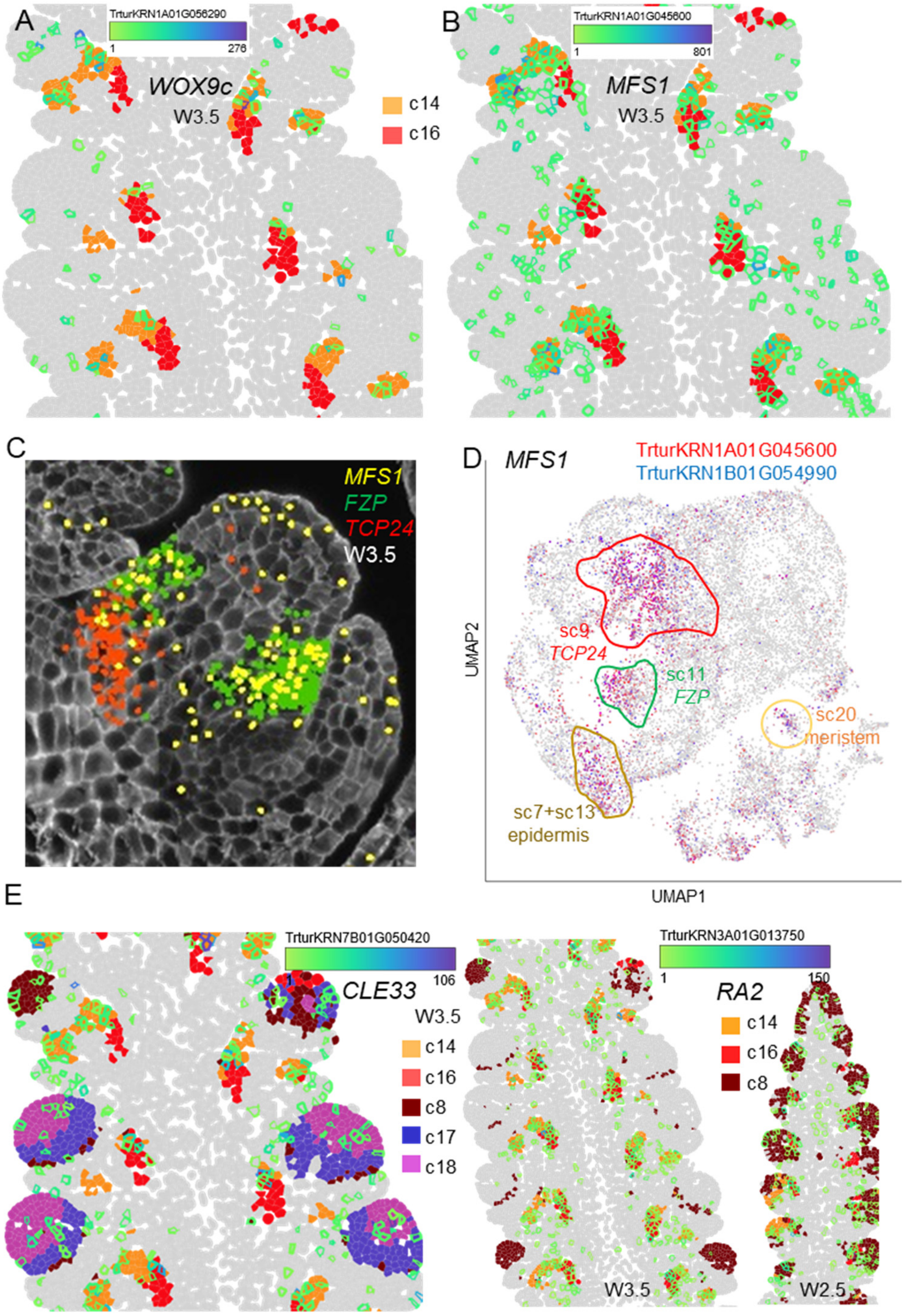

**Fig. S36. Cluster expressing *FZP* and co-expressed genes. A-F** UMAPS and **G** bubble plots for *FZP* and co-expressed genes in the glume axilla cluster sc11. Annotation of the marker genes and references are available in Data S1. Additional genes preferentially expressed in sc11 are available in Data S9.

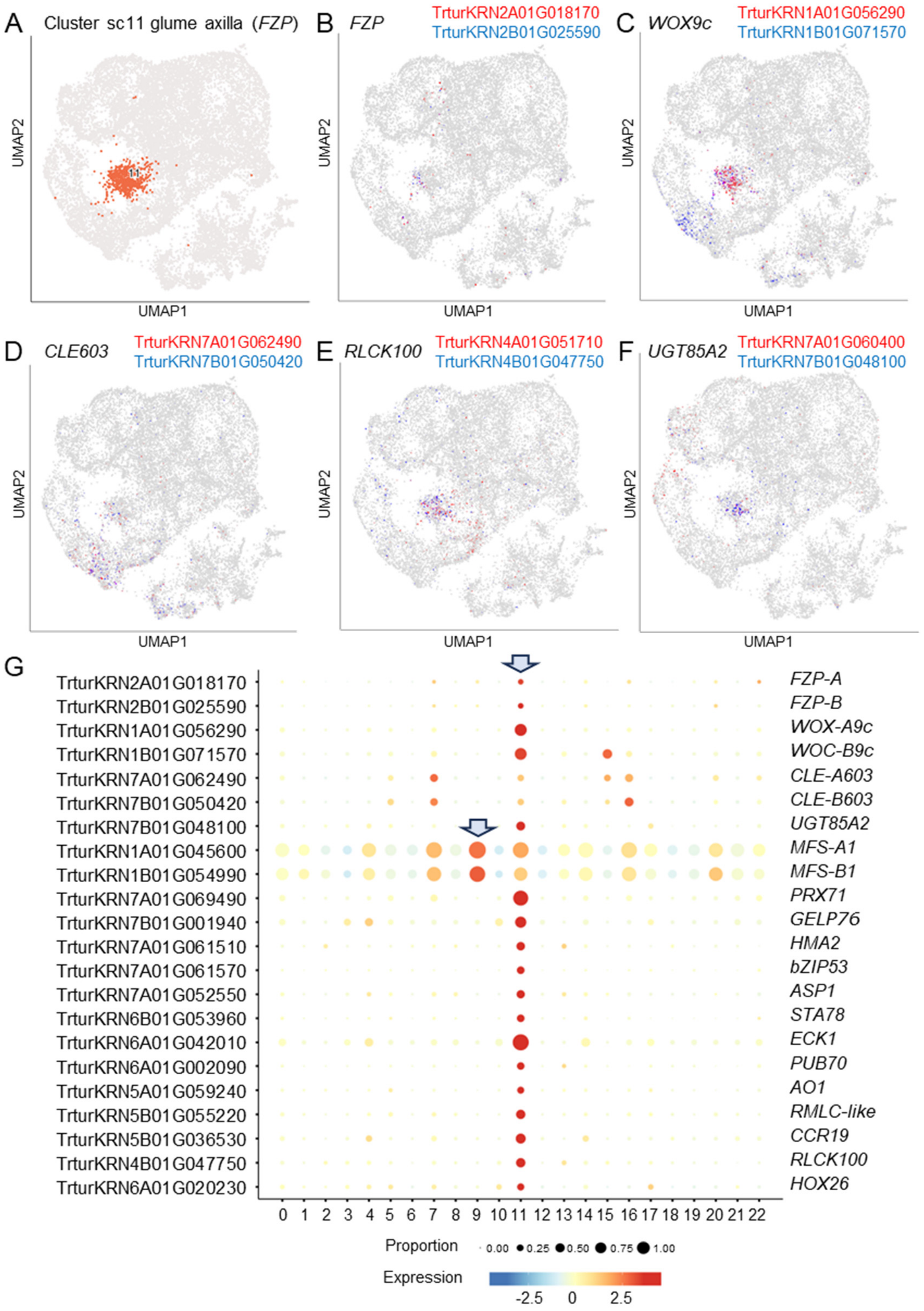

**Fig. S37. Clusters expressing *TCP24* and co-expressed genes. A-F** UMAPS and **G** bubble plots for *TCP24* gene and co-expressed genes in the adaxial boundary clusters sc9 and sc13. Cluster sc13 also expressed epidermal markers. Annotation of the marker genes and references are available in Data S1. Additional genes preferentially expressed in sc9 and sc13 are available in Data S9.

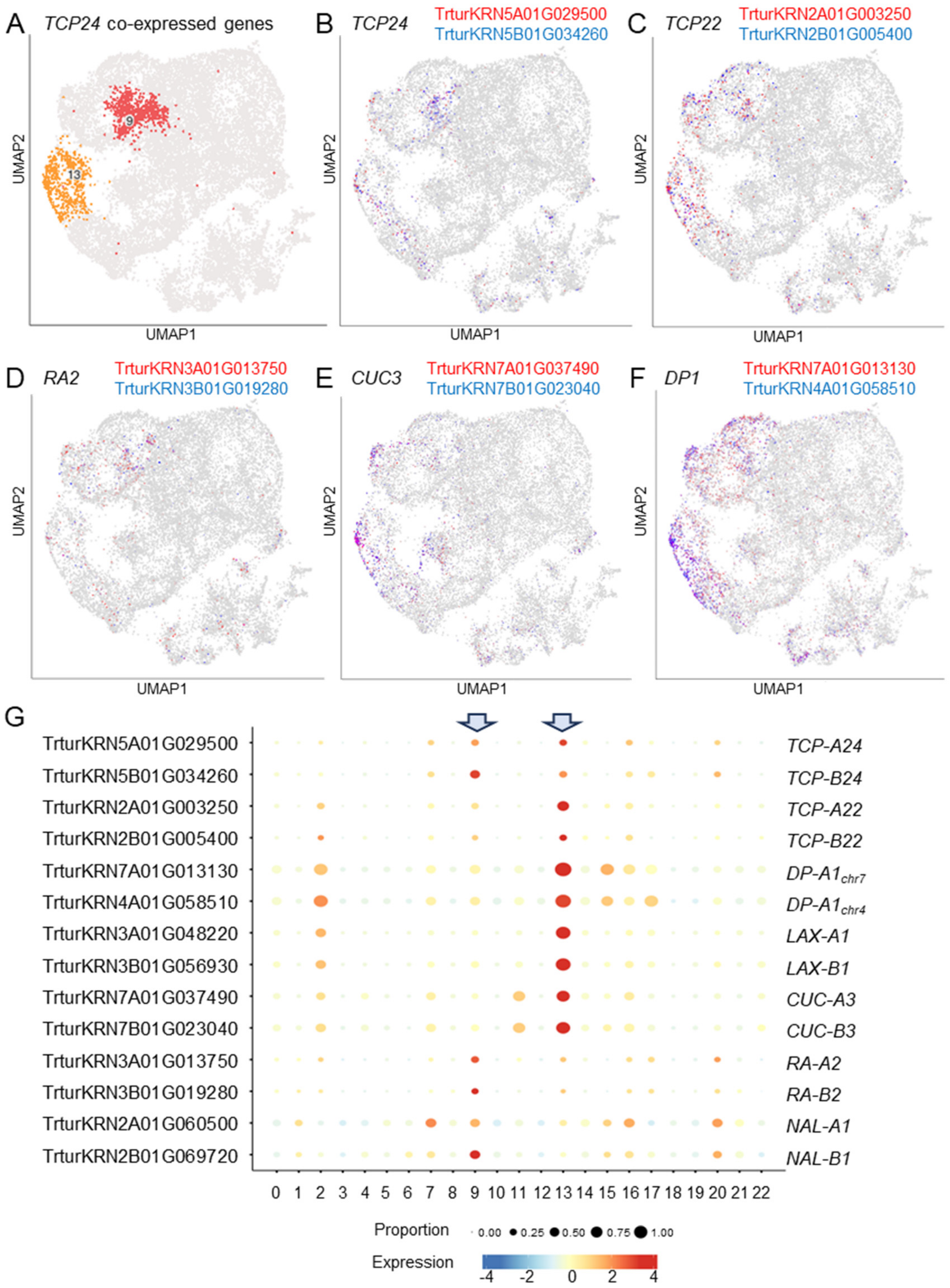

**Fig. S38. Cortex region below developing spikelets. A-F** UMAPS and **G** bubble plots of marker genes for the cortex region below the spike (cluster sc2). **H** Spatial pattern of cortex genes using imputed expression. A-genome homeologs: fill color using yellow-red scale, B-genome: border color using green-blue scale. Annotation of marker genes and references are available in Data S1. Additional genes preferentially expressed in cluster sc2 are provided in Data S9. Many of the preferentially expressed genes in sc2 and are also preferentially expressed in epidermal cluster sc13. Both clusters are highly correlated (*R* =0.96, Data S8).

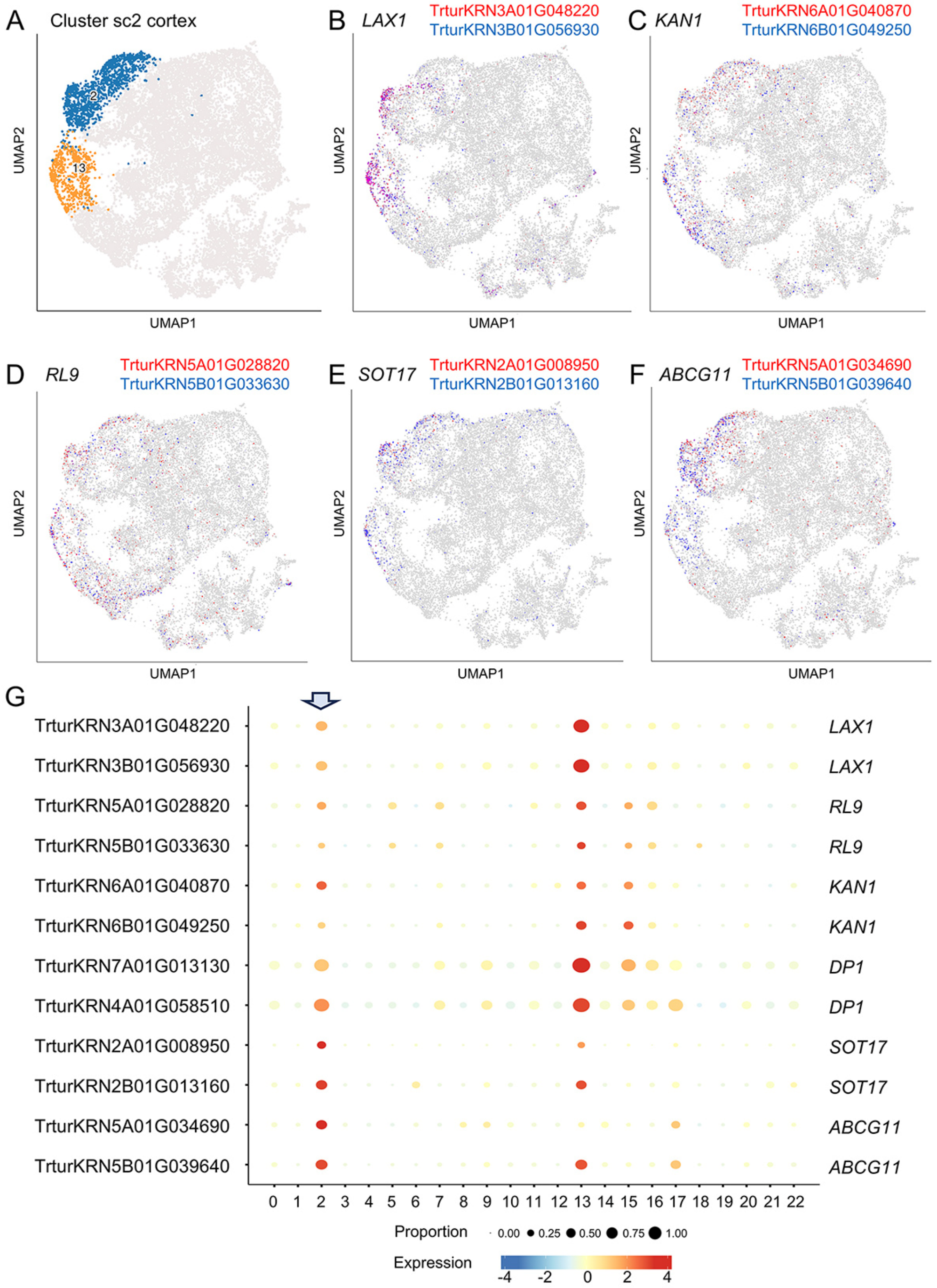

**Fig. S38. Continuation**.

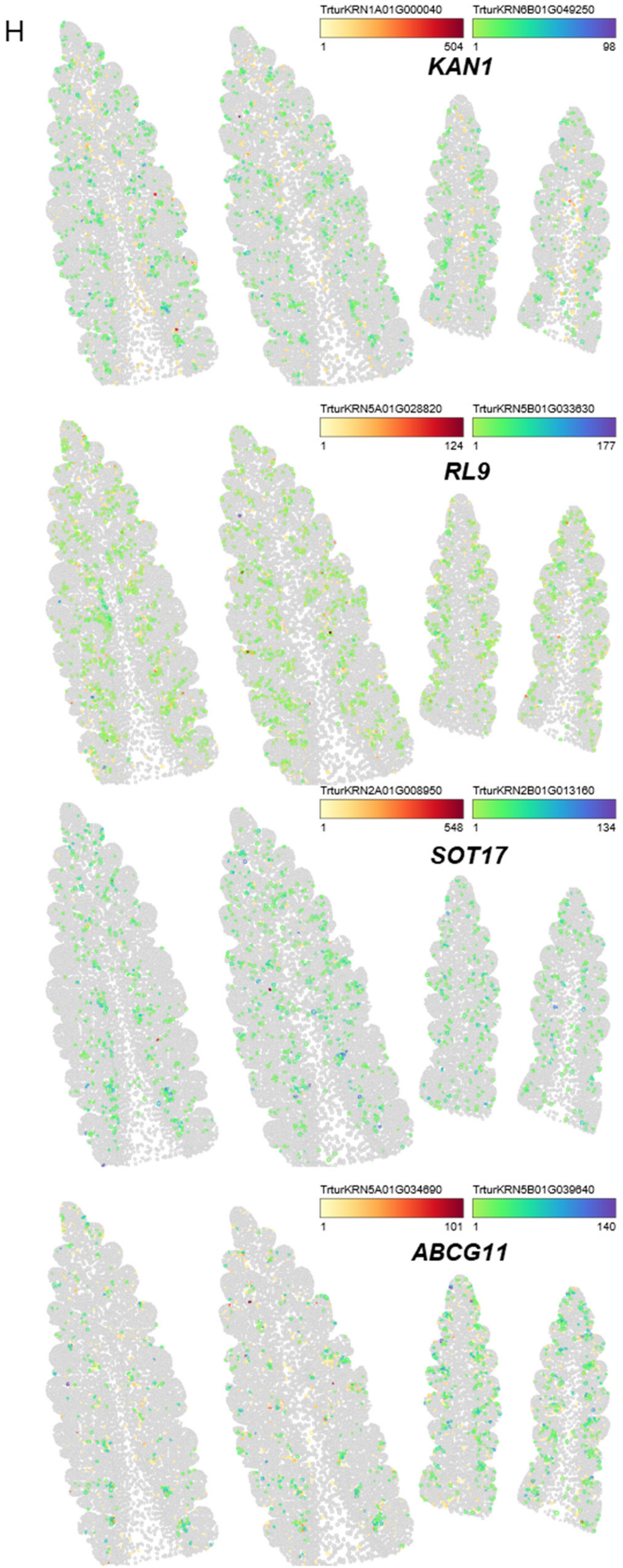

**Fig. S39. Cortex region below developing spikelets.** UMAPS and bubble plots of marker genes for the cortex region below the spike (cluster sc4 & sc17). **A** Clusters sc4 and sc17. **B-D** Examples for sc17. **E-F** Examples for sc4. **G** Bubble plot. Annotation of marker genes and references are listed in Data S1. Additional genes preferentially expressed in clusters sc4 and sc17 are available in Data S9.

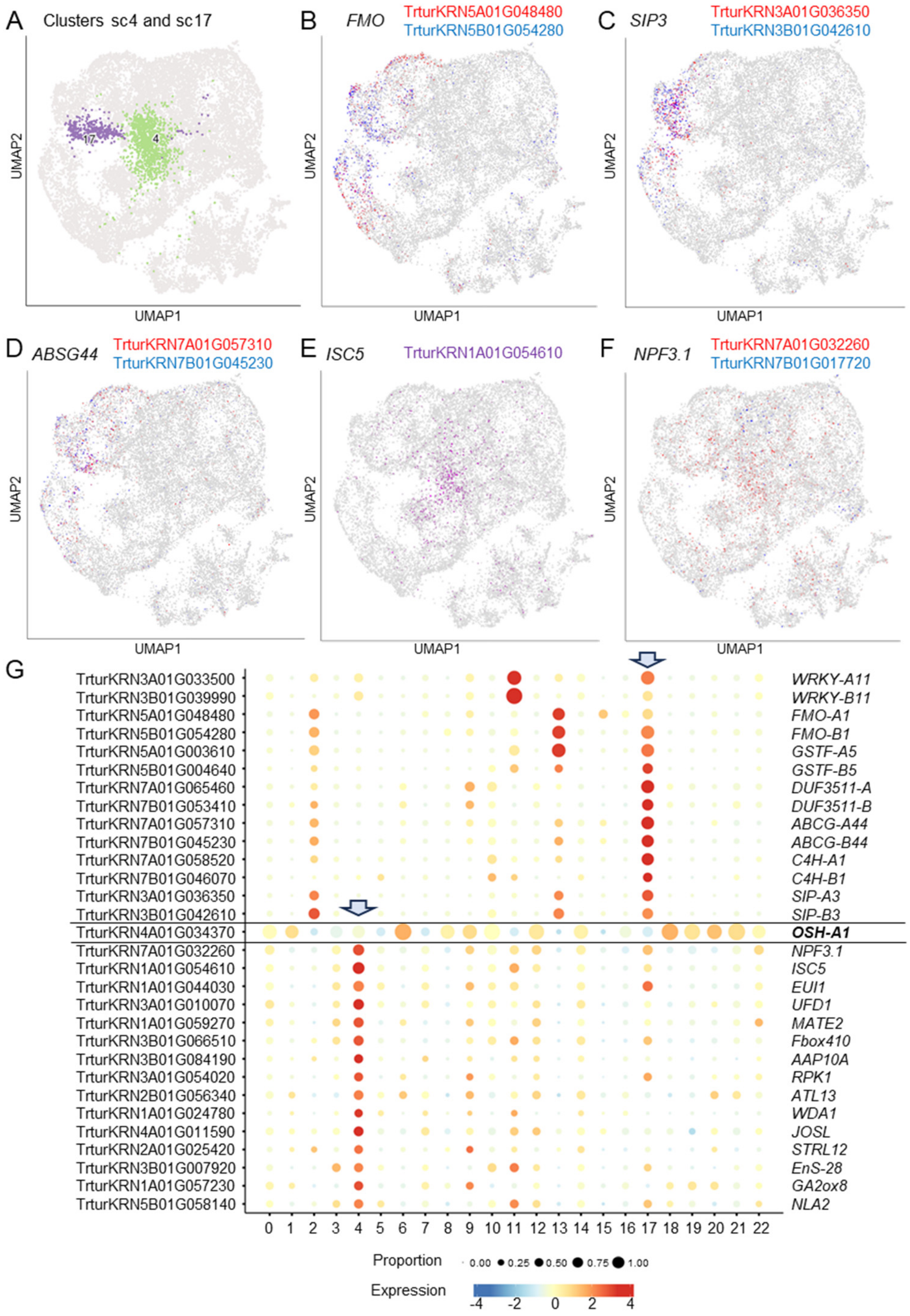

**Fig. S40. Meristem. A-F** UMAPS and **G** bubble plots of meristematic marker genes expressed in sc14 and sc20. **H-J** Differential expression profiles of *ULTRAPETALA1* homeologs *ULT-A1* and *ULT-B1*. **H-I** UMAPS. **J-K** Imputed expression of both homeologs. **L-P** Imputed expression of additional genes expressed in meristematic clusters sc14 and sc20. In **J to P** levels of gene expression are indicated by a green-blue color expression scale for the cell-borders. Additional genes preferentially expressed in sc14 and sc20 are provided in Data S9. Annotation of the marker genes and references are available in Data S1.

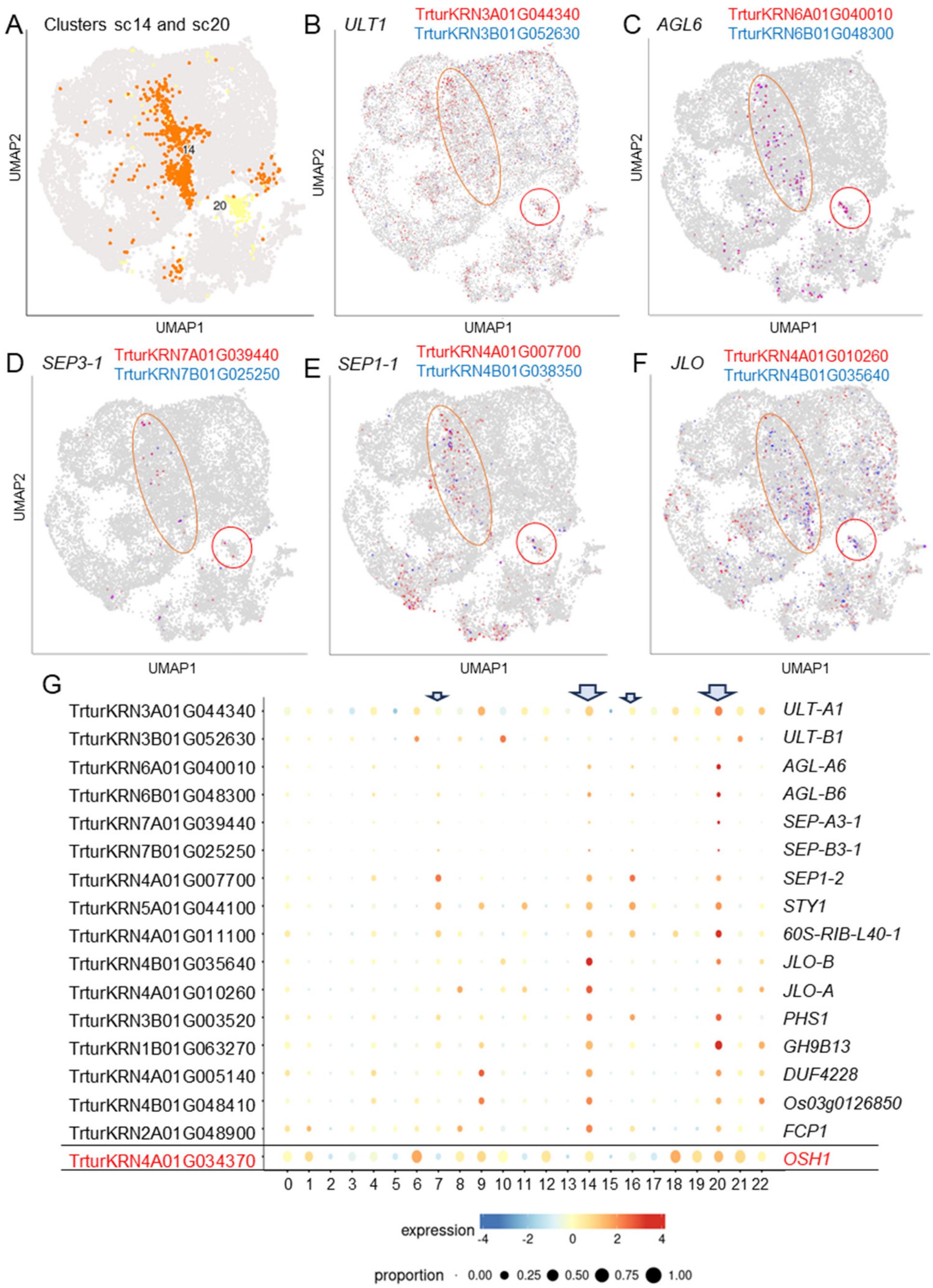

**S40. Continuation**. Differential expression of *ULT-A1* and *ULT-B1*

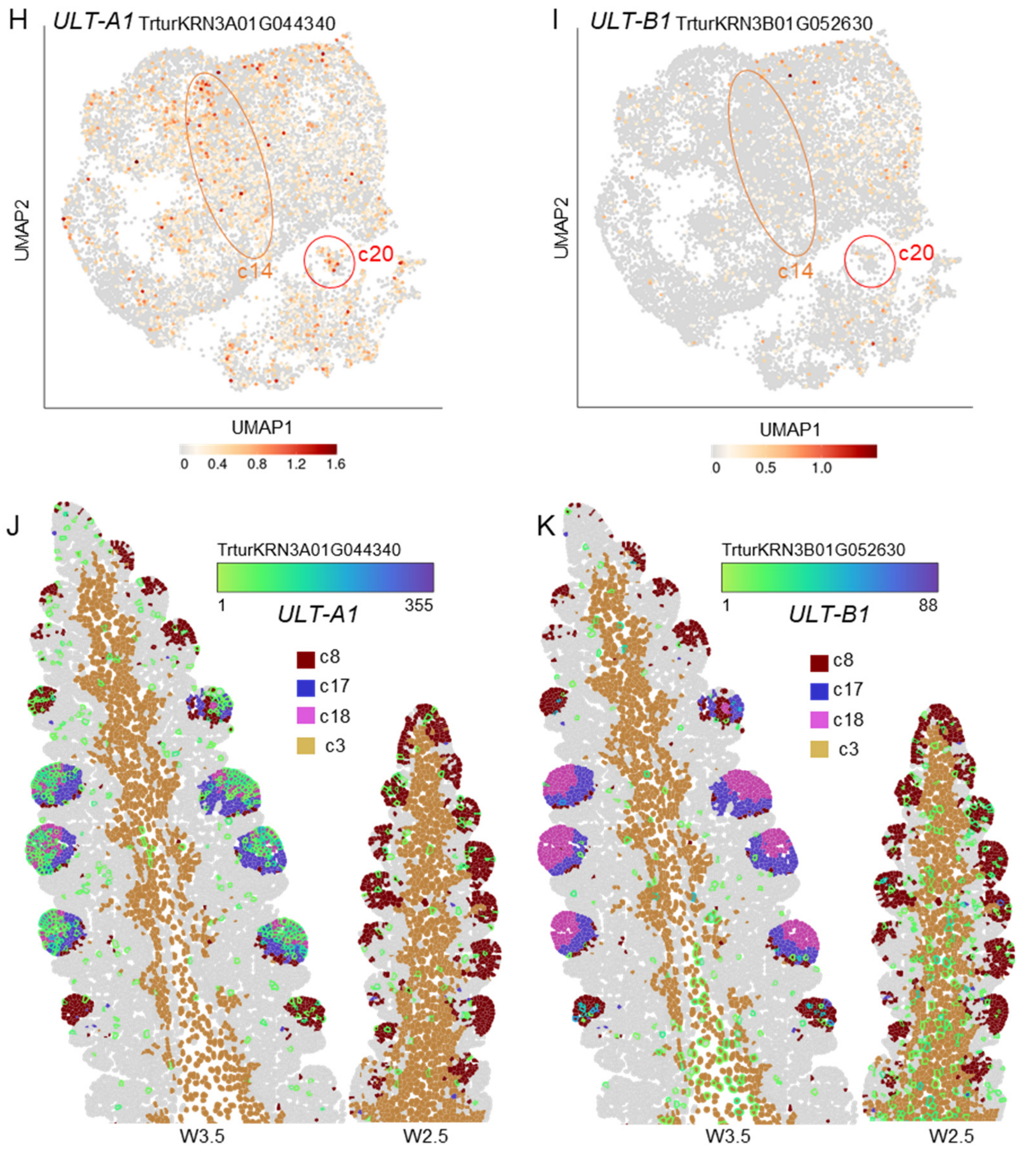

**S40. Continuation**. Genes preferentially expressed in meristematic clusters sc20 and sc14.

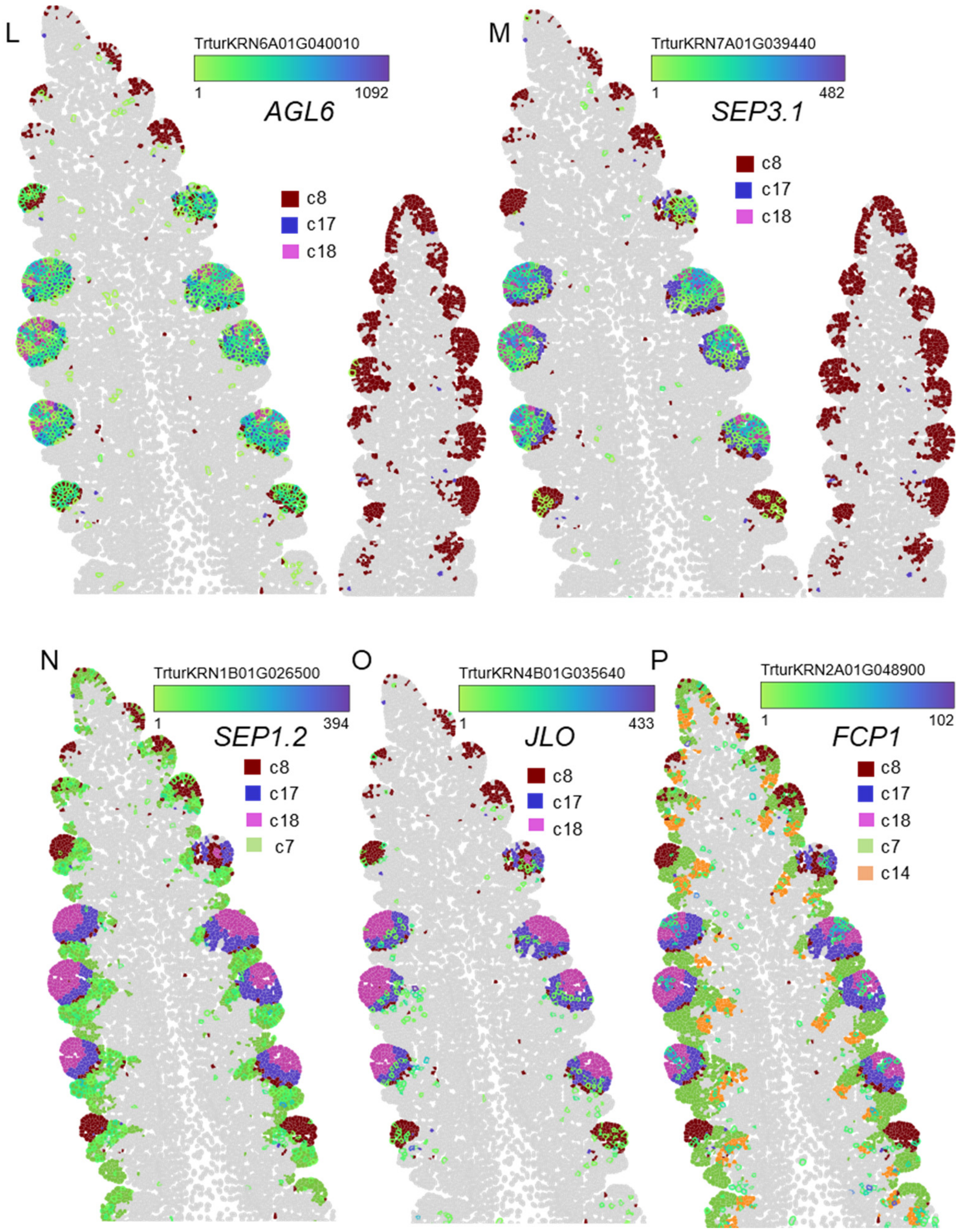

**Fig. S41. Genes co-expressed with *AGL-6*.** Imputed expression of genes selected based on co-expression with *AGL6* and preferential expression in meristems. Only one homeolog is shown using the green-blue scale for the cell-border color. **A** *AG1* is included as control to show correct imputation of c18 location. **B** *DDM1a*, **C** *BRBN16*, and **D** *SHI1*. Annotation of the marker genes and references are available in Data S1.

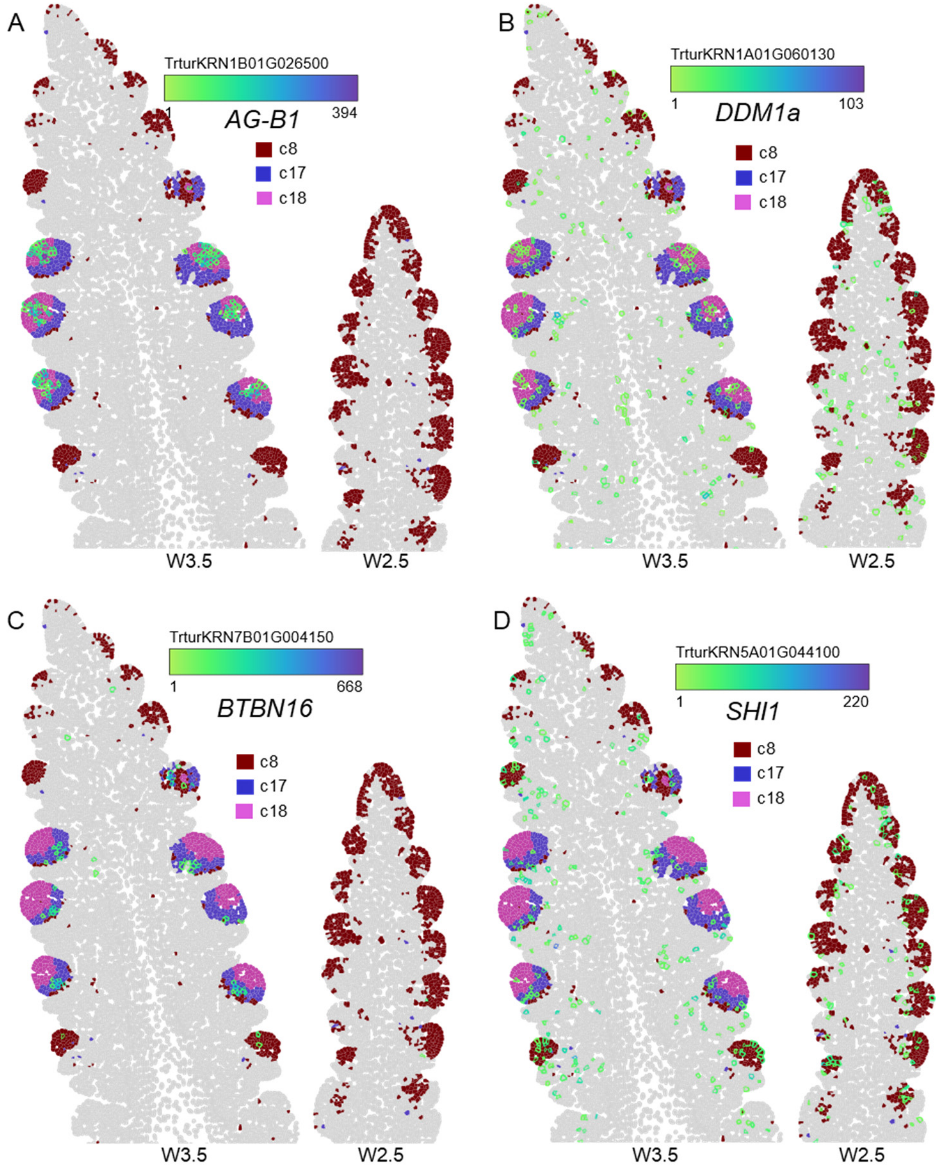

**Fig. S42. Gene network for *FZP* and *TCP24*.** Gene regulatory network constructed with the program GENIE3. The network is based on the 988 genes co-expressed with *FZP* (Data S20) and/or *TCP24* (Data S21) in the scRNA-seq dataset. A summary of the gene interactions is presented in Data S22. The network presented below includes the highly significant interactions visualized with Cytoscape 3.10.3 ([https://cytoscape.org/](https://cytoscape.org/)).

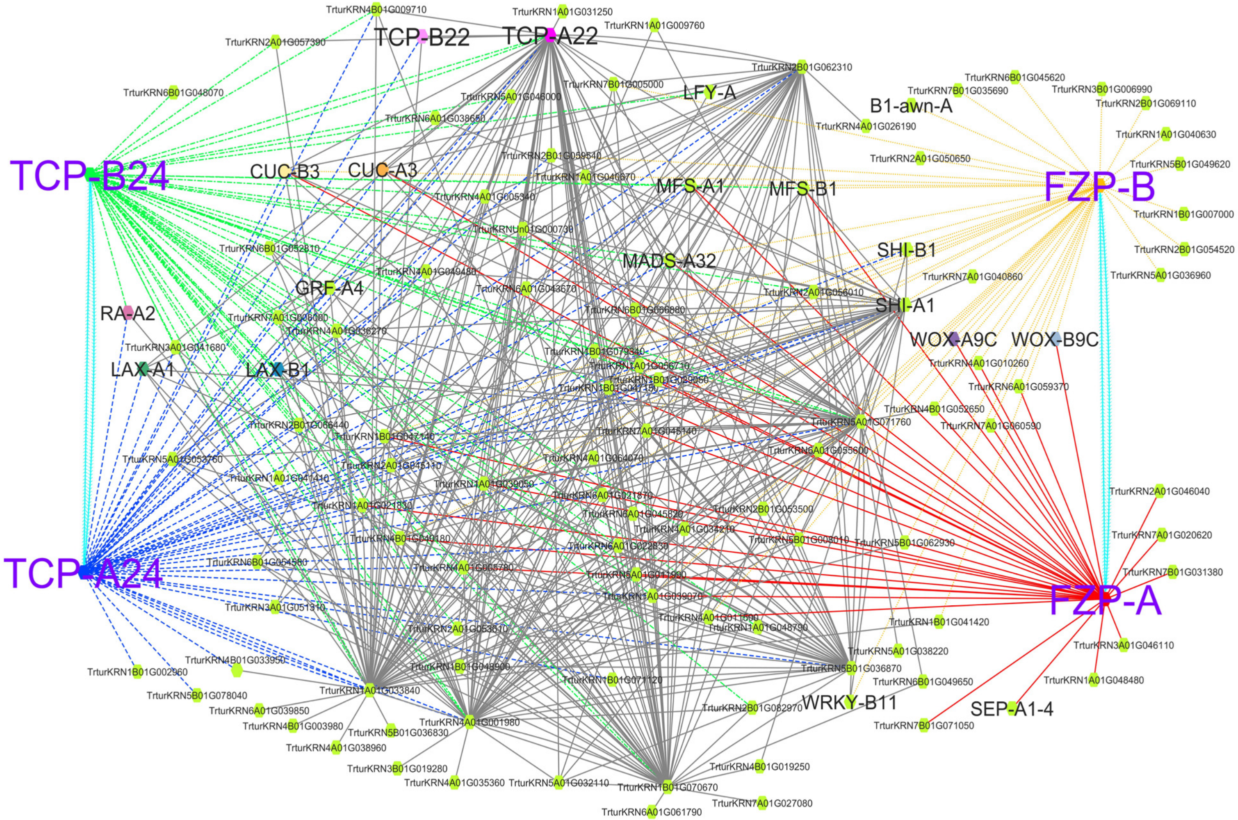

**Fig. S43.** Heatmap for genes encoding ribosomal proteins co-expressed with *ULT-A1* (82 out of the 100 genes with highest correlations with *ULT-A1*). The *ULT-B1* homeolog showed limited co-expression with *ULT-A1* (*R*= 0.0398) and no enrichment in genes encoding ribosomal proteins suggesting functional differentiation between homeologs.

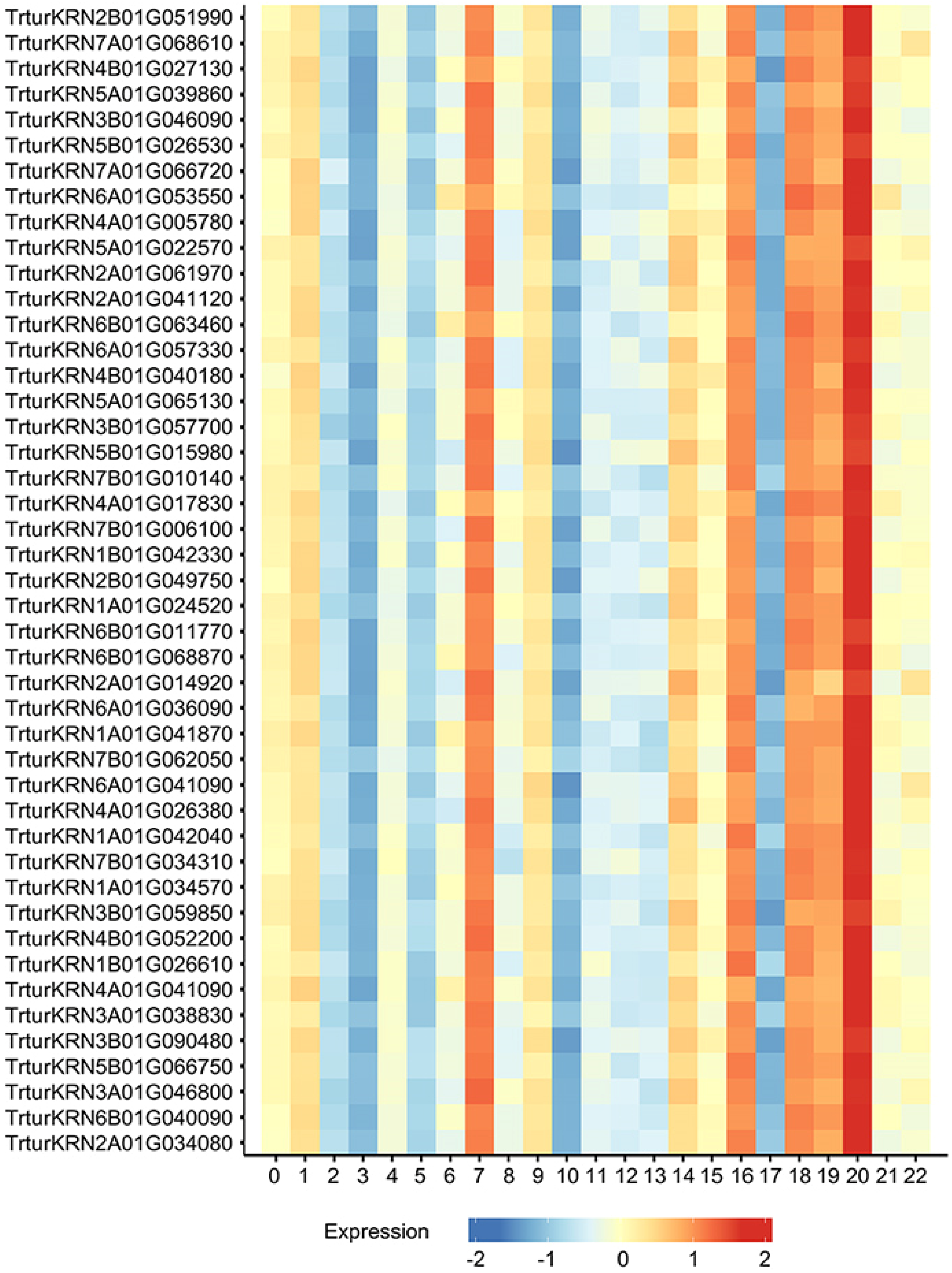

**Fig. S44. Trajectory analysis scRNA-seq clusters.** The 23 cell clusters from the scRNA-seq analysis were used for a trajectory analysis generated with program Monacle 3. The cluster numbers and colors are the same as in Fig. 5. Cluster sc14, which was annotated as meristematic cells was used as root.

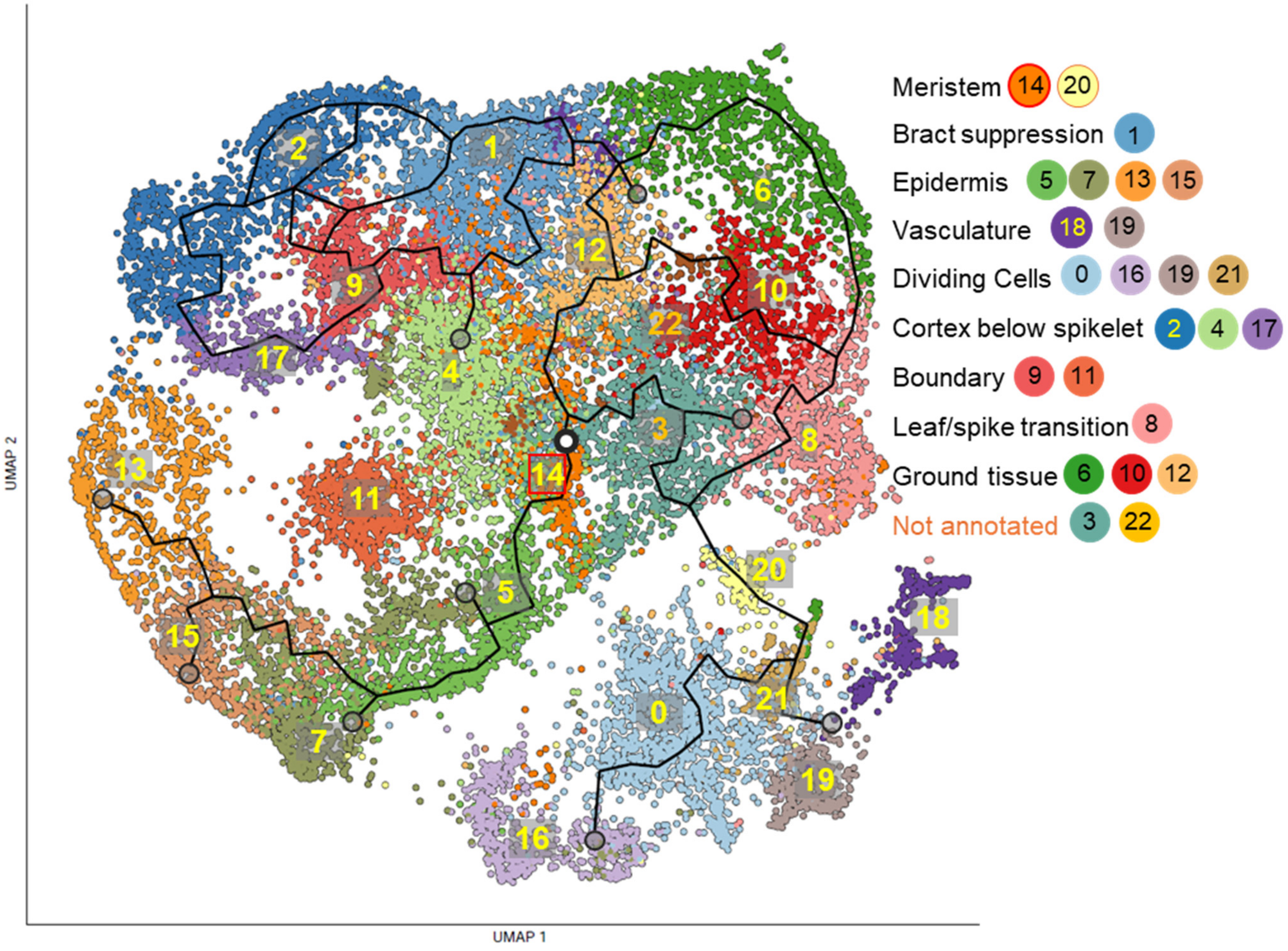
